# Performance comparison of systemic activity correction in functional near-infrared spectroscopy for methods with and without short distance channels

**DOI:** 10.1101/2022.03.31.486522

**Authors:** Franziska Klein, Michael Lührs, Amaia Benitez-Andonegui, Pauline Roehn, Cornelia Kranczioch

**Author notes:** Address all correspondence to Franziska Klein.

## Abstract

**Significance:** Functional near-infrared spectroscopy (fNIRS) is a promising tool for neurofeedback (NFB) or brain–computer interfaces (BCIs). However, fNIRS signals are typically highly contaminated by systemic activity (SA) artifacts, and, if not properly corrected, NFB or BCIs run the risk of being based on noise instead of brain activity. This risk can likely be reduced by correcting for SA, in particular when short-distance channels (SDCs) are available. Literature comparing correction methods with and without SDCs is still sparse, specifically comparisons considering single trials are lacking.

**Aim:** This study aimed at comparing the performance of SA correction methods with and without SDCs.

**Approach:** Semisimulated and real motor task data of healthy older adults were used. Correction methods without SDCs included a simple and a more advanced spatial filter. Correction methods with SDCs included a regression approach considering only the closest SDC and two GLM-based methods, one including all eight SDCs and one using only two *a priori* selected SDCs as regressors. All methods were compared with data uncorrected for SA and correction performance was assessed with quality measures quantifying signal improvement and spatial specificity at single trial level.

**Results:** All correction methods were found to improve signal quality and enhance spatial specificity as compared with the uncorrected data. Methods with SDCs usually outperformed methods without SDCs. Correction methods without SDCs tended to overcorrect the data. However, the exact pattern of results and the degree of differences observable between correction methods varied between semisimulated and real data, and also between quality measures.

**Conclusions:** Overall, results confirmed that both Δ[HbO] and Δ[HbR] are affected by SA and that correction methods with SDCs outperform methods without SDCs. Nonetheless, improvements in signal quality can also be achieved without SDCs and should therefore be given priority over not correcting for SA.

## 1 Introduction

Functional near-infrared spectroscopy (fNIRS) is a promising tool in neuroimaging, in particular in the context of brain–computer interface^1^ (BCI) or neurofeedback^2^ (NFB) applications for motor neurorehabilitation. In active overt^3^ NFB training, participants learn to self-regulate their task-related brain activity by receiving feedback with the main goal of facilitating brain plasticity.^2,4,5^ For the purpose of motor neurorehabilitation the task can be any motor task, however, most commonly, participants perform kinesthetic motor imagery, that is, imagining the sensation of a motor task without actually performing it.^2,6,7^ To achieve this goal, a spatially specific brain imaging tool that allows cost-efficient, repeated training is desirable. FNIRS combines these qualities.^2,8^ Moreover, fNIRS allows for measure difficult populations such as children and patients, it is relatively robust against movement, and, because fNIRS can be mobile and portable, it offers environmental flexibility.^5,8–10^ One major drawback of fNIRS, however, is the contamination of the measured signals with task-evoked systemic extracerebral and cerebral activity^11,12^ (in short: systemic activity, SA). FNIRS captures hemodynamic activity by transmitting NIR light through the brain from a light source to a light detector. On this journey, the light does not only penetrate the cerebral tissue but also the extracerebral layers (scalp and skin), resulting in a signal that includes both cerebral and extracerebral hemodynamic activity.^11,12^ The problems arising from the SA artifact are diverse: the artifact differs between signal types oxygenated (Δ[HbO]) and deoxygenated (Δ[HbR]) hemoglobin,^11–13^ between and within subjects as well as tasks.^14^ Furthermore, it is not homogeneously distributed across the head^12,13^ and it can mimic task-related activity.^12,13^ Since the frequency of the artifacts can overlap with the task frequency, the conventionally used temporal filters are not sufficient. The consequence is that statistical results might either be inflated due to false positives or depleted by false negatives.^11,12,15,16^ For NFB and BCI applications this could mean that they are potentially based on noise and not on brain activity.

So far, the gold standard for systemic artifact correction (SAC) of the extracerebral part of the SA artifact involves short-distance channels (SDCs). For SDCs, NIR light source and detector are placed at a distance of <10 mm^11–13,17^ (ideally at 8.4 mm for adults^18^). Because of the short distance, the SDC measures mostly extracerebral SA, which can then be used to correct the data, e.g., by applying a regression-based approach. ^13,17,19–21^ The most promising approach so far is to add the SDC data as (additional) regressors to the General Linear Model (GLM).^13,17,21^ The optimum number of SDCs within this approach is still unknown, but it has been shown for up to eight SDCs that results keep improving when adding more SDCs.^17^ Also, including SDCs of both signal types together, i.e., Δ[HbO] and Δ[HbR], improves the results.^17^ However, not all fNIRS researchers have access or will have access to SDCs in the near future. For instance, von Lühmann and colleagues^21^ found that only 4% of published fNIRS BCI studies used SDCs for correction. For instances in which no SDCs are available, a number of alternative SA correction methods have been proposed, e.g., based on spatial filters^15,16,22–24^ or by using individual baseline measurements for a filter based on principal component analysis.^17,25^

Awareness of the SA artifact and its need for correction is steadily growing and has already resulted in a number of publications proposing and/or comparing different correction methods.^13,15–17,21,23,24,26,27^ What is still scarce are performance comparisons of the proposed correction methods that can guide researchers in their choice. Particularly with respect to researchers who are not equipped with SDCs, there is a growing need for performance comparisons of methods with and without SDCs. To address this need, in this exploratory study, the performance of two spatial filters without SDC was compared with three regression-based SDC correction methods. Requirements for the chosen methods were that they are not only suitable for offline data analysis but also that they have a straightforward implementation in online scenarios, e.g., BCI or NFB applications. Also with consideration of BCI and NFB implementations, performance of the SA correction methods was evaluated at the single trial level wherever possible. Comparisons were based on measures quantifying signal improvement and spatial specificity. Expectations were that (i) signal quality will improve and spatial specificity will increase after the application of SAC methods compared with no correction, irrespective of whether SDCs were applied or not but that (ii) signal quality and spatial specificity improvements will be stronger for SAC methods with SDCs as compared with those without SDCs.

## 2 Materials and Methods

### 2.1 Subjects and Data Sets

For the current study two existing data sets were used. Because in real data the true underlying hemodynamic response function is unknown, the first data set was used to generate ground-truth semisimulated (SIM) data by adding a hemodynamic response function to resting state data (SIM DATA). The second data set consisted of real data of motor imagery (MI) and motor execution (ME). While MI data were chosen for their relevance for motor neurorehabilitation, ME data were included because of their more stable hemodynamic response.

#### 2.1.1 SIM data

Semisimulated data were generated based on resting-state measurements from *N* = 30 individuals that originated from an intermittent NFB study (unpublished). One participant was excluded because the rest data was missing. For the remaining 29 subjects, ~3 min of fNIRS data were recorded while subjects were looking at a black fixation cross on a gray background displayed on a computer screen. After applying the channel-based exclusion criteria of the present study (cf. Sec. 2.3.1 paragraph channel pruning for details), *N* = 23 older adults (11 females, 12 males; age [mean ± SD]: [65.52 ± 4.13] years; range: 58 to 71 years) were included for the final analyses.

#### 2.1.2 ME and MI data

The real data set comprised ME and MI data. Details of the study design can be found in Klein et al.^28^ In short, *N* = 32 healthy older adults performed five motor tasks, i.e., a self-paced sequential finger tapping task that was physically performed with either the left (ME LEFT) or the right (ME RIGHT) hand or kinesthetically imagined with either the left or the right hand. The fifth task was the kinesthetic imagination of an arbitrarily chosen whole-body movement involving both arms and legs. Each task consisted of 12 trials and the trial structure was a blocked design with 15 s of task period preceded and followed by 18 to 22 s of rest period. The order of tasks was pseudorandomized, with the restriction that not all of the three MI tasks appeared consecutively. Furthermore, a 15 s break after six trials of the same task and between two tasks was included in which subjects were allowed to slightly move their head, shoulders, and/or limbs if they felt that this helps for relaxation. All three MI tasks were combined into one MI task set (36 trials). After applying study specific exclusion criteria (cf. Sec. 2.3.1 paragraph channel pruning), *N* = 24 subjects (13 females, 11 males; age [mean ± SD]: [63.63 ± 4.73] years; range: 56 to 71 years) remained for analysis.

Participants of both studies gave written informed consent and were paid 10 Euros/h as reimbursement. General inclusion criteria for both studies were right-handedness, >4 h of sleep, and no history of neurological, psychiatric, or psychological diseases (cf. Klein et al.,^28^ for more details and all exclusion criteria). Both studies were approved by either the regular (resting-state data) or the medical (real data) Ethics Committee of the University of Oldenburg.

### 2.2 Functional Near-Infrared Spectroscopy (fNIRS)

FNIRS data were recorded using a NIRScout 816 device (NIRStar 15.2, NIRx Medizintechnik GmbH, Berlin, Germany) including 20 regular distance channels (in short: regular channels) and eight short distance channels (NIRx Medizintechnik GmbH, Berlin, Germany). The eight LED sources (intensity 5 mW/wavelength) had a distance of approximately 3 cm to the eight (regular distance) detectors and a distance of 0.8 cm to the short distance detectors. The regions of interests (ROIs) were bilateral M1 and SMA. For designing the probe layout, initially, ROI masks were generated based on the automated anatomical labeling (AAL) atlas as implemented in the wfupickatlas in SPM12 (v2.4^29,30^). Bilateral M1 was represented by left and right precentral gyrus (3D dilatation =1) and SMA was represented by bilateral supplementary motor area (3D dilatation = 1). The masks were loaded into the into the fNIRS Optodes’ Location Decider (fOLD) toolbox (v2.2^31^; https://github.com/nirx/fOLD-public) where the specificity was kept at 30% (brain atlas: SPM12). One restriction for the probe design was that the probe should be compatible with both the NIRScout system (i.e., our version with 12 detectors and 8 sources, one detector used for SDC bundle, diminishing the number of available detectors to 11; NIRx Medizintechnik, Berlin, Germany) as well as the NIRSport2 system (i.e., version with eight detectors and eight sources, one detector used for SDC bundle, diminishing the number of available detectors to 7; NIRx Medizintechnik, Berlin, Germany). The second restriction was that the distribution of the channels should be primarily focused on the SMA. To meet the criterion to be compatible with both fNIRS systems, optodes CCP5h and CCP6h were removed for analysis, resulting in 16 out of the 20 regular channels and eight SDCs [cf. Fig. 1(d) right]. The final probe layout is completely covered by the mask generated by the wfupickatlas and is visualized in Fig. 1(d) and matches with the following adjustments in the fOLD toolbox: For ROI M1, BrainAtlas: Brodmann; anatomical landmarks: four—primary motor cortex; specificity: 20%; for ROI SMA, both BrainAtlas: AAL2; anatomical landmarks: Supp_Motor_Area_L and Supp_Motor_Area_R; specificity: 10% or SMA portion of BrainAtlas: Brodmann; anatomical landmarks: six—premotor and supplementary motor area; specificity: 30%. The SMA channel distribution was slightly increased by additionally adding optodes to the EEG 10-5 positions FC1h and FC2h [cf. Fig. 1(d) left]. Note that ROI M1 (Brodmann area 4) and premotor areas (part of Brodman area 6) are highly overlapping and thus channels in M1 LEFT and M1 RIGHT might not exclusively represent M1. The NIRS optodes were placed according to the international 10-5 system in a custom-made cap, in which Cz, Nz, and Iz positions as well as left and right ears were used as markers for correctly positioning the cap.^32^ Optodes were attached to the cap with (spring loaded) grommets (NIRx Medizintechnik GmbH, Berlin, Germany) in order to reduce optode movement and to improve the contact between optodes and scalp. The sources emitted NIR light at wavelengths of 760 and 850 nm and raw light intensity was sampled at a rate of 7.8125 Hz.

**Fig. 1.**
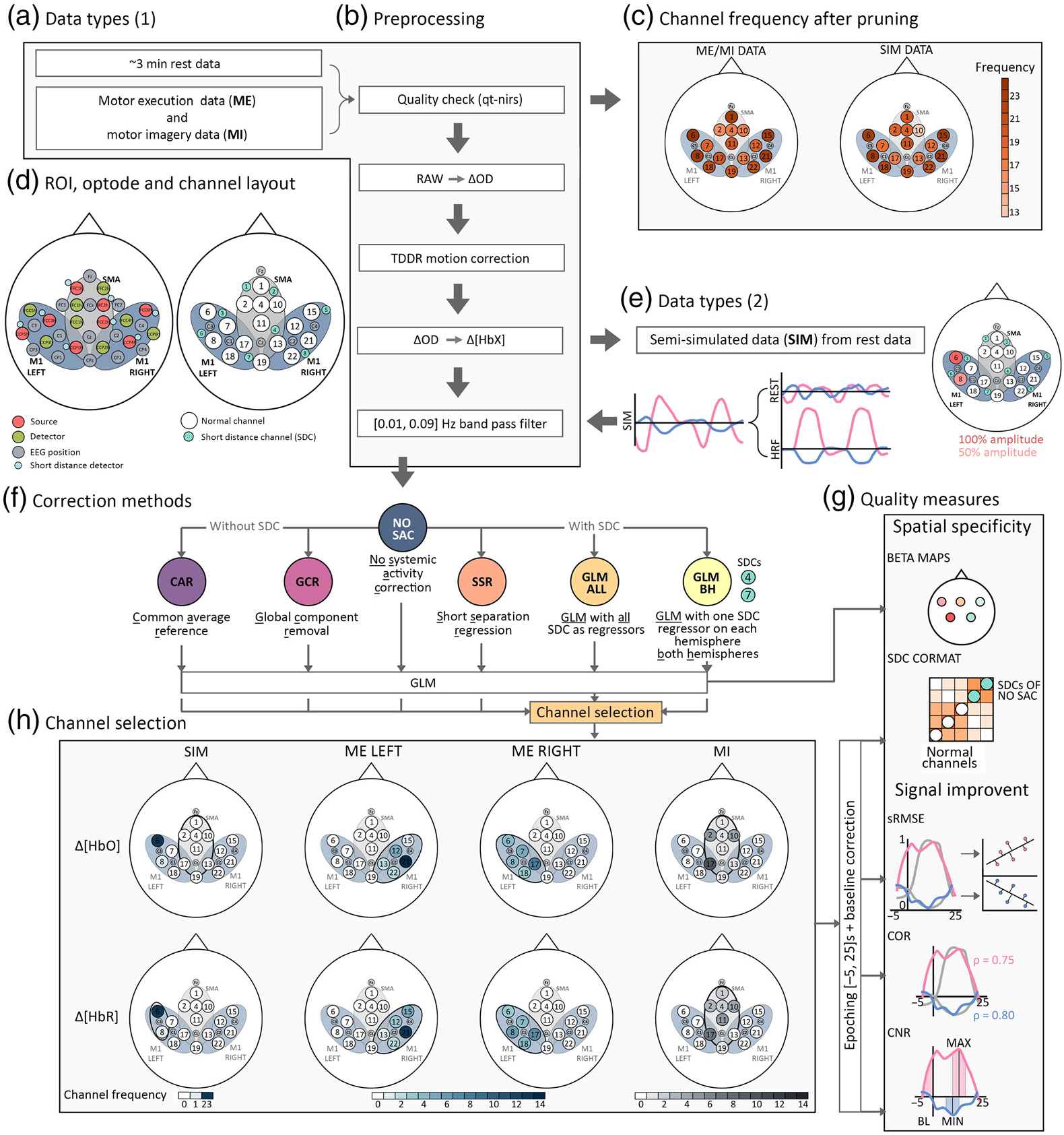
Illustration of the data processing stream in the present study. First, (a) both data sets underwent the (b) preprocessing stream of which the first step involved a quality check by using the qt-toolbox. Based on that, channels with poor quality were pruned. (c) The channel frequency across subjects after pruning for the SIM and the ME/MI data sets. (d) The probe layout and the channel layout including regular channels and short distance channels (SDCs) as well as the ROIs. (e) Semisimulated data were generated based on the rest data after the data were converted from raw data to optical density changes. Here, canonical HRFs were added to channels 6 and 8 with 100% and 50% amplitudes, respectively. After data were band-pass filtered, (f) the correction methods were applied, which resulted in six different data sets. NO SAC refers to the data that was preprocessed but was not corrected for SA. Correction methods without SDCs included the common average reference (CAR) and the global component removal (GCR) methods and methods with SDCs comprised the short separation regression (SSR), GLM with all SDCs as regressors (GLM ALL), and the GLM with one SDC (SDCs 4 and 7) regressor for each hemisphere (GLM BH). For each method, a GLM was performed. Based on the GLM output of GLM ALL the channel selection for the subsequent analyses was conducted. (h) The resulting channel frequencies across subjects for each task (columns) and data types (rows) are visualized. Data of the selected channel were then epoched and baseline corrected and finally, based on single trials the (g) quality measures for signal improvement sRMSE, COR, and CNR and for spatial specificity SDC CORMAT were calculated. Spatial specificity measure BETA MAPS was not based on single trials but on the individual GLM output resulting from each method.

#### 2.2.1 Additional unused data

Optode positions and fiducial points were digitized with an optical digitizer (Xensor, ANT Neuro, The Netherlands). Because most of the digitizer data of the SIM data set was corrupted due to missing or inaccurate optode locations, digitzer data was not further considered. Moreover, the data sets contain electromyography data that were used in a previous publication^28^ but are of no relevance for the present study.

### 2.3 Data Processing and Statistical Analysis

#### 2.3.1 fNIRS analysis

FNIRS data processing was conducted with a combination of the NIRS Brain AnalyzIR toolbox^33^ (https://github.com/huppertt/nirs-toolbox) and custom made scripts. For a better overview, in Fig. 1 the workflow from data types (A, E) over the preprocessing pipeline (B), the applied correction methods (F) to the generated quality measures (G) is visualized.

##### Quality check and channel pruning

Channel quality was assessed by means of the qt-nirs toolbox that uses the scalp coupling index (SCI^34,35^) and the peak spectral power (PSP^35^) for evaluating signal quality (SCI threshold = 0.6, *Q* threshold = 0.65, and PSP threshold = 0.1; https://github.com/lpollonini/qt-nirs). Note that the GLM-based correction methods are using the AR-IRLS model^36^ (cf. Sec. 2.3.2), which is relatively robust against channels with poor signal quality and thus these channels do not necessarily need to be removed. However, as the quality assessment is important for other correction methods it was however performed as a general step in the processing pipeline. The first inclusion criterion was that all short-distance channels remained for analyses, which was the case for 25 participants of the SIM data set and for 24 of the ME/MI data set. For the SIM data, the second inclusion criterion concerned the consistent number of identical channels for adding the simulated HRF across subjects. Here, the aim was to find the same two channels across subjects that remain within an ROI after channel pruning. This was the case for channels 6 and 8 of ROI M1 LEFT in *N* = 23 subjects [cf. Figs. 1(d) and 1(e)]. Across the final sample of the SIM data, on average (±SD) 2.26 (± 2.56; ranging from 0 to 8 channels) regular channels were pruned. Pruned channels were replaced by NaNs in the respective data set.

For the ME/MI data, the second inclusion criterion was that in each of the three ROIs at least two channels remained for analysis after pruning. Based on this, all *N* = 24 subjects were considered for analysis and on average across subjects 2.71 of all regular channels (SD± 3.20; ranging from 0 to 9 channels) were pruned. Data of the pruned channels were replaced by NaNs in the respective data set. A visualization of the frequency of the remaining channels of both data sets after pruning can be found in Fig. 1(c) and exact numbers are documented in Tables S21–S26 in the Supplementary Material.

##### Preprocessing

Preprocessing was applied to all channels, including both short and regular-distance channels. Raw data were transformed into optical density changes and were then corrected for possible motion artifacts by means of the temporal derivative distribution repair (TDDR) method.^37^ Afterwards, data were converted into hemoglobin concentration changes by means of the modified Beer–Lambert law (mBLL). Therefore, the individual age-related^38^ differential pathlength factor (DPF) was used in order to calculate the partial pathlength factor [PPF = DPF × partial volume factor (PVF = 1/60)], which was used for the mBLL. To generate the SIM data, the canonical HRF was hereupon added to the rest data [see following paragraph *Generation of semisimulated (SIM) data* for details]. Then, data were first low-pass filtered with a zero-phase digital Butterworth filter (cut-off = 0.09 Hz; order = 2) and then high-pass filtered with a zero-phase digital Butterworth filter (cut-off = 0.01 Hz; order = 2). The cut-off frequencies of [0.01, 0.09] Hz were selected based on the recommendations by Pinti et al.;^39^ however, departing from this, in the present data analysis, no finite impulse response (FIR) filter with a high filter order (>1000) was applied because the SIM data had no sufficient length for this filter order (~3 min of rest data) and we wanted to keep the preprocessing the same across data sets.

##### Generation of semisimulated (SIM) data

Resting-state data of the first data set were used to generate the SIM data. Canonical HRFs (peak at 6 s) with different amplitudes were generated and added to the rest data of neighboring channels 6 and 8, located in the ROI M1 LEFT [cf. Fig. 1(c)]. Semisimulated data were generated for two channels and with different amplitudes to simulate a spatial pattern with a signal strength gradient (cf. Sec. 2.4.3). The amplitude for channel 6 was 10 *μ*M for Δ[HbO] and 10/3 *μ*M for Δ[HbR] (100% amplitude) and 5 *μ*M for Δ[HbO] and 5/3 *μ*M for Δ[HbR] (50% amplitude) for channel 8. For each subject, a total of five trials were simulated as a blocked design with 15 s of task period preceded and followed by 18 to 22 s of rest period.

#### 2.3.2 Systemic activity correction (SAC) methods

##### No systemic activity correction (NO SAC)

Data after the band-pass filtered preprocessing stage served as a reference measure and are referred to as no systemic activity corrected (NO SAC) data.

###### Common average reference (CAR)

A very simple correction method without the use of SDCs is the common average reference (CAR) spatial filter, which has its origin in EEG analysis.^22^ Based on the idea that a global signal is present in all channels, a spatial filter can be generated by simply subtracting the average time signal across all *N* channels from each single channel **x**_*i*_, (*i* = 1,….,*N*) to reduce the global signal influence, resulting in the corrected channels **x**_CAR_*i*__ [cf. Eq. (1)]^22^

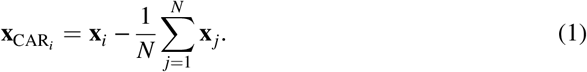

###### Global Component Removal (GCR)

The second correction method without using SDCs is the global component removal (GCR) method.^23,24^ The GCR is a more advanced spatial filter based on Gaussian spatial filtering and singular value decomposition (SVD). The Gaussian kernel smoother *G* is a two-dimensional kernel smoother that is applied to remove detail and noise from a data set.^23^ It requires the Montreal Neurological Institute (MNI) coordinates of each channel that are stored in a N × N distance matrix *D*

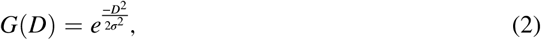

where *σ* represents the width of the kernel and was shown to be effective with a value of *σ* = 46 deg.^40^ The SVD decomposes the data matrix *Y*_task_ into the three matrices *U*, Σ, and *V^T^*

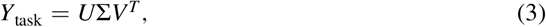

where the *n* × *m* matrix *U* represents the temporal information of the *n* channels, the *n* × *n* diagonal matrix Σ consists of the singular values, i.e., the square root of the eigenvalues, and the *n* × *n* matrix *V^T^* represents the transpose of the spatial information. The vectors **v**_*i*_ of *V* are then smoothed by the Gaussian kernel *G*

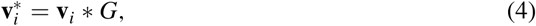

which results in the vectors 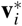 of matrix *V*\*. The convolution with G removes the localized neuronal pattern and, therefore, *V*\* represents only the spatial information of the global component.^23^ By replacing *V* with *V*\* in the SVD formula, the waveforms of the GC *Y*_global_ can be reconstructed

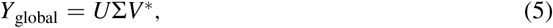

and to get the neuronal component *Y*_neuronal_, the GC *Y*_global_ can then simply be subtracted from the data *Y*_task_

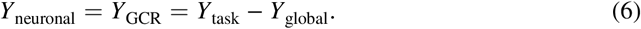

###### Short Separation Regression (SSR)

A simple and easy to implement SAC method that involves SDCs is the short separation regression (SSR).^11,19^ By simply applying Eq. (7), it was shown that SA can be reduced

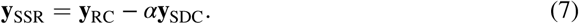

Here, *α* is defined as the quotient of two scalar products (〈·,·〉)

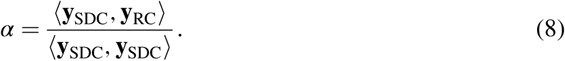

In this method, **y**_SDC_ can be either the closest SDC, the SDC showing the strongest correlation with the regular channel **y**_RC_ or an average across all available SDCs. In the present study **y**_SDC_ represents the signal of the closest SDC to each regular channel determined by the Euclidian distance.

###### General Linear Model with all SDCs as Regressors (GLM ALL)

The most recommended SAC method when using SDCs is to incorporate the SDCs as regressors within the general linear model (GLM).^13,17^ With this approach one can add the SDC regressors X_SDC_ in addition to the task-related regressors X_task_ directly to the GLM analysis

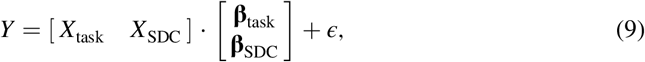

where **β**_task_ and **β**_SDC_ represent the beta values of the task and of the SDC, respectively, and *ϵ* the residuals of that model. However, in an NFB or BCI experiment, one might be more interested in further processing the corrected time signal data; therefore, the GLM can be applied for SAC by adding only the SDC channels to the design matrix X_SDC_

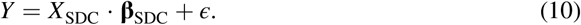

In this model, each SDC time course in *X*_SDC_ results in a beta value in **β**_SDC_ estimating its contribution to the time signals in *Y* and the part that can not be explained by the model is included in the residual term *ϵ*. By subtracting the product of design matrix *X*_SDC_ and beta values **β**_SDC_ from the data *Y, ϵ* results in the cleaned data set *Y*_GLMALL_

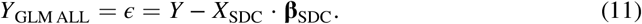

As recommended by Santosa and colleagues,^17^ for the GLM ALL method all eight SDCs of both Δ[HbO] and Δ[HbR] were included in the present study as regressors for running the GLM. For solving the GLM, an autoregressive iterative least square (AR-IRLS^36^) algorithm implemented in the Brain AnalyzIR toolbox^33^ (https://github.com/huppertt/nirs-toolbox) was applied.

###### General Linear Model with one SDC on Both Hemispheres as Regressors (GLM BH)

Similar to the GLM ALL method, the GLM BH correction method uses SDCs as regressors within the GLM in order to correct for SA. However, here only two SDCs were used for correction to pretend to be in a scenario where only a small number of SDCs are available.^41,42^ As for GLM ALL, by subtracting the product of design matrix *X*_SDC_BH__ and beta values **β**_SDC_BH__ from the uncorrected data Y, ε results in the cleaned data set Y_GLMBH_

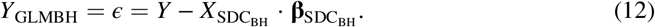

For the GLM BH method, SDCs 4 and 7 were chosen because they cover both hemispheres (BH) and were included in all of the three ROIs [cf. Fig. 1(d)]. Also for this GLM method, regressors of both Δ[HbO] and Δ[HbR] of the respective channels were included in the model.

### 2.4 Channel Selection and Quality Measures

#### 2.4.1 Channel selection

The majority of quality measures were calculated for selected channels. To enhance comparability across SDC methods, the same channel set was used for all correction methods. Channels were not selected from the whole channel pool but selection was restricted to the task-specific ROI. The task-specific ROI was defined *a priori* as the ROI that most likely shows the strongest activation for a given task. For ME LEFT the task-specific ROI was M1 RIGHT, for ME RIGHT and SIM it was M1 LEFT for MI it was SMA.

For each subject, the best channel was selected separately for each task (SIM, ME LEFT, ME RIGHT, and MI) and signal type (Δ[HbO] and Δ[HbR]), resulting in four tasks × two signal types = eight channels per subject. The channel selection was based on the individual output of two AR-ILS^36^ GLMs (one for SIM data and one for real data) run on the GLM ALL corrected data because it was expected that this correction method results in the cleanest data.^13,17^ For tasks ME LEFT, ME RIGHT, and MI, besides the three task-related regressors, regressors of the instruction and the breaks were added to the GLM. Additionally, first and second derivatives of each regressor were included in the model. For SIM data, the GLM included one task-related regressor consisting of all simulated onsets. From the GLM outputs, the individual channels with the highest (Δ[HbO]) and lowest (Δ[HbR]) beta values were selected from within the task-specific ROIs. The resulting channel frequencies across tasks and signal types are visualized in Fig. 1(h).

#### 2.4.2 Quality measures for signal improvement

Signal improvement was assessed with three different measures expected to be reflective of single trial noise reduction after applying SAC. It was expected that for SAC methods that effectively remove SA, quality measures for signal improvement should be enhanced relative to uncorrected data. It was also expected that the measures demonstrate a superiority of SDC methods compared with methods without SDCs.

For the individually selected channels (see Sec. 2.4.1), measures were calculated for data epoched [−5,25] s around stimulus onset and baseline corrected with the average of the signal from five seconds prior to stimulus onset. For ME LEFT and ME RIGHT, quality measures were first calculated separately and then averaged for each subject to generate a single value termed ME.

##### Scaled root mean square error (sRMSE)

For this quality measure, a linear regression was fitted (using the function lm() of the ‘stats’ package v4.0.2^43^ and rescale() of the ‘scales’ package v1.1.1 [https://github.com/r-lib/scales] in R^43^ in combination with RStudio^44^) between a single trial epoch of either Δ[HbO] or Δ[HbR] scaled to [0, 1] **y**_*s*_ and a single trial canonical HRF scaled to [0, 1] **x**_*s*_

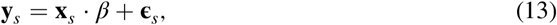

where *β* represents the strength of the fit and the residual term **∈**_*s*_ is a measure of the error of the fit. The scaling of the epochs was performed using the min–max normalization

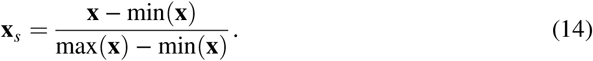

Then, the residuals **∈**_*s*_ of the model fit were extracted and the sRMSE was calculated as follows:

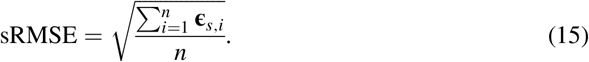

For each subject and selected channel, the single trial sRMSE values were then averaged across trials to have a single value per subject. When comparing different models resulting from different SAC methods, a low sRMSE value indicates low error of the corresponding model and is therefore interpreted as stronger signal improvement as compared with a model with a higher sRMSE value. Note that the RMSE is an important and often applied measure to validate the performance of a linear regression model. However, since the RMSE is a scale-dependent measure, it is only meaningful when comparing models based on the same data.^45–47^ If it is of interest to compare several models based on different data sets, as in the present study, a scaling or normalization step should be performed.^47^

##### Correlation (COR)

For this quality measure, the Spearman rank correlation coefficient *ρ* between a single trial epoch of either Δ[HbO] or Δ[HbR] *y* and the corresponding canonical HRF *x* was calculated (using the function cor() of the ‘stats’ package v4.0.2^43^ in R^43^ in combination with RStudio^44^). To use this value for statistical analysis, each single trial *ρ* was then Fisher’s z-transformed (function FisherZ() of the ‘DescTools’ package v0.99.38 [https://github.com/AndriSignorell/DescTools] in R^43^ in combination with RStudio^44^) and for each subject and selected channel averaged across trials. With respect to the COR quality measure, when comparing two methods, the method with the higher COR value indicates a stronger relationship between HRF and trial and therefore a stronger signal improvement.

##### Contrast-to-noise ratio (CNR)

The third single trial quality measure for signal improvement is the contrast-to-noise ratio^25,48,49^

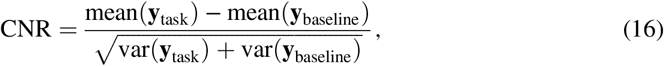

where mean and var indicate the mean and variance of either the single trial baseline **y**_baseline_ or the task period **y**_task_. As compared with previous implementations,^25,48,49^ in the present study, the task period was slightly adapted by using the averaged data of the peak value ±2 s of a given trial within the time window of [−5,25] s around stimulus onset [using a custom-made function in MATLAB based on the MATLAB functions mean() and var()]. For each subject and selected channel, the single trial CNR were then averaged across trials. The higher the CNR value the larger the ratio of task-related signal to noise and, correspondingly, the better data quality.

#### 2.4.3 Quality measures for spatial specificity

Spatial specificity characterizes how well defined a signal is in space. To prepare the data, the time series signal of all regular channels were epoched [−5,25] s around stimulus onset and baseline corrected with the average of the signal from 5 s prior stimulus onset. In addition, the beta values resulting from the GLMs of each correction method were extracted for both Δ [HbO] and Δ[HbR].

##### Correlation matrices with SDCs (SDC CORMAT)

For each single trial, a correlation matrix^16^ [Spearman correlation using the MATLAB function corr()] was calculated based on the data of each of the 16 regular channels concatenated to the data of the eight SDCs from the NO SAC data. Matrices consisted thus of correlations between all regular channels as well as between SDCs and all regular channels. For pruned channels, correlations were replaced by a NaNs. The single trial matrices were then averaged [using function *nanmean()*] first within a subject and then across subjects separately for each method (NO SAC, CAR, GCR, SSR, GLM ALL, and GLM BH), data type (Δ[HbO] and Δ[HbR]) and task (SIM, ME LEFT, ME RIGHT, and MI), resulting in a total of 6 × 2 × 4 = 48 correlation matrices with dimensions 24 × 24. A more spatially specific signal would, on a descriptive level, be indicated by (i) reduced correlations between regular channels and SDCs as well as (ii) exclusively high correlations in the respective task-specific ROIs.

##### Topographical beta maps (BETA MAPS)

For this quality measure no single trial data were considered, instead, the whole time series was of interest. For both data sets (SIM data and real data) and each correction method a separate AR-ILS^36^ GLM was applied, resulting in two data sets × six correction methods = 12 GLMs per subject. For the SIM data one task-related regressor consisting of all simulated onsets was added to the model. For the real (ME/MI) data, besides the three task-related regressors (ME LEFT, ME RIGHT, and MI) also regressors of the instruction and the breaks were added. Additionally, first and second derivatives of each regressor were included in the model.

Beta maps^28^ were generated based on the GLM outputs by averaging beta values [using the MATLAB function nanmean()] for each regular channel across subjects separately for SDC method (NO SAC, CAR, GCR, SSR, GLM ALL, and GLM BH), data type (Δ[HbO] and Δ[HbR]), and task (SIM, ME LEFT, ME RIGHT, and MI), resulting in a total of 6 × 2 × 4 = 48 beta maps. Beta values of pruned regular channels were replaced by an NaNs. For the topographical beta maps, spatial specificity would be judged as poor if most of the channels show strong activation and as good if mainly channels of the expected task-specific ROI show strong activation. That is, for the SIM data, only channels 6 and 8 should be activated, for ME LEFT channels of ROI M1 RIGHT, for ME RIGHT channels of ROI M1 LEFT, and for MI channels in ROI SMA should be activated.

### 2.5 Statistical Analyses

Statistical analyses were conducted with MATLAB 2019a, R (version 4.0.2, “Taking Off Again”^43^) in combination with RStudio (v1.3.1093^44^) and Jeffreys’s Amazing Statistics Program (JASP; v0.16.1^50^). Bayesian ANOVAs (BANOVAs) and corresponding posthoc tests were conducted in JASP, and individual Bayesian t-tests were performed in R in combination with the *BayesFactor* package (function *ttestBF*; v0.9.12-4.2^51^).

Except for generating the correlation matrices, all statistical analyses were conducted within the framework of Bayesian statistics instead of the inferential framework. The advantage of using Bayesian statistics over the frequentist approach is that it allows to interpret both the alternative hypothesis (*H*_1_) and the null hypothesis (*H*_0_). Note that instead of reporting a p-value in the Bayesian framework, a Bayes factor (*BF*_10_) helps to quantify how much more likely the data are under either the *H*_0_ or the *H*_1_.^52,53^ Moreover, as BF_10_ reflects how likely the evidence for either *H*_0_ or *H*_1_ is, correction for multiple comparisons is not required.^54^ In the present work, the Bayes factors BF_10_ will be classified into different categories following the guidelines by Lee and Wagenmakers,^52^ which are as follows: BF_10_ < 100 extreme evidence for 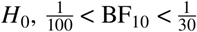 very strong evidence for 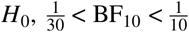 strong evidence for 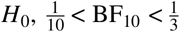 moderate evidence for 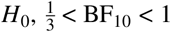 anecdotal evidence for 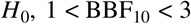 anecdotal evidence for 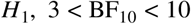 moderate evidence for *H*_1_, 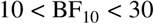 strong evidence for *H*_1_, 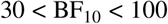 very strong evidence for *H*_1_, BF_10_ > 100 extreme evidence for *H*_1_.

For better readability, only a subset of the results of the vast number of statistical comparisons are reported in the main body of the manuscripts. These are results with Bayes factors >3 (moderate to extreme evidence in favor of *H*_1_). However, all statistical results as well as the mean value (±SD) of each variable of interest are reported in tables in the Supplementary Material.

#### 2.5.1 Signal improvement

For the comparison of quality measures sRMSE, COR, and CNR for each task (SIM, ME, and MI) and signal type (Δ[HbO] and Δ[HbR]) a one factorial Bayesian repeated-measures ANOVA (rmBANOVA) with the factor METHOD (NO SAC, CAR, GCR, SSR, GLM ALL, and GLM BH) compared with the null model was conducted and Bayes factors BF_10_ were reported. Prior odds *P*(*M*) were set as the default of *P*(*M*) = 0.5. Posthoc analyses were only conducted if the rmBANOVA showed at least moderate (BF_10_ > 3) evidence favoring the alternative hypothesis (*H*_1_). As implemented by default in JASP, the posthoc analyses are based on the default t-test with a Cauchy 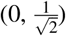 prior, i.e., prior odds were set as the default value of *P*(*M*) = 0.26 and posterior odds *P*(*M*|data) were corrected for multiple testing by fixing to 0.5 the prior probability that the *H*_0_ holds across all comparisons.^50^

For all three measures the general expectation was that (i) all correction methods show lower (sRMSE) or higher (COR and CNR) values, respectively, as compared with NO SAC and that (ii) values resulting from correction methods with SDCs (SSR, GLM ALL, and GLM BH) are lower (sRMSE) or higher (COR and CNR), respectively, as compared with values resulting from correction methods without SDCs (CAR and GCR).

#### 2.5.2 Spatial specificity

##### Correlation Matrices with SDCs (SDC CORMAT)

On a purely descriptive level, it was expected that data from corrected channels show less correlation with the SDCs and that matrices show higher spatial specificity by showing stronger correlations within ROIs as compared with the NO SAC data.

For statistical analysis, a Bayesian paired *t*-test was performed on the lower triangular part of the correlation matrices for any two pairs of correction methods (total of 15 pairs) within task (SIM, ME LEFT, ME RIGHT, and MI) and signal type (Δ[HbO] and Δ[HbR]). In total, 15 × 4 tasks × 2 signal types = 120 t-tests were conducted for which Bayes factors BF_10_ were reported. The default of a medium prior with 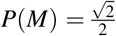 was applied. Across tasks and signal types, we expected (i) that matrices resulting from any correction method will differ from the correlation matrix of NO SAC as well as (ii) differences between correlation matrices resulting from methods with SDC (SSR, GLM ALL, and GLM BH) and those resulting from methods without SDCs (CAR and GCR).

###### Topographical beta maps (BETA MAPS)

For visual inspection and qualitative comparisons, the beta values of the GLMs were used to generate average topographical beta maps for each method (NO SAC, CAR, GCR, SSR, GLM ALL, and GLM BH), signal type (Δ[HbO] and Δ[HbR]), and task (SIM, ME LEFT, ME RIGHT, and MI), resulting in a total of six methods × two signal types × four tasks = 48 topographical beta maps. For statistical comparisons, for each of the 16 regular channels of any of these beta maps an one-sample Bayesian *t*-test was conducted by using the default of a medium prior with 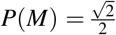 and Bayes factors BF_10_ are reported.

On a purely descriptive level, the general expectation within each task and signal type was that (i) spatial specificity increases after applying any SAC methods as compared with NO SAC and that (ii) methods with SDC correction show a higher spatial specificity as compared with methods without SDCs. Statistically, this would be reflected by the strongest Bayes factors in channels 6 and 8 of the SIM DATA and for ME LEFT and ME RIGHT by the strongest Bayes factors resulting from channels on the hemisphere contralateral to the performing hand. For MI data, the expectation is less strong, but the general expectation would be to get the strongest Bayes factors for ROI SMA.

## 3 Results

### 3.1 Signal Improvement

#### 3.1.1 Scaled root mean square error (sRMSE)

The mean sRMSE values (±*SEM*) are visualized in Fig. 2. Descriptively, the sRMSE values of Δ[HbO] show only for the SIM data a clear difference between data of all SAC methods and the data of NO SAC [cf. Fig. 2(a)]. For Δ[HbR], the pattern for the correction methods of the SIM data is less clear. However, mean sRMSE values of all correction methods seem to decrease as compared with NO SAC [cf. Fig. 2(b)]. For both data types of ME and MI data, descriptively, the mean sRMSE values either increase for the correction methods or seem to be comparable to those of the NO SAC data.

**Fig. 2.**
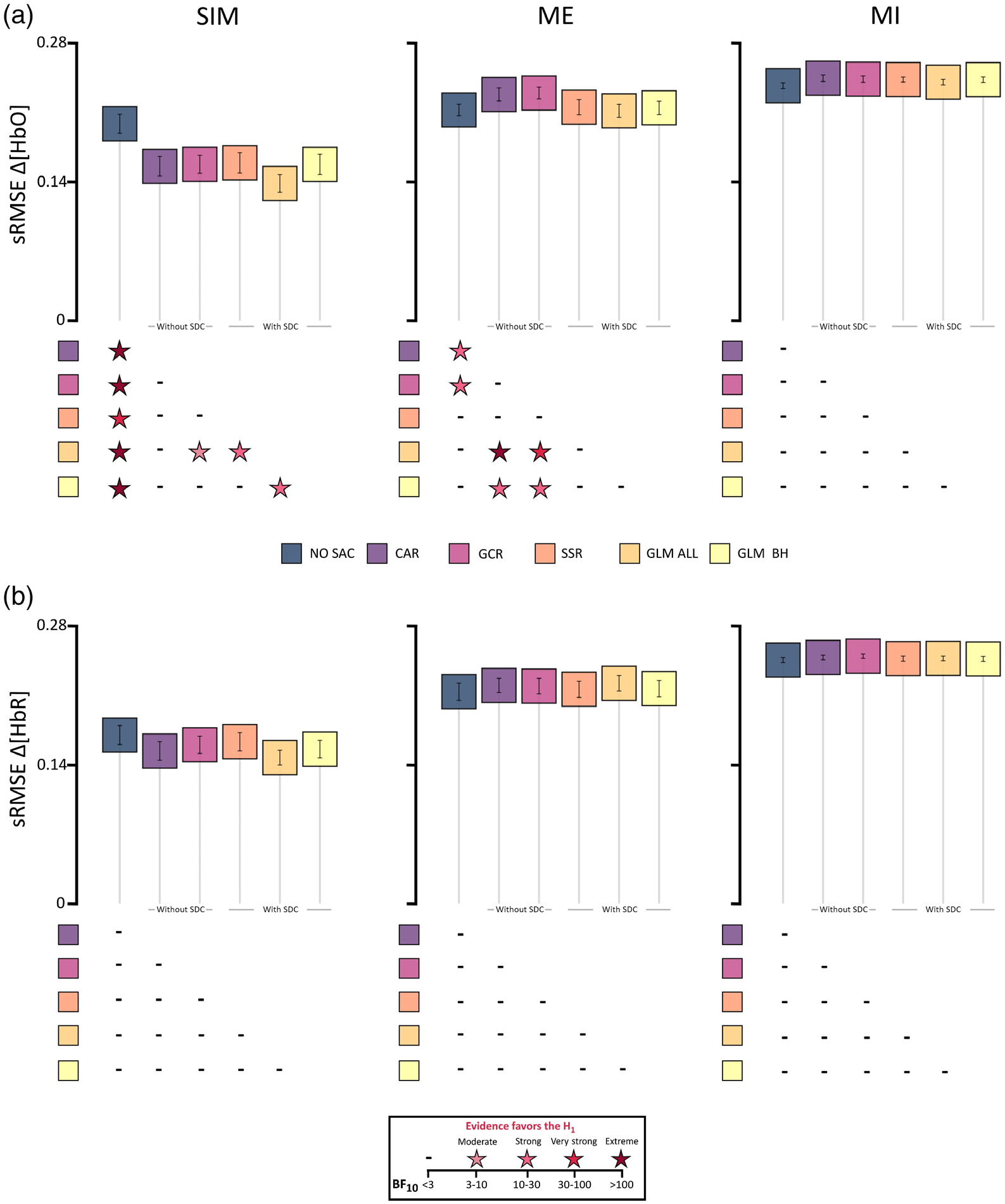
Illustration of the sRMSE results including all three data types (columns; SIM, ME, and MI) of (a) Δ[HbO] and (b) Δ[HbR]. Boxes represent mean sRMSE values and error bars represent standard error of the mean (SEM) across participants. Below each subfigure statistical results are displayed by color coded stars representing Bayes factors BF_10_ > 3, which indicate evidence favoring the alternative hypothesis *H*_1_, i.e., that there are differences between two correction methods. For details of statistical results including BF_10_ < 3, see Tables S1–S6 in the Supplementary Material.

##### SIM data

Detailed statistical results are reported in Tables S1 and S2 in the Supplementary Material. With respect to the Δ[HbO] data, the rmBANOVA revealed extreme evidence favoring the *H*_1_. Post hoc tests revealed very strong to extreme evidence that all of the correction methods differed from NO SAC in that their sRMSE values were lower than that of NO SAC (**all** < **NO SAC**). Additionally, the comparisons between GLM ALL and SSR, as well as GLM BH, indicated strong evidence and between GLM ALL and GCR moderate evidence for a difference in favor of GLM ALL (**GLM ALL** < **SSR, GLM BH, GCR**).

For Δ [HbR], data did not provide evidence for a difference in sRMSE between SAC methods. Instead, rmBANOVA indicated anecdotal evidence favoring the *H*_0_.

##### ME Data

Detailed statistical results are reported in Tables S3 and S4 in the Supplementary Material. For the Δ[HbO] data, the rmBANOVA revealed extreme evidence favoring the *H*_1_. Posthoc tests indicated that evidence was strong to very strong evidence for a raised sRMSE for GCR. This included the comparisons between GCR and NO SAC, GCR and GLM ALL, and GCR and GLM BH (**GCR** > **NO SAC, GLM ALL, GLM BH**). Similarly, there was strong evidence for a higher sRMSE for CAR compared with NO SAC, CAR compared with GLM BH, and extreme evidence for CAR compared with GLM ALL (**CAR** > **NO SAC, GLM ALL, GLM BH**). Note that for the comparisons with NO SAC the opposite direction was expected, that is, expected were lower sRMSE values for GCR and CAR as compared with NO SAC.

For Δ [HbR], data did not provide evidence for a difference in sRMSE between SAC methods. Instead, rmBANOVA indicated anecdotal evidence favoring the *H*_0_.

##### MI Data

Detailed statistical results are reported in Tables S5 and S6 in the Supplementary Material. For the Δ[HbO] data, the rmBANOVA revealed anecdotal evidence favoring the *H*_0_. Regarding the Δ[HbR] data, the rmBANOVA revealed moderate evidence favoring the *H*_0_.

#### 3.1.2 Correlation (COR)

The means of the Fisher’s *z*-transformed Spearman correlations COR (±SEM) are illustrated in Fig. 3. Descriptively, for both Δ[HbO] and Δ[HbR] SIM data mean COR values were higher for all SAC methods as compared with NO SAC. For ME Δ[HbO] data, only the SAC methods with SDCs showed an increase in COR as compared with NO SAC, whereas both methods without SDCs showed a decrease in COR. For Δ[HbR] of ME data and both Δ[HbO] and Δ[HbR] of the MI data, COR values of the SAC methods either decreased or were comparable to NO SAC.

**Fig. 3.**
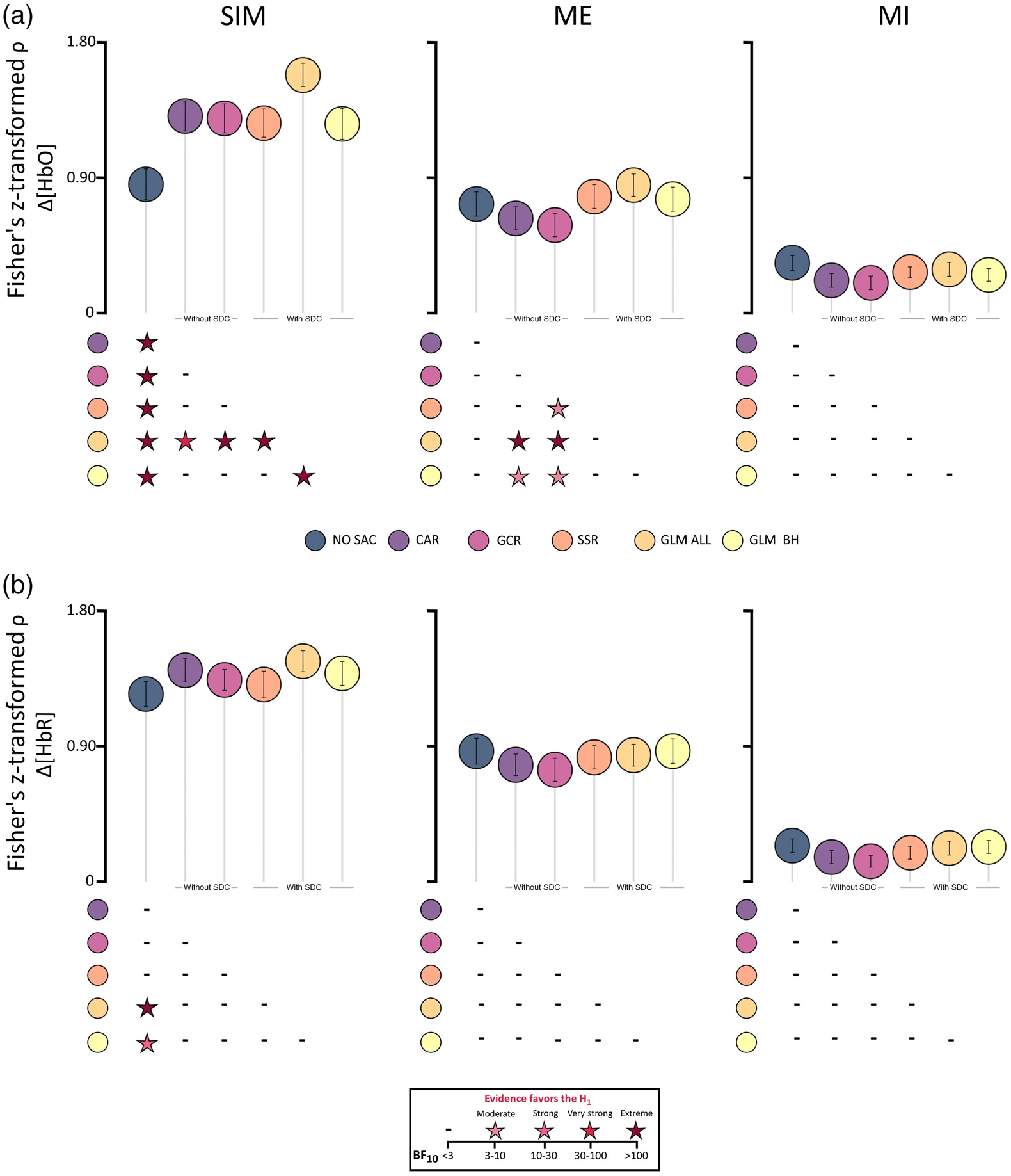
Illustration of the Fisher’s z-transformed Spearman correlation (COR) results including all three data types (columns: SIM, ME, and MI) of (a) Δ[HbO] and (b) Δ[HbR]. Circles represent mean COR values and error bars represent standard error of the mean across participants. Below each subfigure statistical results are displayed by color coded stars representing Bayes factors BF_10_ > 3 which indicate evidence favoring the alternative hypothesis *H*_1_, i.e., that there are differences between two correction methods. For details of statistical results including BF_10_ < 3, see Tables S7-S12 in the Supplementary Material.

##### SIM data

Detailed statistical results are reported in Tables S7 and S8 in the Supplementary Material. For Δ[HbO], the rmBANOVA revealed extreme evidence favoring the *H*_1_. Posthoc tests indicated extreme evidence for the differences between NO SAC and all other correction methods, resulting from higher COR values of the correction methods as compared with NO SAC. For GLM ALL, there was also extreme evidence for higher COR values compared with GCR, SSR, and GLM BH and very strong evidence for higher COR values compared with CAR (**GLM ALL** > **GCR, SSR, GLM BH, and CAR**).

For Δ[HbR], the rmBANOVA revealed moderate evidence favoring the *H*_1_. Posthoc tests indicated higher COR values for GLM ALL and GLM BH compared with NO SAC, with extreme evidence for the difference between NO SAC and GLM ALL and strong evidence for differences between GLM BH and NO SAC (**GLM ALL and GLM BH** > **NO SAC**).

##### ME data

Detailed statistical results are reported in Tables S9 and S10 in the Supplementary Material. For the Δ[HbO] data, the rmBANOVA revealed extreme evidence favoring the *H*_1_. Posthoc tests indicated extreme evidence for differences between GLM ALL and and GCR and between GLM ALL and CAR with higher COR values for GLM ALL in both comparisons (**GLM ALL** > **GCR, CAR**). Moreover, there was evidence for particularly low COR values for GCR, with moderate evidence for differences between GCR and SSR and between GCR and GLM BH (**SSR, GLM BH** > **GCR**). There was also moderate evidence for the difference between CAR and GLM BH, with higher COR values for GLM BH (**GLM BH** > **CAR**).

With respect to the Δ[HbR] data, the rmBANOVA revealed anecdotal evidence favoring the *H*_1_.

##### MI data

Detailed statistical results are reported in Tables S11 and S12 of the Supplementary Material. For both Δ[HbO] and Δ[HbR], the rmBANOVAs revealed anecdotal evidence favoring the *H*_1_.

#### 3.1.3 Contrast-to-noise ratio (CNR)

The mean CNRs (±SEM) are illustrated in Figs. 4(a) and 4(b). Descriptively, with respect to the SIM data of both Δ[HbO] and Δ[HbR] all SAC methods show increased CNR values as compared with NO SAC. For Δ[HbO] of the ME data, all SAC methods with SDCs show increased CNR values as compared with NO SAC and for Δ[HbR] of the ME data, only both GLM methods show increased CNR values. In contrast, methods without SDCs show slightly decreased CNR values as compared with NO SAC. With respect to both data types of the MI data, all methods seem to be comparable with NO SAC.

**Fig. 4.**
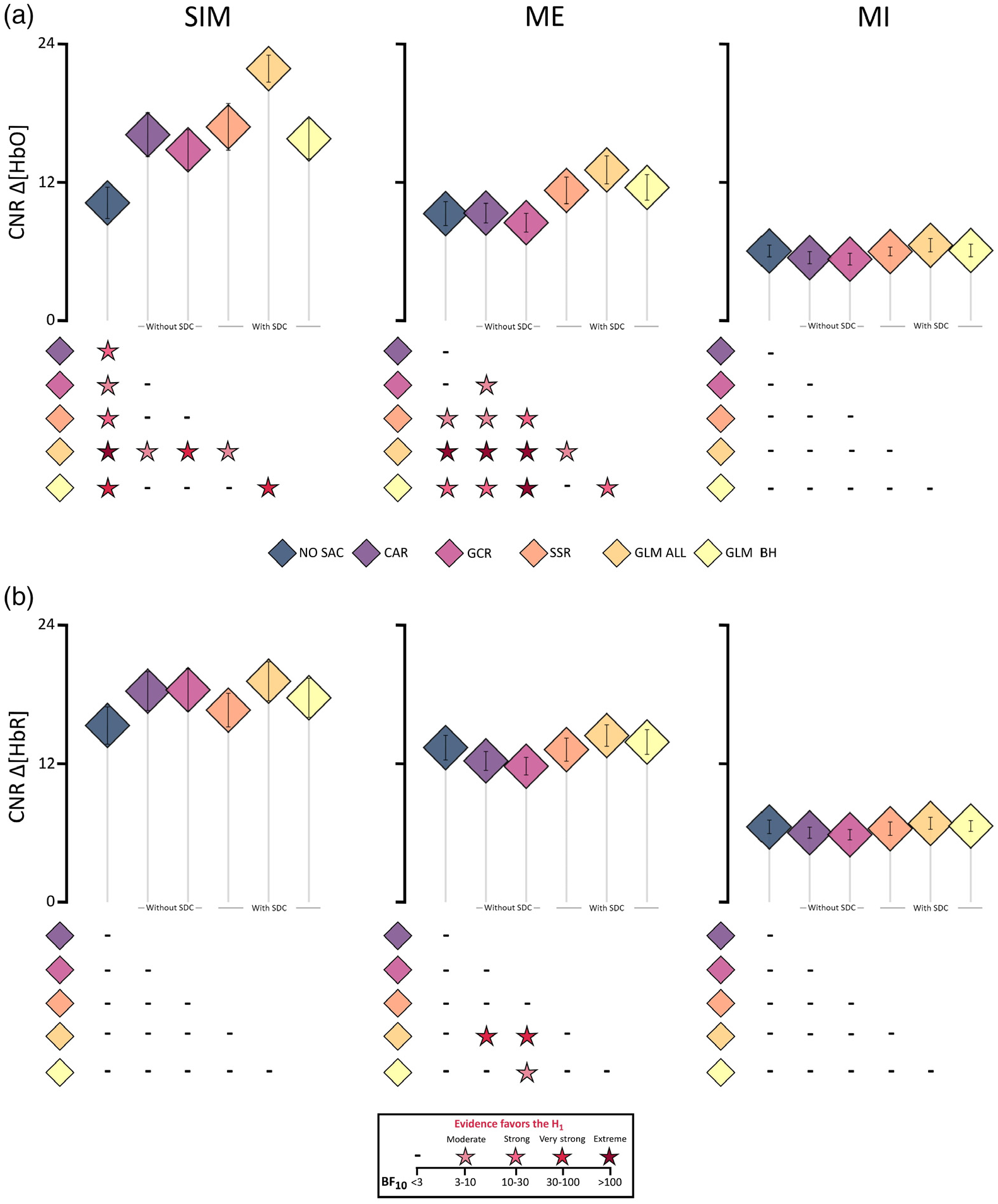
Illustration of the contrast-to-noise ratio (CNR) results including all three data types (columns: SIM, ME, and MI) of (a) Δ[HbO] and (b) Δ[HbR]. Diamonds represent mean CNR values and error bars represent standard error of the mean across participants. Below each subfigure statistical results are displayed by color coded stars representing Bayes factors BF_10_ > 3, which indicate evidence favoring the alternative hypothesis *H*_1_, i.e., that there are differences between two correction methods. For details of statistical results including BF_10_ < 3, see Tables S13–S18 in the Supplementary Material.

##### SIM data

Detailed statistical results are reported in Tables S13 and S14 in the Supplementary Material. With respect to the Δ[HbO] data, the rmBANOVA revealed extreme evidence favoring the *H*_1_. As compared with NO SAC, posthoc tests revealed extreme evidence for differences to GLM ALL, very strong evidence for differences to GLM BH, strong evidence for differences to CAR and SSR and moderate evidence for differences to GCR, all resulting from higher CNR values in favor of the correction methods as compared with NO SAC (**NO SAC** < **CAR, GCR, SSR, GLM BH, and GLM ALL**). Moreover, there was very strong evidence for differences between GLM ALL and GCR and between GLM ALL and GLM BH and moderate evidence for differences between GLM ALL and CAR as well as between GLM ALL and SSR, reflecting the overall highest CNR for GLM ALL (**GLM ALL** > **GCR, SSR, GLM BH, and CAR**).

Regarding the Δ[HbR] data, the rmBANOVA revealed anecdotal evidence favoring the *H*_0_.

##### ME data

Detailed statistical results are reported in Tables S15 and S16 in the Supplementary Material. For Δ[HbO], the rmBANOVA revealed extreme evidence favoring the *H*_1_. Posthoc tests revealed extreme evidence for differences between GLM ALL and NO SAC, CAR as well as GCR, strong evidence for differences between GLM ALL and GLM BH, and moderate evidence for differences between GLM ALL and SSR, all resulting from higher CNR values of GLM ALL as compared with all other methods (**GLM ALL** > **CAR, GCR, SSR, GLM BH, and NO SAC**). Evidence was strong for a difference between GCR and SSR and extreme evidence was found for differences between GCR and GLM BH, with reduced CNR values for GCR (**SSR, GLM BH** > **GCR**). Moreover, evidence was strong for a smaller CNR for CAR compared with GLM BH and moderate for a smaller CNR for CAR compared with SSR (**GLM BH, SSR** > **CAR**). There was extreme evidence for differences between GLM BH and GCR, and strong evidence was found for the difference between NO SAC and GLM BH as well as CAR and GLM BH with higher CNR for GLM BH (**GLM BH** > **NO SAC**).

For Δ[HbR], the rmBANOVA indicated extreme evidence favoring the H**j**. Posthoc tests indicated that this was related to reduced CNR values for CAR compared with GLM ALL, for which strong evidence was found (**GLM ALL** > **CAR**). Similarly, there was strong evidence for differences between GCR and GLM ALL and moderate evidence between GCR and GLM BH reflecting smaller CNR values for GCR (**GLM ALL, GLM BH** > **GCR**).

##### MI data

Detailed statistical results are reported in Tables S17 and S18 in the Supplementary Material. Regarding both the Δ[HbO] and the Δ[HbR] data, the rmBANOVA revealed anecdotal evidence favoring the *H*_0_.

### 3.2 Spatial Specificity

#### 3.2.1 Correlation matrices with SDCs (SDC CORMAT)

Overall and across tasks, descriptively, the correlation matrices of Δ[HbO] for NO SAC showed a strong influence of the SA across channels, indicated by high correlation coefficients between regular channels as well as between regular channels and SDCs. The high correlations are already attenuated but not absent when considering Δ[HbR] [cf. Figs. 5(a), 6(a), and 7(a)]. For both signal types correlations with SDCs are highly reduced after applying correction methods irrespective of using SDCs for correction or not. However, negative correlations between regular channels as well as between regular channels and SDCs resulted from correction with methods CAR and GCR.

**Fig. 5.**
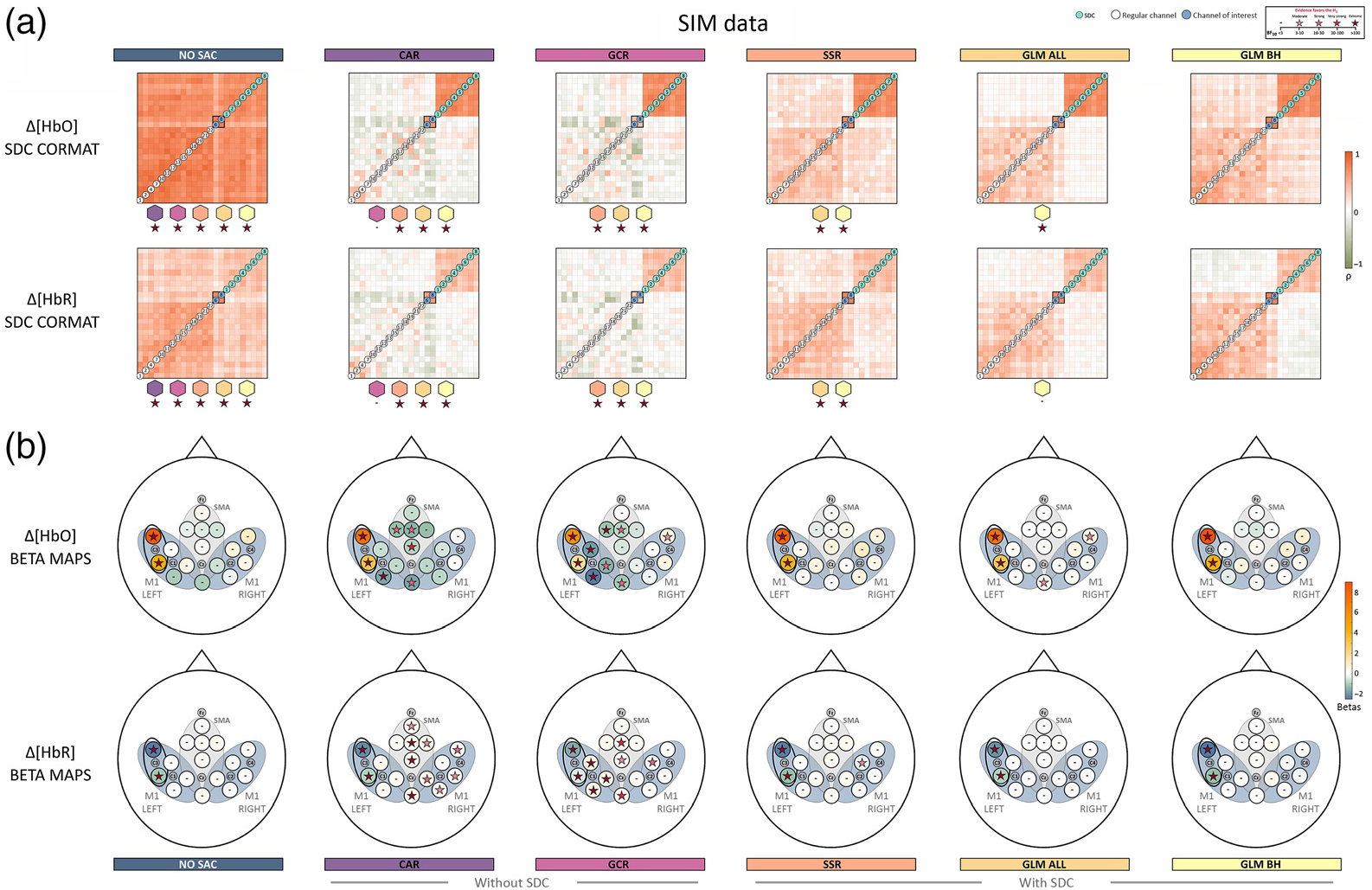
Illustration of the spatial specificity results for SIM data. (a) The averaged correlation matrices of Δ[HbO] and Δ[HbO]. Colored stars below a correlation matrix represent Bayes factors of the statistical comparison between the respective matrix and the matrix belonging to the correction method that is indicated by the colored polygon above the Bayes factor. Note that the order of the regular channels in the matrix is arbitrarily chosen in that the order always starts in the lower left corner with channels outside the ROI (in ascending order, indicated by white circles) and are followed by channels inside the ROI (in ascending order, indicated by gray circles and corresponding to ROI M1 left). (b) Visualizes the topographical beta maps (BETA MAPS) of Δ[HbO] and Δ[HbR]. Each circle represents a channel with its corresponding mean beta value. The star within a circle represents the Bayes factor resulting from the Bayesian *t*-test of the beta values of the respective channel. The black oval marks the ROI, M1 LEFT.

**Fig.6.**
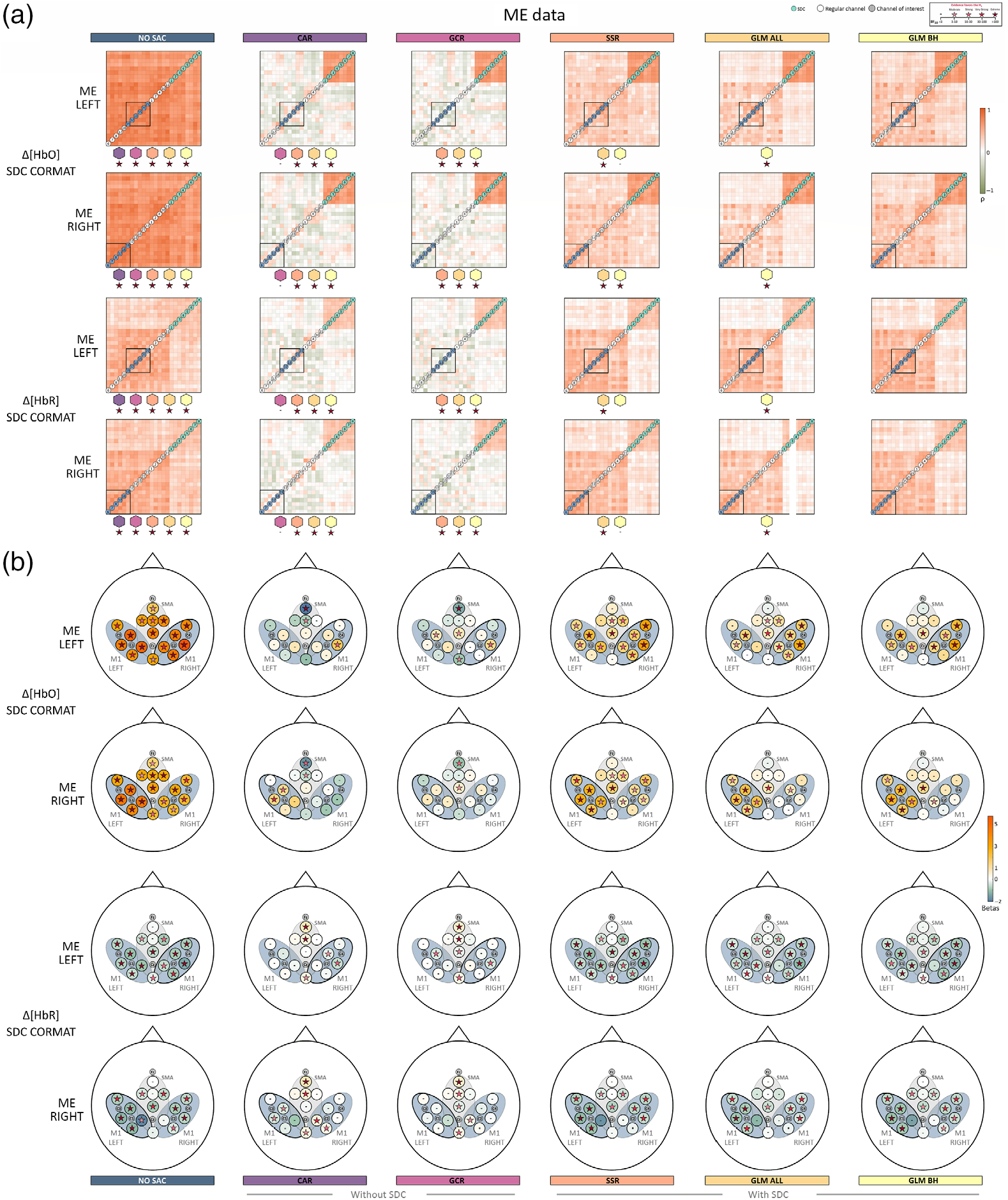
Illustration of the spatial specificity results of ME LEFT and ME RIGHT data. (a) The averaged correlation matrices of Δ[HbO] and Δ[HbR]. Colored stars below a correlation matrix represent Bayes factors of the statistical comparison between the respective matrix and the matrix belonging to the correction method that is indicated by the colored polygon above the Bayes factor. Note that the order of the regular channels in the matrix is arbitrarily chosen in that the order always starts in the lower left corner with channels outside the ROI (in ascending order, indicated by white circles) and are followed by channels inside the ROI (in ascending order, indicated by gray circles and corresponding to ROI M1 left). (b) shows the topographical beta maps (BETA MAPS) of Δ[HbO] and Δ[HbR]. Each circle represents a channel with its corresponding mean beta value. The star within a circle represents the Bayes factor resulting from the Bayesian *t*-test of the beta values of the respective channel. The black ovals mark the ROIs, M1 LEFT and RIGHT.

**Fig. 7.**
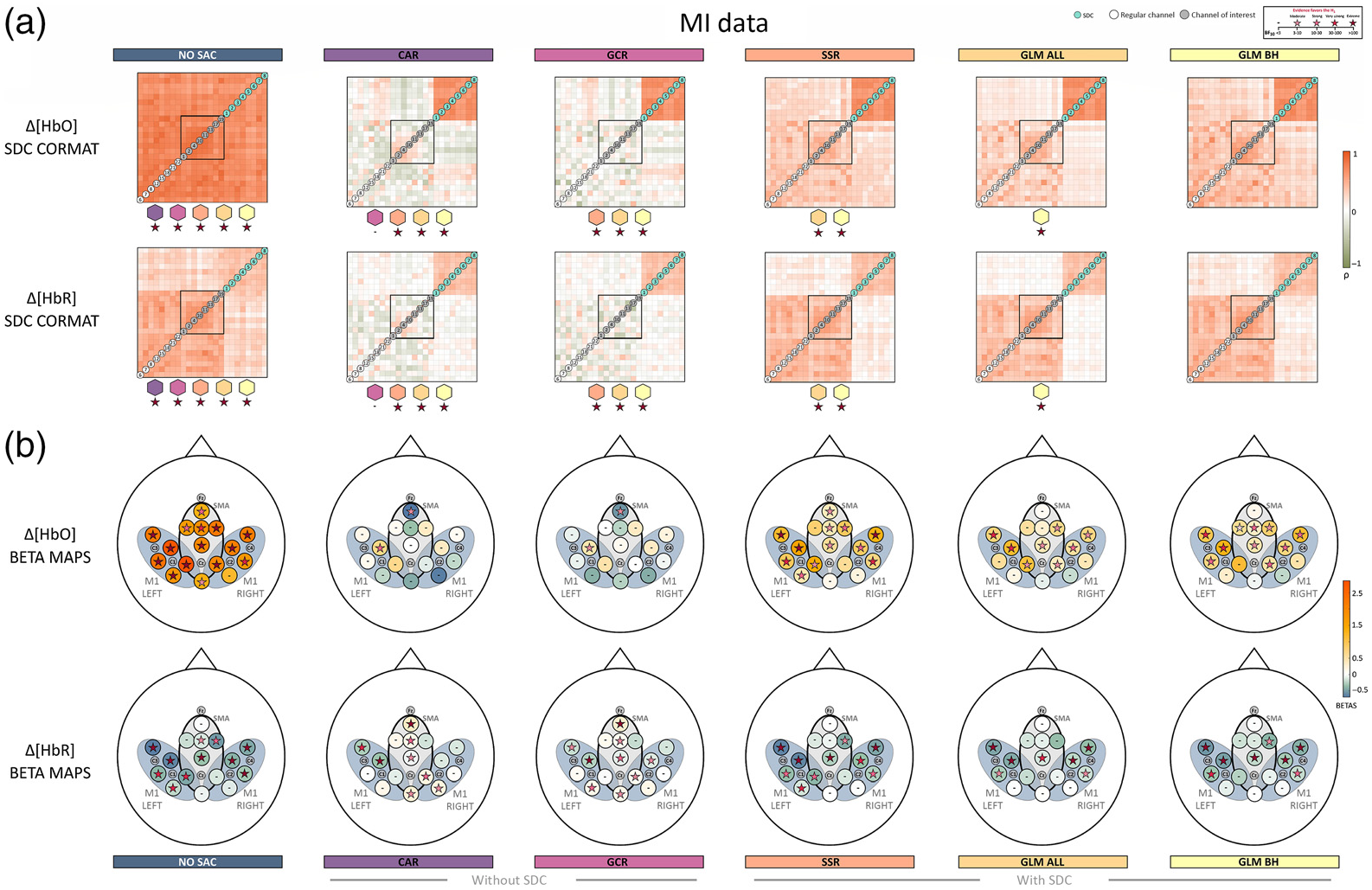
Illustration of the spatial specificity results for SIM data. (a) The averaged correlation matrices of Δ[HbO] and Δ[HbR]. Colored stars below a correlation matrix represent Bayes factors of the statistical comparison between the respective matrix and the matrix belonging to the correction method that is indicated by the colored polygon above the Bayes factor. Note that the order of the regular channels in the matrix is arbitrarily chosen in that the order always starts in the lower left corner with channels outside the ROI (in ascending order, indicated by white circles) and are followed by channels inside the ROI (in ascending order, indicated by gray circles and corresponding to ROI M1 left). (b) The topographical beta maps (BETA MAPS) of Δ[HbO] and Δ[HbR]. Each circle represents a channel with its corresponding mean beta value. The star within a circle represents the Bayes factor resulting from the Bayesian *t*-test of the beta values of the respective channel. The black oval marks the ROI, SMA.

**Fig. 8.**
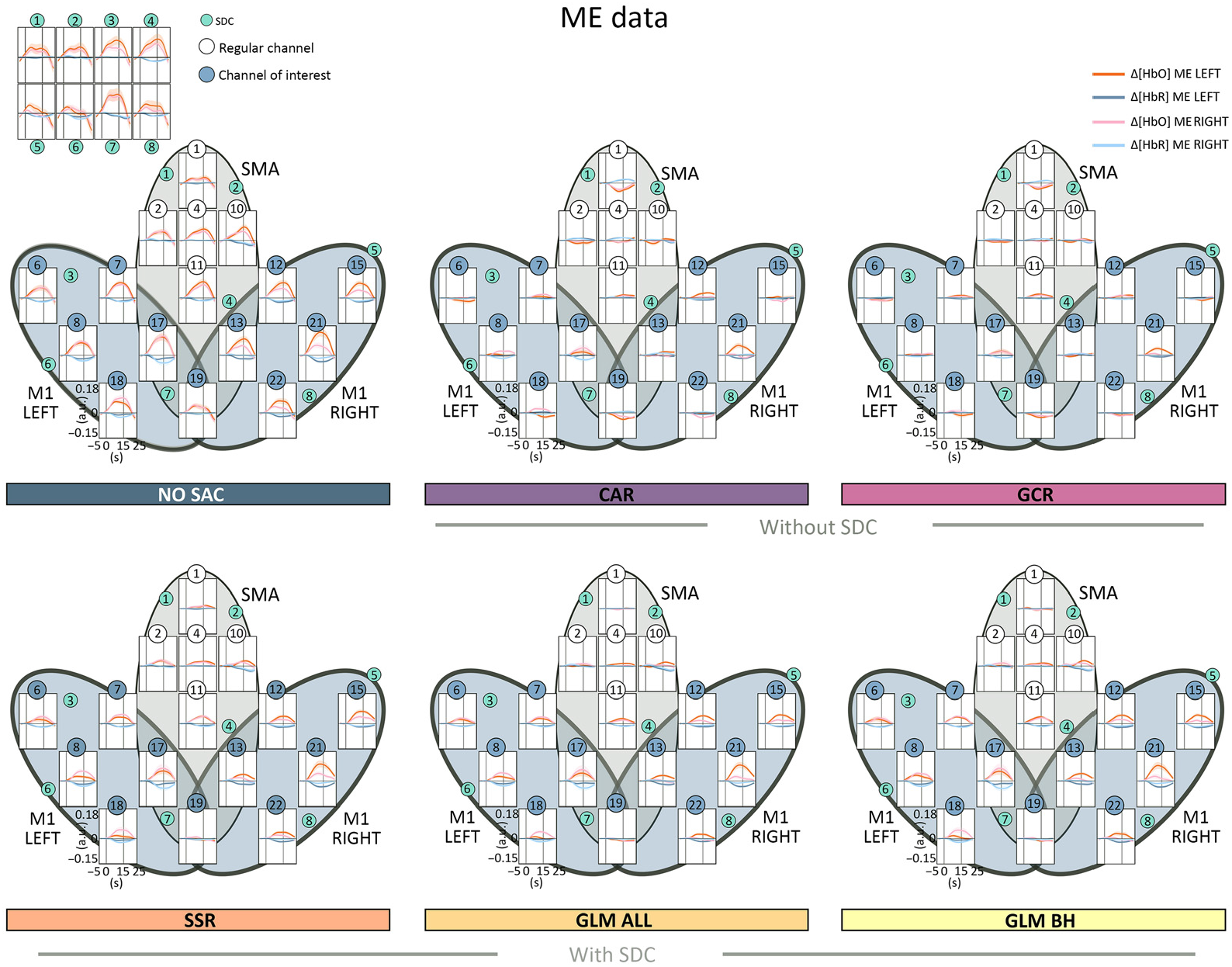
Illustration of the averaged time series data of both Δ[HbO] and Δ[HbR] including all channels for all correction methods. For method NO SAC, additionally, the averaged time series data of the SDCs is visualized (top left panel). The black ovals mark the ROI, M1 LEFT and RIGHT.

Statistical analyses were run with Bayesian paired t-tests, performed on the lower triangular part of the correlation matrices for any two pairs of correction methods. The results of all t-tests are summarized in Tables S19 and S20 in the Supplementary Material.

##### SIM data

As visualized in Fig. 5(a), for both Δ[HbO] and Δ[HbR], nearly all SDC CORMAT comparisons indicated extreme evidence for differences between correction methods. Exceptions were comparisons between CAR and GCR with respect to Δ[HbO] and Δ[HbR] as well as between methods GLM ALL and GLM BH for the Δ[HbR] data.

##### ME data

For both ME LEFT and ME RIGHT [cf. Fig. 6(a)], results indicated extreme evidence for differences between correlation matrices. Exceptions were the comparisons between CAR and GCR for both Δ[HbO] and Δ[HbR] of ME LEFT and ME RIGHT and between SSR and GLM BH for both Δ[HbO] and Δ[HbR] of ME LEFT and for Δ[HbR] of ME RIGHT.

##### MI data

As illustrated in Fig. 7(a), all comparisons between correlation matrices resulted in extreme evidence for differences. Sole exception was the comparison between CAR and GCR. This pattern was identical for Δ[HbO] and Δ[HbR].

#### 3.2.2 Topographical beta maps (BETA MAPS)

The beta maps for each method are visualized for SIM in Fig. 5(b), ME in Fig. 6(b), and MI in Fig. 7(b). As expected, descriptively, the beta maps of the SIM data showed the strongest activation in channel 6 (100% amplitude) directly followed by channel 8 (50% amplitude) across methods and data types. For the real data, visual inspection shows all SAC methods differ clearly from the NO SAC data for which beta values indicated comparable activation across channels. For instance, a clearer lateralization in the ME beta maps can be seen after correction [cf. Fig. 6(b)]. In addition, clear differences between SAC methods with and without SDCs are visible across tasks. That is, for SAC methods without SDCs but not for SAC methods with SDCs beta maps show either unexpectedly negative (Δ[HbO]) or positive (Δ[HbR]) beta values for some channels.

##### SIM data

With respect to the Δ[HbO] BETA MAPS [cf. Fig. 5(b) top], for **all methods** the one sample Bayesian t-tests of channels 6 and 8 revealed extreme evidence for differing positively from zero. This was expected because these channels contain the simulated hemodynamic response. Two additional channels differed positively from zero for method **GLM ALL**. For methods, **CAR** and **GCR** one sample Bayesian t-tests indicated at least moderate evidence for a difference from zero at five and seven additional channels, respectively. Differences were mostly but not exclusively due to negative beta values (cf. Table 21 in the Supplementary Material for exact values).

Regarding the Δ[HbR] BETA MAPS [cf. Fig. 5(b) bottom], results indicated as expected extreme evidence for a negative difference from zero for data of simulated channels 6 and 8 of all methods. Only methods **CAR**, **GCR**, and **SSR** resulted in additional channels mostly with evidence for positive differences from zero (cf. Table 22 in the Supplementary Material for exact values).

##### ME data

Regarding the Δ[HbO] data of both ME LEFT and ME RIGHT [cf. Fig. 6(b) top two rows], BETA MAPS for method **NO SAC** showed moderate to extreme evidence for differing positively from zero across all channels (16/16). For **CAR** one sample Bayesian t-tests indicated that a small number of channels differed positively (ME LEFT: 1/6 in ROI M1 RIGHT; ME RIGHT: 2/6 in ROI M1 LEFT) or for negatively (both ME LEFT and ME RIGHT: 2/16 in ROI SMA) from zero. Results were similar for **GCR** (positive differences ME LEFT: 3/16 in all ROIs, 1/16 in M1 RIGHT; ME RIGHT: 1/16 in ROI SMA; negative differences ME LEFT: 2/16 in all ROIs; ME RIGHT: 1/16 in ROI SMA). Statistical analysis of BETA MAPS of method **SSR** showed for most channels moderate to extreme evidence for differing positively from zero (ME LEFT: 12/16 in all ROIs, 5/6 in M1 RIGHT; ME RIGHT: 12/16 in all ROIs, 5/6 in M1 LEFT). BETA MAPS for methods **GLM ALL** and **GLM BH** differed with moderate to extreme evidence positively from zero in a medium number of channels (**GLM ALL**: ME LEFT: 10/16 in all ROIs, 5/6 in M1 RIGHT; ME RIGHT: 7/16 in all ROIs, 5/6 in M1 LEFT; **GLM BH**: ME LEFT: 8/16 in all ROIs, 4/6 in M1 RIGHT; ME RIGHT: 7/16 in all ROIs, 5/6 in M1 LEFT) (cf. Table S23 in the Supplementary Material for exact values).

For Δ[HbR] of **NO SAC**, for ME LEFT data 14 of 16, and for ME RIGHT data of 12 of 16 channels provided evidence for moderate to extreme evidence for negatively differing from zero as indicated by the results of one sample Bayesian t-tests [cf. Fig. 6(b) bottom two rows]. Statistical results for methods **CAR** and **GCR** revealed in more channels evidence for differing positively from zero (**CAR**: ME LEFT: 3/16 in all ROIs; ME RIGHT: 5/16 in all ROIs; **GCR**: ME LEFT: 3/7 in all ROIs; ME RIGHT: 4/16 in all ROIs) than for differing negatively from zero (**CAR**: ME LEFT: 1/6 in ROI M1 RIGHT; ME RIGHT: 0/16; **GCR**: ME LEFT: 3/16 in all ROIs; ME RIGHT: 1/8 in ROI SMA), which would however be the expected direction. Results for **SSR** showed moderate to extreme evidence for most channels to differ negatively from zero (ME LEFT: 14/16 in all ROIs, 6/6 in M1 RIGHT; ME RIGHT: 10/16 in all ROIs, 4/6 in M1 LEFT). For **GLM ALL** and **GLM BH**, the number of channels with evidence for differing negatively from zero was similar or slightly reduced compared with SSR (**GLM ALL**: ME LEFT: 13/16 in all ROIs, 5/6 in M1 RIGHT; ME RIGHT: 8/16 in all ROIs, 4/6 in M1 LEFT; **GLM BH**: ME LEFT: 13/16 in all ROIs, 5/6 in M1 RIGHT; ME RIGHT: 11/16 in all ROIs, 4/6 in M1 LEFT). For more details see Table S24 in the Supplementary Material.

##### MI data

Regarding the Δ[HbO] data of **NO SAC**, data of 15 out of 16 channels across all ROIs provided moderate to extreme evidence for differing positively from zero [cf. Fig. 7(b) top]. Regarding **CAR** and **GCR**, there was evidence for one channel to differ positively (ROI M1 LEFT) and for one channel to differ negatively (SMA) from zero. For **SSR**, 12 out of 16 channels across all ROIs differed positively from zero with moderate to extreme evidence. For **GLM ALL**, data of 8 out of 16 and for **GLM BH** 10 out of 16 channels across all ROIs provided moderate to very strong evidence for differing positively from zero (cf. Table S25 in the Supplementary Material for exact values).

With respect to Δ[HbR], one sample Bayesian t-tests provided moderate to extreme evidence for beta values to differ negatively from zero in 11 out of 16 channels across all ROIs for **NO SAC** [cf. Fig. 7(b) bottom]. For method **CAR**, for 5 out of 16 channels across all ROIs evidence was at least moderate for a positive difference from zero, whereas for three further channels evidence was at least moderate for a negative difference (ROIs M1 LEFT and M1 RIGHT). For **GCR**, there was evidence for differing positively from zero for 5 out of 16 channels across all ROIs and for differing negatively from zero for 4 out of 16 channels across all ROIs. Method **SSR** was linked to evidence indicating a negative difference from zero for 10 out of 16 channels across all ROIs. For **GLM ALL**, this was only the case for 7 and for **GLM BH** for 8 out of 16 channels and across all ROIs (cf. Table S26 in the Supplementary Material for exact values).

## 4 Discussion

The present study aimed at comparing SA correction (SAC) methods with (SSR, GLM ALL, and GLM BH) and without (CAR and GCR) short distance channels (SDCs) as well as preprocessed but otherwise uncorrected data (NO SAC) on the single trial level. SAC methods were applied to semisimulated (SIM) and real (ME and MI) data of healthy older adults. Quality measures quantified signal improvement and spatial specificity associated with SAC correction for both Δ[HbO] and Δ[HbR].

### 4.1 Signal Improvement

Signal improvement was assessed on a single trial basis using scaled root mean squared error (sRMSE), Spearman correlation (COR), and contrast-to-noise ratio (CNR). Across all tasks and quality measures, we observed that the (uncorrected) Δ[HbO] data is more affected by SA as compare with the (uncorrected) Δ [HbR] data. In addition, both semisimulated and real data the quality measures changed for Δ [HbR] after correction, which confirms that Δ [HbR] is influenced by SA too and should thereby be corrected accordingly.^28^

Across all three measures, the effect of SA correction was most prominent for SIM data, which is in line with previous findings.^13^ Of the SAC methods applied, GLM ALL outperformed all other approaches for SIM data. For the real data, descriptively, GLM ALL also tended to rank first, however, statistically this could only be confirmed for a subset of comparisons. Moreover, for the real data but not the SIM data, methods without SDCs showed in most cases a signal decline rather than an improvement. Potential reasons for the varying results between SIM data and the real data might be the different signal-to-noise ratios (SNR) and the lacking task-related activation in the SDCs of the SIM data.^13^ The latter might have resulted in an overestimation of the effect of the correction methods for the SIM data, because here, in contrast to real data, the SA measured at SDCs and the synthetic canonical HRF are virtually uncorrelated. Moreover, quality measures sRMSE and COR were based on the assumption that the underlying hemodynamic response in the signal corresponds to a canonical HRF peaking at 6s after stimulus onset. This assumption is met perfectly for SIM data, but, in all likelihood, to a much smaller degree for the real data. This likely resulted in reduced sensitivity of measures sRMSE and COR for the assessment of signal improvement in real data. To improve the sensitivity of HRF-based quality measures a GLM-based deconvolution could be applied in future studies to estimate the individual unknown task-related HRF.^55,56^ The individually estimated HRF could then be used for quality measures sRMSE and COR. In general, however, it would be beneficial to have quality measures assessing signal improvement that have less assumptions regarding shape and timing properties of the underlying signal. In the present study, this class of quality measure was represented by the CNR, in which the ratio of the baseline and the peak of a given trial is evaluated independent of the general shape or the peak latency. Across the quality, measures quantifying signal improvement COR and CNR seemed indeed to be the most sensitive quality measure as they picked up the expected differences in signal improvement also for real data. The CNR is, however, also not without problems, as one of its defining features reflects the activation strength resulting from a task or event. This could, e.g., have contributed to the larger CNR differences between corrected and uncorrected data for ME DATA compared with MI DATA [cf. Figs. 4(a) and 4(b)], in spite of the underlying data set being the same.

To sum up, single trial signal improvement was clearest and most consistent for the SIM data. It was generally higher for methods with SDCs as compared with methods without SDCs and as compared with the uncorrected data. The overall best performing method was GLM ALL. However, neither signal improvement quality measure seems suitable to exclusively quantify signal improvement alone, because measures are either dependent on potentially inaccurate assumptions regarding the shape of the HRF or because they are influenced by the magnitude of the activation resulting from a task or event. For the given real data set, however, CNR seems to be the most sensitive quality measure followed by COR. In contrast, the sRMSE quality measure was not able to reliably reflect the differences between correction methods.

### 4.2 Spatial Specificity

Spatial specificity was investigated by means of Spearman correlation matrices (SDC CORMAT) at the single trial level and by considering topographical beta maps (BETA MAPS) resulting from GLM analyses based on the whole time series. Spatial specificity measures were primarily intended to be used as descriptive measures, statistical comparisons were performed only complementary.

Overall, uncorrected (NO SAC) Δ[HbO] and Δ[HbR] data were clearly affected by SA. However, spatial specificity of Δ[HbO] data was already much lower as compared with Δ[HbR] data as was evident from very high correlations between regular channels as well as between regular channels and SDCs (SDC CORMAT). Also, for NO SAC Δ[HbO], and here in particular for real data, beta activation was strong across nearly all regular channels (cf. Figs. 5(b), 6(b) and 7(b)). Across all tasks and data types, correlations between regular channels and SDCs were highly reduced for all applied correction methods. Likewise, beta activation revealed in most cases a more defined spatial pattern following SAC correction. For both spatial specificity measures, methods without SDCs (CAR and GCR) resulted in negative correlations and negative activation. This pattern is highly suggestive of an overcorrection resulting from these methods, which is also a problem for other methods without SDCs, e.g., ICA-based methods.^55^

Summarized, both SDC CORMAT and BETA MAPS seem well suited for picking up improvements in spatial specificity following SA correction but also for spotting potential over-correction. A drawback is that not all the differences between matrices and maps that are easily spotted by the eye are suitable for statistical testing. Moreover, a single trial equivalent of the BETA MAPS measure would be useful for assessing changes in spatial specificity at the single trial level, at which NFB and BCIs work.

### 4.3 Correction Methods

#### 4.3.1 Common average reference (CAR)

The CAR method^22^ is a spatial filter for which in the present study the averaged signal across all regular channels was subtracted from each individual channel. Of the two methods without SDCs, CAR resulted in a greater signal improvement. Surprisingly, for the SIM data, signal improvement following CAR was larger as compared with SSR and GLM BH across all three quality measures. However, for the real data CAR seems to overcorrect the data. Overcorrection was also suggested by spatial specificity measures with negative correlations (SDC CORMAT) and negative beta values (BETA MAPS) across all tasks. On the other hand, channels indicated as active by BETA MAPS were comparable to the methods with SDCs, as for instance reflected in the lateralization pattern for the ME DATA [cf. Fig. 6(b)]. This indicates a good spatial specificity of CAR for strongly activating tasks.

The negativity in the signal following CAR is not surprising given that strongly activated channels affect the amplitude of the averaged signal and channels with no or lower amplitudes get pushed into the reversed direction. A workaround would be to omit the channels in the critical ROI from the averaged signal. However, this only works if the ROI covers the peak activation and when channels outside of the ROI are not expected to be active. This is rather unlikely for any but purely perceptual tasks, and even here it is doubtful. A more advanced version of this method could be to regress out the average across all channels, e.g., by using the SSR^19^ method,^15^ instead of simply subtracting it. Moreover, for the implemented version of the CAR method or any modification thereof, a proper channel quality check and channel pruning are important in order to avoid to subtract data of poor quality into a given channel.

Overall, although it can result in artificial negativity of inactive channels, CAR seems to be a good and easy-to-implement choice if no SDCs are available and if only data of one channel or a very circumscribed ROI covering the peak activation will be used for (online) processing.

#### 4.3.2 Global component removal (GCR)

The GCR method^23^ is a spatial filter that combines a Gaussian kernel smoothing by taking into account the MNE coordinates of the channel positions and a singular value decomposition. Regarding the signal improvement measures, for SIM the results indicated a signal improvement across all quality measures. For the real data the opposite was evident, that is, results indicated a signal decline. Overcorrection was indicated by strong negative correlations between regular channels and between regular channels and SDCs, and by negative beta activation across tasks and data types. The reason for the artificial negative activation is probably the combination of the relatively low spatial coverage and the large-sized kernel.^23^ Although the probe in the present study falls into the recommended minimum area^23^ it seems that this method needs a much higher head coverage. In a recent study,^24^ it was shown that higher-density data corrected by means of the GCR was comparable with that corrected by a regression-based approach using SDCs. However, this approach considered only the first two principal components^24^ resulting from a principal component analysis (PCA) of the SDC data and therefore missed potentially important spatial information that could have made a difference to the results.

On the whole, the GCR method performed well for SIM data but not for the real data. Therefore, it should be used with care and should only be applied with sufficiently high spatial coverage to avoid overcorrection. To decide whether the number of channels is sufficient, a comparison of the beta output of a GLM (as done for BETA MAPS) before and after correction might be helpful.

#### 4.3.3 Short separation regression (SSR)

The SSR method^19^ is a simple regression method. In the present study, we applied SSR-based correction using the spatially closest SDC to a given regular channel. For both signal improvement and spatial specificity measures, out of the correction methods with SDCs the SSR performed poorest and in some cases performed worse than methods without SDCs. Moreover, for real data, the spatial specificity measures suggested a stronger impact of SA after correction compared with the two GLM methods (cf. Figs. 6–9). One explanation for this limited performance might be that the closest SDC is not necessarily the best choice for SSR. Alternatively, the SDC with the highest correlation, with the average across SDCs or with the first *n* principal components resulting from a PCA based on all SDCs^24^ could be used.

**Fig. 9.**
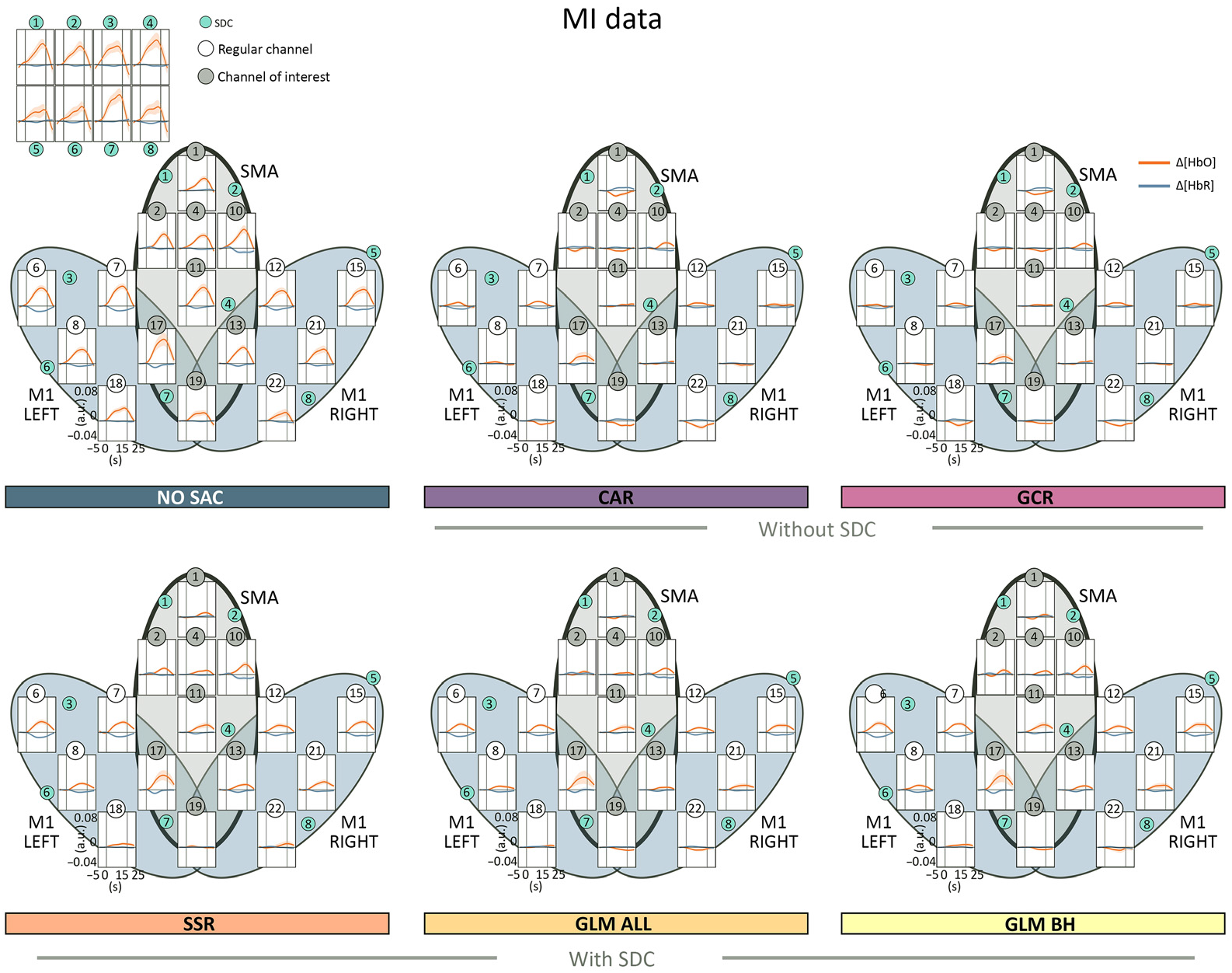
Illustration of the averaged time series data of both Δ[HbO] and Δ[HbR] including all channels for all correction methods. For method NO SAC, additionally, the averaged time series data of the SDCs is visualized (top left panel). The black oval marks the ROI, SMA.

**Fig. 10.**
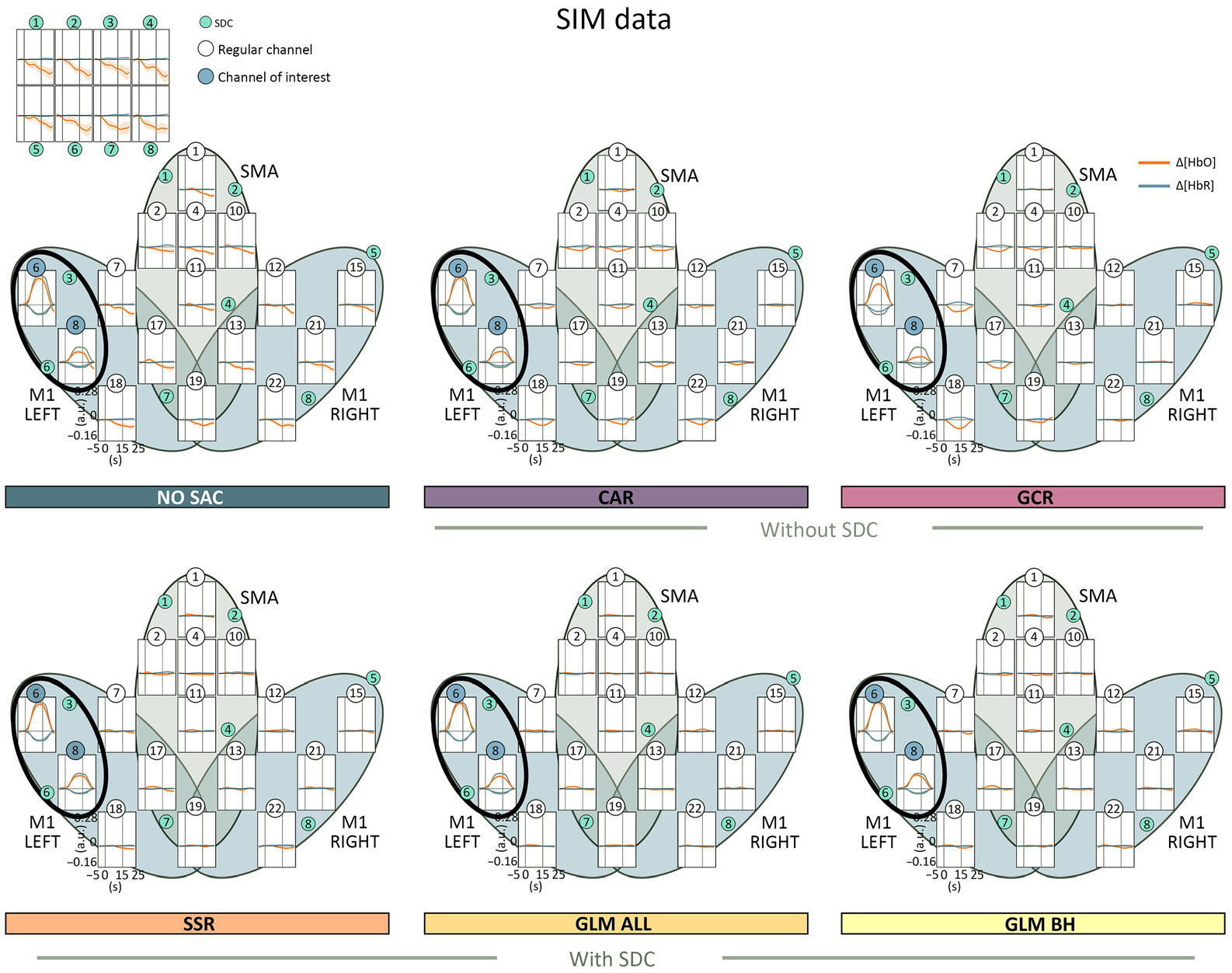
Illustration of the averaged time series data of both Δ[HbO] and Δ[HbR] including all channels for all correction methods. For method NO SAC, additionally, the averaged time series data of the SDCs is visualized (top left panel). The black oval marks the ROI, channels 6 and 8 of M1 LEFT.

In spite of these shortcomings, given the simplicity of the SSR method especially regarding online implementation, it can still be recommended. However, one important step before applying this method, especially when considering the closest SDC for correction, is the signal quality check. If SDCs or regular channels of poor signal quality are not properly pruned before correction, regular channels might either be corrupted, remain uncorrected or SA will get amplified.

#### 4.3.4 General linear model with all SDCs as regressors (GLM ALL)

In the present study, of the selected SAC methods, the GLM ALL method^13,17^ was overall performing best. For this correction method, all eight SDCs of both Δ[HbO] and Δ[HbR] were included as regressors to the SAC GLM, because previous work provided evidence that the SA is not a homogeneous but rather a heterogeneous artifact^13,57^ and therefore, a higher number of SDCs covering a large portion of the head improves correction.^17^

That the GLM ALL method seems to be the most accurate method for SA correction is in line with previous findings.^13,17^ Although the AR-ILS^36^ GLM is generally computational more intensive as compared with the ordinary least squares (OLS) GLM, its feasibility for online application has been shown^58^ and it has been implemented in online preprocessing tools such as TurboSatori.^59^ Moreover, AR-ILS can handle the presence of motion artifacts very well.^36^ This is, however, not the case for the other SA correction methods and hence, could have been a gain for the GLM-based approaches. However, if motion artifacts were present in our data sets then usually at the beginning/end of the recording or during the breaks and only in very rare cases during the task blocks. Hence, even if some motion artifacts survived the TDDR correction, the advantage of the AR-ILS GLM over the other correction methods should be low.

In spite of the good performance of GLM ALL, it is likely that results can still be improved by adding more information to the GLM. For instance, gains can be expected by adding additional regressors based on auxiliary recorded physiological measures like blood pressure and oxygen saturation,^55^ information extracted from the fNIRS measurement itself like Mayer waves,^13^ or a global signal regressor generated from the SDCs.^13^ Another promising way would be to increase the number of SDCs as it was shown that the number of SDCs has a strong effect on the correction performance of the GLM, i.e., the more information in terms of SDCs is available the better the correction performance.^17^ Another promising approach would be to cover blind spots by, e.g., adding short separation sources to the detectors the same way as for the available hardware solutions short separation detectors are added to the sources.^13^ Moreover, in the present study for method GLM ALL but also for method GLM BH, two GLMs were applied consecutively. That is, first a GLM was applied only for SA correction by including all SDCs as regressors. From this GLM, SAC corrected data was extracted for further evaluation, among others based on a second GLM in which task-induced activation was modeled. This two-step procedure was chosen because it provided the SAC corrected data required as input for all quality measures. In principle, if only the task-related activation is of interest, it would also be possible to run a single GLM with both the SDC regressors and the task-related regressors. Indeed, for BCI and NFB online scenarios in which the feedback is based on the GLM output this would seem like the natural choice. One major issue with this one-step approach is, however, the risk of multicollinearity due to the strong task-evoked activity in the SDCs, which can lead to unpredictable results.^17^ To prevent this problem a two-step GLM approach as applied in the present study should be considered, even if the main focus of the GLM analysis is the extraction of task-related activation.

In sum, the GLM ALL method strongly increased both signal quality and spatial specificity in the present data sets. Accordingly, of the evaluated methods, it is the best choice for SA correction if SDCs are available. Within a real-time GLM calculation, it can be easily applied for online SA correction either by implementing it independently or by using available toolboxes.^59^

#### 4.3.5 General linear model with one SDC on both hemispheres as regressors (GLM BH)

For the GLM BH method only two out of the eight available SDCs (SDC 4 and 7 of both Δ[HbO] and Δ[HbR]) were included in the GLM for correction. The intention was to evaluate GLM correction performance for constellations in which only a small number of SDCs are available. GLM BH produced overall satisfying results that were, however, inferior to the results from GLM ALL. The finding of overall inferiority to a GLM relying on larger numbers of SDCs is in line with previous findings.^17^ On closer inspection, results for GLM BH were comparable to those of the SSR method. For spatial specificity, however, GLM BH results were, descriptively, more comparable to GLM ALL. As a workaround for unavailable SDCs, some researchers help themselves by sacrificing one or two regular channels and placing the vacated sources and detectors as close as possible to each other to create a self-made SDC.^41,42^ This approach should be considered with care because depending on the manufacturer the resulting distance between source and detector is likely ≥10 mm. SDCs placed at distances ≥10 mm may capture different signals than SDCs placed at the recommended 8 mm as used in the present study because of NIR light penetrating to cortical brain tissue.^18^ This will likely influence the recovery of the true brain signal and bears the risk of overcorrection.^18^ This is, however, similarly true for all correction methods using SDCs, regardless of the number of SDCs used. Moreover, if not handled correctly, e.g., by the recording software, self-made SDCs can possibly lead to a saturation of the self-made SDCs. Nevertheless, if properly used, self-made SDCs are should still be better than using no SDCs.

In sum, with a reduced set of 8-mm SDCs, GLM BH achieved satisfying results for GLM-based correction, but results were somewhat inferior to GLM ALL. Similar to GLM-ALL, this method can easily be implemented online as part of a real-time GLM calculation.

#### 4.3.6 Recommendations

Based on the present results and in line with previous findings,^13,17^ the best performing SA correction method is the GLM ALL method directly followed by the GLM BH and the SSR method. For the GLM ALL method all eight SDCs were used as regressors, for the GLM BH method two predefined SDCs were used and for SSR only the spatially closest SDC was used for correction. These results indicate that correction performance increases with the number of SDCs used for the correction, which is in line with findings of Santosa and colleagues^17^ who investigated the effect of the number of SDCs for a GLM-based correction approach. In the study by Santosa et al.^17^ the maximum number of available SDCs was likewise eight, however, the authors predicted that the performance will likely increase with more SDCs. The methods without SDCs showed a lower performance compared with the methods with SDCs and tended to overcorrect the data. However, based on the performance results of CAR and GCR and given the strong influence of the SA artifact, the application of either of these or another correction method^17,27^ should be preferred over the option to neglect the SA correction step. Based on the present results and especially if only a limited optode coverage is available (cf. next Sec. 4.4), we would recommend the CAR method over the GCR method because the overcorrection seems slightly lower and CAR results in a higher spatial specificity as compared with the GCR method.

### 4.4 Limitations

The present study is subject to several limitations. First of all, although most of the quality measures investigated in the present study were calculated based on single trials in order to mimic an online scenario, it is highly recommended to perform a validation additionally in an online scenario^60^ in order to ensure comparable results when methods have to perform with (nearly) real-time output.

In the present study, only data from healthy older adults were analyzed. There is evidence that the hemodynamic response differs between age groups with respect to, e.g., shape^7^ and strength.^61^ Therefore, the present findings might not generalize to younger individuals. However, whether there are age-related differences regarding the SA remains to be shown.

One limitation regarding the SIM data is that it was based on a relatively short rest recording lasting ~3 min, allowing the simulation of only five block-designed trials. For the same purpose, previous studies used for the same purpose longer periods of rest data ranging from 5^17^ to 15 min^13^ and could thus create a higher number of semisimulated trials. A higher number of trials usually increases the reliability of the signal of interest. However, given that the SNR for semisimulated data is generally somewhat higher as compared with real data, also a low number of trials should be acceptable for the purpose of comparing the performance of correction methods.

Moreover, and more importantly, no task-evoked activation was added to the SDCs for the SIM data, though previous studies^62,63^ implicate that the SA captured by the SDCs also contains a task-evoked component. This was also evident in the present study as can be seen in all figures depicting SDC and regular channel activation: In figures showing group-level real data (cf. Figs. 8 and 9), task-related SA mimicking brain activation is evident in almost all SDCs. The task-related nature of the SDC signal becomes particularly obvious when comparing SDCs of SIM data to those resulting from the real data set (cf. Figs. 8–10). Moreover, single trial data (cf. single trial and single subject data of selected channels as well as SDCs in figures of the Supplementary Material) demonstrate not only that most SDCs are affected by task-evoked SA, but also that the artifact is quite diverse: SA differs between SDCs (heterogeneity^13^) within the same trial, between subjects, between signal types^14^ Δ[HbO] and Δ[HbR]), and between tasks.^14^ Moreover, it is not necessarily the closest SDC that shows the highest similarity with the uncorrected data. Perhaps due to this diversity, so far, there exist no recommendations on how to best semisimulate task-evoked SA in SDCs, and hence, studies on simulated data restrict simulated evoked activity to regular channels.^13,21^ For making results from SDC correction of semisimulated data more realistic, however, semisimulating task-evoked SA is a highly desirable next step. Important impulses for it could come from a systematic collection of information on, e.g., the spatial distribution of task-evoked SA across the head, its peak latency as compared with a regular channel, amplitude characteristics of task-evoked SA, or differences between Δ [HbO] and Δ[HbR].

For the present study, the effect of correction method was evaluated for both Δ[HbO] and Δ[HbR] data, following the most recent recommendations in the field.^56^ Beyond that, it would also be possible to evaluate performance for total hemoglobin concentration changes^8^ (i.e., Δ[HbR] = Δ[HbO] + Δ[HbR]) and the difference between types of concentration changes^64^ (i.e., Δ[HbDiff] = Δ[HbO] — Δ[HbR]), both data types are however rarely reported in current publications.

One more specific limitation of the present study concerns the GCR method. More precisely, for this correction method MNI coordinates are used for the smoothing process and therefore, it would be recommended to use individual coordinates of the channels.^23^ However, since much of the digitizer data of the SIM data set were corrupted we decided to use uniform optode locations estimated by means of the ICBM-152 head model (ICBM 152 Nonlinear atlases version 2009) in the NIRSite toolbox (v2020.7; NIRx Medizintechnik GmbH, Berlin, Germany), positioned according to the International 10-20 system. For reasons of unity, the same procedure was applied to the real data. While individual optode positions might of course be more accurate, for the GCR method the distances between channels are of importance. Very likely, the distances would be similar to those of the common optode locations because all optodes were attached to a flexible cap with slits generated according to the 10-20 system. Another issue with respect to the GCR method concerns the optode coverage. Zhang and colleagues^23^ recommended to cover an area of at least 9 cm^2^, otherwise, the correction might result in artificial negative activity.^23^ Although we cover >9 cm^2^ with our optode layout, we saw in our data a substantial amount of channels resulting in reversed amplitudes.

## 5 Conclusions

This study aimed to evaluate SA correction methods with and without SDCs with regard to signal improvement and spatial specificity on a single trial level. The single trial level was chosen to simulate the application of the correction methods in a real-time online scenario, such as NFB or BCIs. Although methods with SDCs did result in a more accurate SA correction for both Δ[HbO] and Δ[HbR] as compared with methods without SDCs, our study confirms that without SDCs some improvement can be achieved too. Therefore, SA correction should become the norm for any preprocessing pipeline.^12,13^

## Acknowledgments

We would like to thank Merle Marek and Hatice Sahin for helping with the data collection of the ME/MI data set. The project was funded by a grant from the Oldenburg School of Medicine and Healthcare Sciences (2017–2010).

## Contributions

FK, CK, ML, and AB-A conceptualized the study. PR collected the data set of which the rest data was deployed. FK analyzed the data, conducted the statistical analyses, generated the figures, and wrote the first draft of the manuscript. All authors reviewed the manuscript.

## Data, Materials, and Code Availability

Data sets are not publicly available due to data protection issues. Code and materials can be shared by the corresponding author upon request.

**Franziska Klein** has a background in math and physics as well as neurocognitive psychology. She joined the neuropsychology lab as well as the Neurocognition and Functional Neurorehabilitation (NeuroCFN) group at the University of Oldenburg in 2016 as a Master’s student and continued there 2018 as a PhD student. The focus of her research lies in the development of a fNIRS-guided neurofeedback for motor neurorehabilitation and in validating and advancing signal processing techniques.

**Michael Lührs** studied computational visualistics at the Otto-von-Guericke-University in Magdeburg (2007–2012). Afterwards he began his work as a PhD student at the Department of Cognitive Neuroscience, Maastricht University, as part of the European Community Seventh Framework Initial Training Network Project Adaptive Brain Computations (ABC), which he finished in 2018. His primary research focuses on the advancement of neurofeedback and BCI tools for fMRI and fNIRS. In addition, he works at the company Brain Innovation B.V. which build free and commercial software for neuroimaging analysis and clinical application.

**Amaia Benitez-Andonegui** studied biomedical engineering at Tecnun School of Engineering in San Sebastian (Spain), before receiving her PhD in cognitive neuroscience from Maastricht University (Netherlands) in 2021. She is currently a postdoctoral fellow in the MEG Core, at the National Institutes of Health (Maryland, United States). Her research focuses on advancing wearable neuroimaging techniques, namely OPM-based MEG and fNIRS.

**Pauline Roehn** has a background in neurocognitive psychology and joined the Neurocognition and Functional Neurorehabilitation (NeuroCFN) group during her Master’s degree in 2018. After finishing her Master’s degree, she started working as data scientist.

**Cornelia Kranczioch** has a background in psychology. She joined the Neuropsychology Lab at the University of Oldenburg in 2010, where she is now a senior researcher and heads the Neurocognition and Functional Neurorehabilitation (NeuroCFN) group. Her research focus is in (motor) cognition, motor imagery and neurofeedback, and their relevance for and application in neurorehabilitation.

