## Supplementary Material for "Performance comparison of systemic activity correction in functional near-infrared spectroscopy for methods with and without short distance channels"

*Statistical Results*

**Table 1** Descriptive Statistics of sRMSE resulting from the SIM data

| | **SIM** $\Delta[HbO]$ | | | | | |
|---|---|---|---|---|---|---|
| | NO SAC | CAR | GCR | SSR | GLM ALL | GLM BH |
| Mean | 0.198 | 0.155 | 0.157 | 0.159 | 0.138 | 0.157 |
| Std. Error of Mean | 0.010 | 0.010 | 0.009 | 0.010 | 0.009 | 0.010 |
| | **SIM** $\Delta[HbR]$ | | | | | |
| | NO SAC | CAR | GCR | SSR | GLM ALL | GLM BH |
| Mean | 0.169 | 0.153 | 0.160 | 0.163 | 0.147 | 0.155 |
| Std. Error of Mean | 0.010 | 0.009 | 0.009 | 0.009 | 0.007 | 0.009 |

**Table 2** Results of the rmBANOVAs and corresponding post hoc tests with respect to the sRMSE analysis of SIM data of both $\Delta[HbO]$ and $\Delta[HbR]$. For all post hoc tests, the prior odds $P(M)$ were set to the default of 0.26, $P(M|data)$ represents the posterior odds, $BF_{10}$ the Bayes factor and $error$ the error of the given test. $BF_{10} = 1$ indicates no evidence for neither $H_0$ nor $H_1$, $BF_{10} < 1$ indicates evidence in favor of the $H_0$, that is, that there are no differences between two methods and $BF_{10} > 1$ indicates evidence in favor of the $H_1$, that is, that there are differences between two methods. According to the guidelines of Lee and Wagenmakers (2014), Bayes factors $BF_{10}$ can be categorized in the following way: $BF_{10} < 1/100$ extreme evidence for $H_0$, $1/100 < BF_{10} < 1/30$ very strong evidence for $H_0$, $1/30 < BF_{10} < 1/10$ strong evidence for $H_0$, $1/10 < BF_{10} < 1/3$ moderate evidence for $H_0$, $1/3 < BF_{10} < 1$ anecdotal evidence for $H_0$, $1 < BBF_{10} < 3$ anecdotal evidence for $H_1$, $3 < BF_{10} < 10$ moderate evidence for $H_1$, $10 < BF_{10} < 30$ strong evidence for $H_1$, $30 < BF_{10} < 100$ very strong evidence for $H_1$, $BF_{10} > 100$ extreme evidence for $H_1$.

| | | **SIM** $\Delta[HbO]$ | | | |
|---|---|---|---|---|---|
| Models | $P(M)$ | $P(M\|data)$ | $BF_M$ | $BF_{10}$ | error % |
| Null model (incl. subject) | 0.500 | $1.590e-9$ | $1.590e-9$ | 1.000 | |
| METHOD | 0.500 | 1.000 | $6.290e+8$ | $6.290e+8$ | 0.459 |
| | | Prior Odds | Posterior Odds | $BF_{10,U}$ | error % |
| NO SAC | CAR | 0.260 | 84.329 | 324.441 | $2.719e-8$ |
| | GCR | 0.260 | 333.543 | 1283.249 | $1.773e-8$ |
| | SSR | 0.260 | 21.207 | 81.589 | $1.188e-4$ |
| | GLM ALL | 0.260 | 5792.003 | 22283.701 | $2.704e-9$ |
| | GLM BH | 0.260 | 1282.865 | 4935.597 | $3.068e-9$ |
| CAR | GCR | 0.260 | 0.063 | 0.242 | 0.023 |
| | SSR | 0.260 | 0.062 | 0.239 | 0.023 |
| | GLM ALL | 0.260 | 0.526 | 2.022 | $1.498e-6$ |
| | GLM BH | 0.260 | 0.059 | 0.228 | 0.023 |
| GCR | SSR | 0.260 | 0.058 | 0.223 | 0.022 |
| | GLM ALL | 0.260 | 1.774 | 6.823 | $1.309e-7$ |
| | GLM BH | 0.260 | 0.057 | 0.219 | 0.022 |
| SSR | GLM ALL | 0.260 | 5.514 | 21.214 | $1.287e-7$ |
| | GLM BH | 0.260 | 0.058 | 0.225 | 0.022 |
| GLM ALL | GLM BH | 0.260 | 3.229 | 12.424 | $9.193e-8$ |
| | | **SIM** $\Delta[HbR]$ | | | |
| Models | $P(M)$ | $P(M\|data)$ | $BF_M$ | $BF_{10}$ | error % |
| Null model (incl. subject) | 0.500 | 0.663 | 1.966 | 1.000 | |
| METHOD | 0.500 | 0.337 | 0.509 | 0.509 | 0.554 |

**Table 3** Descriptive Statistics of sRMSE resulting from the ME data

| | | | **ME** $\Delta[HbO]$ | | | |
|---|---|---|---|---|---|---|
| | NO SAC | CAR | GCR | SSR | GLM ALL | GLM BH |
| Mean | 0.212 | 0.227 | 0.229 | 0.215 | 0.211 | 0.214 |
| Std. Error of Mean | 0.006 | 0.007 | 0.006 | 0.008 | 0.007 | 0.007 |
| | | | **ME** $\Delta[HbR]$ | | | |
| | NO SAC | CAR | GCR | SSR | GLM ALL | GLM BH |
| Mean | 0.213 | 0.219 | 0.219 | 0.216 | 0.222 | 0.216 |
| Std. Error of Mean | 0.009 | 0.007 | 0.008 | 0.008 | 0.008 | 0.008 |

**Table 4** Results of the rmBANOVAs and corresponding post hoc tests with respect to the sRMSE analysis of ME data of both $\Delta[HbO]$ and $\Delta[HbR]$. For all post hoc tests, the prior odds $P(M)$ were set to the default of 0.26, $P(M|data)$ represents the posterior odds, $BF_{10}$ the Bayes factor and $error$ the error of the given test. $BF_{10} = 1$ indicates no evidence for neither $H_0$ nor $H_1$, $BF_{10} < 1$ indicates evidence in favor of the $H_0$, that is, that there are no differences between two methods and $BF_{10} > 1$ indicates evidence in favor of the $H_1$, that is, that there are differences between two methods. According to the guidelines of Lee and Wagenmakers (2014), Bayes factors $BF_{10}$ can be categorized in the following way: $BF_{10} < 1/100$ extreme evidence for $H_0$, $1/100 < BF_{10} < 1/30$ very strong evidence for $H_0$, $1/30 < BF_{10} < 1/10$ strong evidence for $H_0$, $1/10 < BF_{10} < 1/3$ moderate evidence for $H_0$, $1/3 < BF_{10} < 1$ anecdotal evidence for $H_0$, $1 < BBF_{10} < 3$ anecdotal evidence for $H_1$, $3 < BF_{10} < 10$ moderate evidence for $H_1$, $10 < BF_{10} < 30$ strong evidence for $H_1$, $30 < BF_{10} < 100$ very strong evidence for $H_1$, $BF_{10} > 100$ extreme evidence for $H_1$.

| | | **ME** $\Delta[HbO]$ | | | |
|---|---|---|---|---|---|
| Models | $P(M)$ | $P(M\|data)$ | BF$_M$ | BF$_{10}$ | error % |
| Null model (incl. subject) | 0.500 | 0.004 | 0.005 | 1.000 | |
| METHOD | 0.500 | 0.996 | 221.233 | 221.233 | 0.356 |
| | | Prior Odds | Posterior Odds | BF$_{10,U}$ | error % |
| NO SAC | CAR | 0.260 | 4.968 | 19.115 | $8.880e-8$ |
| | GCR | 0.260 | 4.472 | 17.205 | $8.082e-8$ |
| | SSR | 0.260 | 0.061 | 0.233 | 0.024 |
| | GLM ALL | 0.260 | 0.057 | 0.218 | 0.024 |
| | GLM BH | 0.260 | 0.062 | 0.239 | 0.024 |
| CAR | GCR | 0.260 | 0.066 | 0.253 | 0.025 |
| | SSR | 0.260 | 0.432 | 1.661 | $1.960e-6$ |
| | GLM ALL | 0.260 | 30.016 | 115.482 | $1.935e-4$ |
| | GLM BH | 0.260 | 3.148 | 12.111 | $7.139e-8$ |
| GCR | SSR | 0.260 | 0.455 | 1.750 | $1.824e-6$ |
| | GLM ALL | 0.260 | 11.200 | 43.090 | $6.961e-4$ |
| | GLM BH | 0.260 | 4.192 | 16.127 | $7.596e-8$ |
| SSR | GLM ALL | 0.260 | 0.074 | 0.286 | 0.025 |
| | GLM BH | 0.260 | 0.056 | 0.216 | 0.024 |
| GLM ALL | GLM BH | 0.260 | 0.078 | 0.298 | 0.025 |
| | | **ME** $\Delta[HbR]$ | | | |
| Models | $P(M)$ | $P(M\|data)$ | BF$_M$ | BF$_{10}$ | error % |
| Null model (incl. subject) | 0.500 | 0.890 | 8.120 | 1.000 | |
| METHOD | 0.500 | 0.110 | 0.123 | 0.123 | 0.481 |

**Table 5** Descriptive Statistics of sRMSE resulting from the MI data

| | | | **MI** $\Delta[HbO]$ | | | |
|---|---|---|---|---|---|---|
| | NO SAC | CAR | GCR | SSR | GLM ALL | GLM BH |
| Mean | 0.236 | 0.244 | 0.243 | 0.242 | 0.240 | 0.242 |
| Std. Error of Mean | 0.003 | 0.003 | 0.003 | 0.003 | 0.003 | 0.003 |
| | | | **MI** $\Delta[HbR]$ | | | |
| | NO SAC | CAR | GCR | SSR | GLM ALL | GLM BH |
| Mean | 0.245 | 0.248 | 0.249 | 0.246 | 0.247 | 0.246 |
| Std. Error of Mean | 0.002 | 0.002 | 0.002 | 0.003 | 0.002 | 0.002 |

**Table 6** Results of the rmBANOVAs and corresponding post hoc tests with respect to the sRMSE analysis of MI data of both $\Delta[HbO]$ and $\Delta[HbR]$. $P(M|data)$ represents the posterior odds, $BF_{10}$ the Bayes factor and $error$ the error of the given test. $BF_{10} = 1$ indicates no evidence for neither $H_0$ nor $H_1$, $BF_{10} < 1$ indicates evidence in favor of the $H_0$, that is, that there are no differences between two methods and $BF_{10} > 1$ indicates evidence in favor of the $H_1$, that is, that there are differences between two methods. According to the guidelines of Lee and Wagenmakers (2014), Bayes factors $BF_{10}$ can be categorized in the following way: $BF_{10} < 1/100$ extreme evidence for $H_0$, $1/100 < BF_{10} < 1/30$ very strong evidence for $H_0$, $1/30 < BF_{10} < 1/10$ strong evidence for $H_0$, $1/10 < BF_{10} < 1/3$ moderate evidence for $H_0$, $1/3 < BF_{10} < 1$ anecdotal evidence for $H_0$, $1 < BBF_{10} < 3$ anecdotal evidence for $H_1$, $3 < BF_{10} < 10$ moderate evidence for $H_1$, $10 < BF_{10} < 30$ strong evidence for $H_1$, $30 < BF_{10} < 100$ very strong evidence for $H_1$, $BF_{10} > 100$ extreme evidence for $H_1$.

| **MI** $\Delta[HbO]$ | | | | | |
|---|---|---|---|---|---|
| Models | $P(M)$ | $P(M\|data)$ | BF$_M$ | BF$_{10}$ | error % |
| Null model (incl. subject) | 0.500 | 0.610 | 1.564 | 1.000 | |
| METHOD | 0.500 | 0.390 | 0.639 | 0.639 | 0.294 |
| **MI** $\Delta[HbR]$ | | | | | |
| Models | $P(M)$ | $P(M\|data)$ | BF$_M$ | BF$_{10}$ | error % |
| Null model (incl. subject) | 0.500 | 0.909 | 9.959 | 1.000 | |
| METHOD | 0.500 | 0.091 | 0.100 | 0.100 | 0.654 |

**Table 7** Descriptive Statistics of COR resulting from the SIM data.

| **SIM** $\Delta[HbO]$ | | | | | | |
|---|---|---|---|---|---|---|
| | NO SAC | CAR | GCR | SSR | GLM ALL | GLM BH |
| Mean | 0.852 | 1.307 | 1.291 | 1.259 | 1.578 | 1.254 |
| Std. Error of Mean | 0.103 | 0.099 | 0.096 | 0.093 | 0.077 | 0.103 |
| **SIM** $\Delta[HbR]$ | | | | | | |
| | NO SAC | CAR | GCR | SSR | GLM ALL | GLM BH |
| Mean | 1.243 | 1.400 | 1.337 | 1.305 | 1.460 | 1.380 |
| Std. Error of Mean | 0.083 | 0.077 | 0.070 | 0.089 | 0.070 | 0.080 |

**Table 8** Results of the rmBANOVAs and corresponding post hoc tests with respect to the COR analysis of SIM data of both $\Delta[HbO]$ and $\Delta[HbR]$. For all post hoc tests, the prior odds $P(M)$ were set to the default of 0.26, $P(M|data)$ represents the posterior odds, $BF_{10}$ the Bayes factor and $error$ the error of the given test. $BF_{10} = 1$ indicates no evidence for neither $H_0$ nor $H_1$, $BF_{10} < 1$ indicates evidence in favor of the $H_0$, that is, that there are no differences between two methods and $BF_{10} > 1$ indicates evidence in favor of the $H_1$, that is, that there are differences between two methods. According to the guidelines of Lee and Wagenmakers (2014), Bayes factors $BF_{10}$ can be categorized in the following way: $BF_{10} < 1/100$ extreme evidence for $H_0$, $1/100 < BF_{10} < 1/30$ very strong evidence for $H_0$, $1/30 < BF_{10} < 1/10$ strong evidence for $H_0$, $1/10 < BF_{10} < 1/3$ moderate evidence for $H_0$, $1/3 < BF_{10} < 1$ anecdotal evidence for $H_0$, $1 < BBF_{10} < 3$ anecdotal evidence for $H_1$, $3 < BF_{10} < 10$ moderate evidence for $H_1$, $10 < BF_{10} < 30$ strong evidence for $H_1$, $30 < BF_{10} < 100$ very strong evidence for $H_1$, $BF_{10} > 100$ extreme evidence for $H_1$.

| | | **SIM** $\Delta[HbO]$ | | | |
|---|---|---|---|---|---|
| Models | $P(M)$ | $P(M\|data)$ | BF$_M$ | BF$_{10}$ | error % |
| Null model (incl. subject) | 0.500 | $6.805e-15$ | $6.805e-15$ | 1.000 | |
| METHOD | 0.500 | 1.000 | $1.470e+14$ | $1.470e+14$ | 0.502 |
| | | Prior Odds | Posterior Odds | BF$_{10,U}$ | error % |
| NO SAC | CAR | 0.260 | 571.506 | 2198.769 | $1.097e-8$ |
| | GCR | 0.260 | 1132.643 | 4357.641 | $4.003e-9$ |
| | SSR | 0.260 | 107.664 | 414.217 | $2.142e-8$ |
| | GLM ALL | 0.260 | $3.257e+6$ | $1.253e+7$ | $4.115e-10$ |
| | GLM BH | 0.260 | 1859.217 | 7153.007 | $1.046e-9$ |
| CAR | GCR | 0.260 | 0.063 | 0.243 | 0.023 |
| | SSR | 0.260 | 0.068 | 0.263 | 0.023 |
| | GLM ALL | 0.260 | 16.380 | 63.020 | $1.796e-5$ |
| | GLM BH | 0.260 | 0.083 | 0.319 | 0.024 |
| GCR | SSR | 0.260 | 0.062 | 0.239 | 0.023 |
| | GLM ALL | 0.260 | 42.216 | 162.417 | $6.414e-8$ |
| | GLM BH | 0.260 | 0.070 | 0.271 | 0.024 |
| SSR | GLM ALL | 0.260 | 1108.489 | 4264.716 | $4.172e-9$ |
| | GLM BH | 0.260 | 0.057 | 0.220 | 0.022 |
| GLM ALL | GLM BH | 0.260 | 1255.218 | 4829.227 | $3.222e-9$ |

| | | **SIM** $\Delta[HbR]$ | | | |
|---|---|---|---|---|---|
| Models | $P(M)$ | $P(M\|data)$ | BF$_M$ | BF$_{10}$ | error % |
| Null model (incl. subject) | 0.500 | 0.170 | 0.205 | 1.000 | |
| METHOD | 0.500 | 0.830 | 4.875 | 4.875 | 0.525 |
| | | Prior Odds | Posterior Odds | BF$_{10,U}$ | error % |
| NO SAC | CAR | 0.260 | 0.480 | 1.847 | $1.722e-6$ |
| | GCR | 0.260 | 0.120 | 0.460 | 0.025 |
| | SSR | 0.260 | 0.122 | 0.470 | 0.025 |
| | GLM ALL | 0.260 | 30.999 | 119.264 | $8.891e-8$ |
| | GLM BH | 0.260 | 3.304 | 12.711 | $9.146e-8$ |
| CAR | GCR | 0.260 | 0.519 | 1.995 | $1.530e-6$ |
| | SSR | 0.260 | 0.122 | 0.471 | 0.025 |
| | GLM ALL | 0.260 | 0.086 | 0.329 | 0.024 |
| | GLM BH | 0.260 | 0.060 | 0.232 | 0.023 |
| GCR | SSR | 0.260 | 0.062 | 0.238 | 0.023 |
| | GLM ALL | 0.260 | 0.277 | 1.066 | 0.023 |
| | GLM BH | 0.260 | 0.072 | 0.276 | 0.024 |
| SSR | GLM ALL | 0.260 | 0.617 | 2.373 | $1.149e-6$ |
| | GLM BH | 0.260 | 0.123 | 0.473 | 0.025 |
| GLM ALL | GLM BH | 0.260 | 0.241 | 0.927 | 0.024 |

**Table 9** Descriptive Statistics of COR resulting from the ME data.

| | **ME** $\Delta[HbO]$ | | | | | |
|---|---|---|---|---|---|---|
| | NO SAC | CAR | GCR | SSR | GLM ALL | GLM BH |
| Mean | 0.722 | 0.626 | 0.582 | 0.771 | 0.848 | 0.754 |
| Std. Error of Mean | 0.081 | 0.077 | 0.077 | 0.079 | 0.074 | 0.080 |
| | **ME** $\Delta[HbR]$ | | | | | |
| | NO SAC | CAR | GCR | SSR | GLM ALL | GLM BH |
| Mean | 0.864 | 0.774 | 0.740 | 0.823 | 0.837 | 0.864 |
| Std. Error of Mean | 0.084 | 0.070 | 0.075 | 0.077 | 0.071 | 0.080 |

**Table 10** Results of the rmBANOVAs and corresponding post hoc tests with respect to the COR analysis of ME data of both $\Delta[HbO]$ and $\Delta[HbR]$. For all post hoc tests, the prior odds $P(M)$ were set to the default of 0.26, $P(M|data)$ represents the posterior odds, $BF_{10}$ the Bayes factor and $error$ the error of the given test. $BF_{10} = 1$ indicates no evidence for neither $H_0$ nor $H_1$, $BF_{10} < 1$ indicates evidence in favor of the $H_0$, that is, that there are no differences between two methods and $BF_{10} > 1$ indicates evidence in favor of the $H_1$, that is, that there are differences between two methods. According to the guidelines of Lee and Wagenmakers (2014), Bayes factors $BF_{10}$ can be categorized in the following way: $BF_{10} < 1/100$ extreme evidence for $H_0$, $1/100 < BF_{10} < 1/30$ very strong evidence for $H_0$, $1/30 < BF_{10} < 1/10$ strong evidence for $H_0$, $1/10 < BF_{10} < 1/3$ moderate evidence for $H_0$, $1/3 < BF_{10} < 1$ anecdotal evidence for $H_0$, $1 < BBF_{10} < 3$ anecdotal evidence for $H_1$, $3 < BF_{10} < 10$ moderate evidence for $H_1$, $10 < BF_{10} < 30$ strong evidence for $H_1$, $30 < BF_{10} < 100$ very strong evidence for $H_1$, $BF_{10} > 100$ extreme evidence for $H_1$.

| | | **ME** $\Delta[HbO]$ | | | |
|---|---|---|---|---|---|
| Models | $P(M)$ | $P(M\|data)$ | $BF_M$ | $BF_{10}$ | error % |
| Null model (incl. subject) | 0.500 | 0.003 | 0.003 | 1.000 | |
| METHOD | 0.500 | 0.997 | 298.511 | 298.511 | 0.323 |
| | | Prior Odds | Posterior Odds | $BF_{10,U}$ | error % |
| NO SAC | CAR | 0.260 | 0.137 | 0.528 | 0.026 |
| | GCR | 0.260 | 0.212 | 0.814 | 0.025 |
| | SSR | 0.260 | 0.069 | 0.266 | 0.025 |
| | GLM ALL | 0.260 | 0.405 | 1.559 | $2.133e-6$ |
| | GLM BH | 0.260 | 0.065 | 0.248 | 0.025 |
| CAR | GCR | 0.260 | 0.141 | 0.541 | 0.026 |
| | SSR | 0.260 | 0.783 | 3.012 | $7.548e-7$ |
| | GLM ALL | 0.260 | 120.828 | 464.864 | $1.788e-8$ |
| | GLM BH | 0.260 | 0.847 | 3.260 | $6.498e-7$ |
| GCR | SSR | 0.260 | 1.680 | 6.465 | $1.364e-7$ |
| | GLM ALL | 0.260 | 32.725 | 125.903 | $2.494e-4$ |
| | GLM BH | 0.260 | 1.599 | 6.153 | $1.541e-7$ |
| SSR | GLM ALL | 0.260 | 0.138 | 0.531 | 0.026 |
| | GLM BH | 0.260 | 0.059 | 0.227 | 0.024 |
| GLM ALL | GLM BH | 0.260 | 0.699 | 2.688 | $9.273e-7$ |
| | | **ME** $\Delta[HbR]$ | | | |
| Models | $P(M)$ | $P(M\|data)$ | $BF_M$ | $BF_{10}$ | error % |
| Null model (incl. subject) | 0.500 | 0.405 | 0.682 | 1.000 | |
| METHOD | 0.500 | 0.595 | 1.467 | 1.467 | 0.526 |

**Table 11** Descriptive Statistics of COR resulting from the MI data.

| | **MI** $\Delta[HbO]$ | | | | | |
|---|---|---|---|---|---|---|
| | NO SAC | CAR | GCR | SSR | GLM ALL | GLM BH |
| Mean | 0.332 | 0.216 | 0.200 | 0.271 | 0.289 | 0.253 |
| Std. Error of Mean | 0.049 | 0.045 | 0.045 | 0.035 | 0.046 | 0.042 |
| | **MI** $\Delta[HbR]$ | | | | | |
| | NO SAC | CAR | GCR | SSR | GLM ALL | GLM BH |
| Mean | 0.238 | 0.162 | 0.134 | 0.191 | 0.222 | 0.229 |
| Std. Error of Mean | 0.045 | 0.043 | 0.039 | 0.043 | 0.046 | 0.043 |

**Table 12** Results of the rmBANOVAs and corresponding post hoc tests with respect to the COR analysis of MI data of both $\Delta[HbO]$ and $\Delta[HbR]$. $P(M|data)$ represents the posterior odds, $BF_{10}$ the Bayes factor and $error$ the error of the given test. $BF_{10} = 1$ indicates no evidence for neither $H_0$ nor $H_1$, $BF_{10} < 1$ indicates evidence in favor of the $H_0$, that is, that there are no differences between two methods and $BF_{10} > 1$ indicates evidence in favor of the $H_1$, that is, that there are differences between two methods. According to the guidelines of Lee and Wagenmakers (2014), Bayes factors $BF_{10}$ can be categorized in the following way: $BF_{10} < 1/100$ extreme evidence for $H_0$, $1/100 < BF_{10} < 1/30$ very strong evidence for $H_0$, $1/30 < BF_{10} < 1/10$ strong evidence for $H_0$, $1/10 < BF_{10} < 1/3$ moderate evidence for $H_0$, $1/3 < BF_{10} < 1$ anecdotal evidence for $H_0$, $1 < BBF_{10} < 3$ anecdotal evidence for $H_1$, $3 < BF_{10} < 10$ moderate evidence for $H_1$, $10 < BF_{10} < 30$ strong evidence for $H_1$, $30 < BF_{10} < 100$ very strong evidence for $H_1$, $BF_{10} > 100$ extreme evidence for $H_1$.

| | | **MI** $\Delta[HbO]$ | | | |
|---|---|---|---|---|---|
| Models | $P(M)$ | $P(M\|data)$ | $BF_M$ | $BF_{10}$ | error % |
| Null model (incl. subject) | 0.500 | 0.256 | 0.345 | 1.000 | |
| METHOD | 0.500 | 0.744 | 2.900 | 2.900 | 0.325 |
| | | **MI** $\Delta[HbR]$ | | | |
| Models | $P(M)$ | $P(M\|data)$ | $BF_M$ | $BF_{10}$ | error % |
| Null model (incl. subject) | 0.500 | 0.441 | 0.788 | 1.000 | |
| METHOD | 0.500 | 0.559 | 1.269 | 1.269 | 0.276 |

**Table 13** Descriptive Statistics of CNR resulting from the SIM data.

| | **SIM** $\Delta[HbO]$ | | | | | |
|---|---|---|---|---|---|---|
| | NO SAC | CAR | GCR | SSR | GLM ALL | GLM BH |
| Mean | 10.178 | 16.073 | 14.768 | 16.747 | 21.791 | 15.717 |
| Std. Error of Mean | 1.359 | 1.910 | 1.727 | 2.016 | 1.169 | 1.737 |
| | **SIM** $\Delta[HbR]$ | | | | | |
| | NO SAC | CAR | GCR | SSR | GLM ALL | GLM BH |
| Mean | 15.254 | 18.220 | 18.330 | 16.580 | 19.073 | 17.647 |
| Std. Error of Mean | 1.613 | 1.685 | 1.831 | 1.449 | 1.690 | 1.694 |

**Table 14** Results of the rmBANOVAs and corresponding post hoc tests with respect to the CNR analysis of SIM data of both $\Delta[HbO]$ and $\Delta[HbR]$. For all post hoc tests, the prior odds $P(M)$ were set to the default of 0.26, $P(M|data)$ represents the posterior odds, $BF_{10}$ the Bayes factor and $error$ the error of the given test. $BF_{10} = 1$ indicates no evidence for neither $H_0$ nor $H_1$, $BF_{10} < 1$ indicates evidence in favor of the $H_0$, that is, that there are no differences between two methods and $BF_{10} > 1$ indicates evidence in favor of the $H_1$, that is, that there are differences between two methods. According to the guidelines of Lee and Wagenmakers (2014), Bayes factors $BF_{10}$ can be categorized in the following way: $BF_{10} < 1/100$ extreme evidence for $H_0$, $1/100 < BF_{10} < 1/30$ very strong evidence for $H_0$, $1/30 < BF_{10} < 1/10$ strong evidence for $H_0$, $1/10 < BF_{10} < 1/3$ moderate evidence for $H_0$, $1/3 < BF_{10} < 1$ anecdotal evidence for $H_0$, $1 < BBF_{10} < 3$ anecdotal evidence for $H_1$, $3 < BF_{10} < 10$ moderate evidence for $H_1$, $10 < BF_{10} < 30$ strong evidence for $H_1$, $30 < BF_{10} < 100$ very strong evidence for $H_1$, $BF_{10} > 100$ extreme evidence for $H_1$.

| | | | | | |
|---|---|---|---|---|---|
| | | **SIM** $\Delta[HbO]$ | | | |
| Null model (incl. subject) | 0.500 | $4.632e-6$ | $4.632e-6$ | 1.000 | |
| METHOD | 0.500 | 1.000 | 215906.220 | 215906.220 | 0.297 |
| | | Prior Odds | Posterior Odds | $\text{BF}_{10,U}$ | error % |
| NO SAC | CAR | 0.260 | 3.042 | 11.704 | $9.352e-8$ |
| | GCR | 0.260 | 1.484 | 5.710 | $1.719e-7$ |
| | SSR | 0.260 | 3.178 | 12.227 | $9.221e-8$ |
| | GLM ALL | 0.260 | 248839.853 | 957367.066 | $2.843e-10$ |
| | GLM BH | 0.260 | 11.586 | 44.575 | $4.422e-4$ |
| CAR | GCR | 0.260 | 0.118 | 0.452 | 0.025 |
| | SSR | 0.260 | 0.060 | 0.232 | 0.023 |
| | GLM ALL | 0.260 | 1.689 | 6.498 | $1.376e-7$ |
| | GLM BH | 0.260 | 0.058 | 0.223 | 0.022 |
| GCR | SSR | 0.260 | 0.096 | 0.368 | 0.025 |
| | GLM ALL | 0.260 | 10.513 | 40.449 | $6.404e-4$ |
| | GLM BH | 0.260 | 0.065 | 0.251 | 0.023 |
| SSR | GLM ALL | 0.260 | 1.418 | 5.457 | $1.916e-7$ |
| | GLM BH | 0.260 | 0.078 | 0.300 | 0.024 |
| GLM ALL | GLM BH | 0.260 | 9.730 | 37.434 | $8.231e-4$ |
| | | **SIM** $\Delta[HbR]$ | | | |
| Models | $P(M)$ | $P(M|data)$ | $\text{BF}_M$ | $\text{BF}_{10}$ | error % |
| Null model (incl. subject) | 0.500 | 0.595 | 1.467 | 1.000 | |
| METHOD | 0.500 | 0.405 | 0.682 | 0.682 | 0.393 |

**Table 15** Descriptive Statistics of CNR resulting from the ME data.

| | | | | | | |
|---|---|---|---|---|---|---|
| | | | **ME** $\Delta[HbO]$ | | | |
| | NO SAC | CAR | GCR | SSR | GLM ALL | GLM BH |
| Mean | 9.248 | 9.294 | 8.467 | 11.260 | 13.025 | 11.508 |
| Std. Error of Mean | 1.031 | 0.844 | 0.810 | 1.154 | 1.211 | 1.105 |
| | | | **ME** $\Delta[HbR]$ | | | |
| | NO SAC | CAR | GCR | SSR | GLM ALL | GLM BH |
| Mean | 13.343 | 12.192 | 11.731 | 13.166 | 14.383 | 13.822 |
| Std. Error of Mean | 1.060 | 0.819 | 0.762 | 0.994 | 0.934 | 1.066 |

**Table 16** Results of the rmBANOVAs and corresponding post hoc tests with respect to the CNR analysis of ME data of both $\Delta[HbO]$ and $\Delta[HbR]$. For all post hoc tests, the prior odds $P(M)$ were set to the default of 0.26, $P(M|data)$ represents the posterior odds, $BF_{10}$ the Bayes factor and $error$ the error of the given test. $BF_{10} = 1$ indicates no evidence for neither $H_0$ nor $H_1$, $BF_{10} < 1$ indicates evidence in favor of the $H_0$, that is, that there are no differences between two methods and $BF_{10} > 1$ indicates evidence in favor of the $H_1$, that is, that there are differences between two methods. According to the guidelines of Lee and Wagenmakers (2014), Bayes factors $BF_{10}$ can be categorized in the following way: $BF_{10} < 1/100$ extreme evidence for $H_0$, $1/100 < BF_{10} < 1/30$ very strong evidence for $H_0$, $1/30 < BF_{10} < 1/10$ strong evidence for $H_0$, $1/10 < BF_{10} < 1/3$ moderate evidence for $H_0$, $1/3 < BF_{10} < 1$ anecdotal evidence for $H_0$, $1 < BBF_{10} < 3$ anecdotal evidence for $H_1$, $3 < BF_{10} < 10$ moderate evidence for $H_1$, $10 < BF_{10} < 30$ strong evidence for $H_1$, $30 < BF_{10} < 100$ very strong evidence for $H_1$, $BF_{10} > 100$ extreme evidence for $H_1$.

| | | | | | |
|---|---|---|---|---|---|
| | | **ME** $\Delta[HbO]$ | | | |
| Null model (incl. subject) | 0.500 | $2.131e-7$ | $2.131e-7$ | 1.000 | |
| METHOD | 0.500 | 1.000 | $4.692e+6$ | $4.692e+6$ | 0.330 |
| | | Prior Odds | Posterior Odds | BF$_{10,U}$ | error % |
| NO SAC | CAR | 0.260 | 0.056 | 0.215 | 0.024 |
| | GCR | 0.260 | 0.075 | 0.289 | 0.025 |
| | SSR | 0.260 | 0.782 | 3.008 | $7.569e-7$ |
| | GLM ALL | 0.260 | 387.678 | 1491.523 | $1.388e-8$ |
| | GLM BH | 0.260 | 5.136 | 19.759 | $9.136e-8$ |
| CAR | GCR | 0.260 | 2.080 | 8.001 | $9.751e-8$ |
| | SSR | 0.260 | 1.102 | 4.240 | $3.771e-7$ |
| | GLM ALL | 0.260 | 128.127 | 492.947 | $1.694e-8$ |
| | GLM BH | 0.260 | 7.763 | 29.867 | $1.155e-7$ |
| GCR | SSR | 0.260 | 4.873 | 18.750 | $8.731e-8$ |
| | GLM ALL | 0.260 | 267.328 | 1028.497 | $1.436e-8$ |
| | GLM BH | 0.260 | 29.698 | 114.259 | $1.866e-4$ |
| SSR | GLM ALL | 0.260 | 1.700 | 6.541 | $1.326e-7$ |
| | GLM BH | 0.260 | 0.061 | 0.234 | 0.024 |
| GLM ALL | GLM BH | 0.260 | 5.758 | 22.153 | $9.938e-8$ |
| | | **ME** $\Delta[HbR]$ | | | |
| Models | $P(M)$ | $P(M|data)$ | BF$_M$ | BF$_{10}$ | error % |
| Null model (incl. subject) | 0.500 | 0.010 | 0.010 | 1.000 | |
| METHOD | 0.500 | 0.990 | 103.042 | 103.042 | 0.298 |
| | | Prior Odds | Posterior Odds | BF$_{10,U}$ | error % |
| NO SAC | CAR | 0.260 | 0.180 | 0.693 | 0.025 |
| | GCR | 0.260 | 0.444 | 1.709 | $1.886e-6$ |
| | SSR | 0.260 | 0.059 | 0.229 | 0.024 |
| | GLM ALL | 0.260 | 0.459 | 1.764 | $1.803e-6$ |
| | GLM BH | 0.260 | 0.104 | 0.398 | 0.026 |
| CAR | GCR | 0.260 | 0.088 | 0.340 | 0.026 |
| | SSR | 0.260 | 0.138 | 0.529 | 0.026 |
| | GLM ALL | 0.260 | 8.889 | 34.200 | $1.198e-7$ |
| | GLM BH | 0.260 | 0.320 | 1.232 | 0.023 |
| GCR | SSR | 0.260 | 0.404 | 1.556 | $2.138e-6$ |
| | GLM ALL | 0.260 | 17.751 | 68.295 | $6.716e-5$ |
| | GLM BH | 0.260 | 1.278 | 4.916 | $2.684e-7$ |
| SSR | GLM ALL | 0.260 | 0.628 | 2.415 | $1.114e-6$ |
| | GLM BH | 0.260 | 0.095 | 0.366 | 0.026 |
| GLM ALL | GLM BH | 0.260 | 0.081 | 0.313 | 0.025 |

**Table 17** Descriptive Statistics of CNR resulting from the MI data.

| | NO SAC | CAR | GCR | SSR | GLM ALL | GLM BH |
|---|---|---|---|---|---|---|
| **MI** $\Delta[HbO]$ | | | | | | |
| Mean | 6.012 | 5.434 | 5.307 | 5.962 | 6.511 | 6.061 |
| Std. Error of Mean | 0.507 | 0.529 | 0.512 | 0.377 | 0.573 | 0.544 |
| **MI** $\Delta[HbR]$ | | | | | | |
| | NO SAC | CAR | GCR | SSR | GLM ALL | GLM BH |
| Mean | 6.490 | 5.982 | 5.819 | 6.336 | 6.795 | 6.569 |
| Std. Error of Mean | 0.574 | 0.478 | 0.440 | 0.585 | 0.524 | 0.453 |

**Table 18** Results of the rmBANOVAs and corresponding post hoc tests with respect to the CNR analysis of MI data of both $\Delta[HbO]$ and $\Delta[HbR]$. $P(M|data)$ represents the posterior odds, $BF_{10}$ the Bayes factor and $error$ the error of the given test. $BF_{10} = 1$ indicates no evidence for neither $H_0$ nor $H_1$, $BF_{10} < 1$ indicates evidence in favor of the $H_0$, that is, that there are no differences between two methods and $BF_{10} > 1$ indicates evidence in favor of the $H_1$, that is, that there are differences between two methods. According to the guidelines of Lee and Wagenmakers (2014), Bayes factors $BF_{10}$ can be categorized in the following way: $BF_{10} < 1/100$ extreme evidence for $H_0$, $1/100 < BF_{10} < 1/30$ very strong evidence for $H_0$, $1/30 < BF_{10} < 1/10$ strong evidence for $H_0$, $1/10 < BF_{10} < 1/3$ moderate evidence for $H_0$, $1/3 < BF_{10} < 1$ anecdotal evidence for $H_0$, $1 < BBF_{10} < 3$ anecdotal evidence for $H_1$, $3 < BF_{10} < 10$ moderate evidence for $H_1$, $10 < BF_{10} < 30$ strong evidence for $H_1$, $30 < BF_{10} < 100$ very strong evidence for $H_1$, $BF_{10} > 100$ extreme evidence for $H_1$.

| | | | | | |
|---|---|---|---|---|---|
| **MI** $\Delta[HbO]$ | | | | | |
| Null model (incl. subject) | 0.500 | 0.611 | 1.572 | 1.000 | |
| METHOD | 0.500 | 0.389 | 0.636 | 0.636 | 0.352 |
| **MI** $\Delta[HbR]$ | | | | | |
| Models | $P(M)$ | $P(M\|data)$ | BF$_M$ | BF$_{10}$ | error % |
| Null model (incl. subject) | 0.500 | 0.711 | 2.459 | 1.000 | |
| METHOD | 0.500 | 0.289 | 0.407 | 0.407 | 0.360 |

**Table 19** Results of the Bayesian t-tests of the SDC CORMAT analysis of the $\Delta[HbR]$ data of SIM, ME LEFT, ME RIGHT and MI data. $BF_{10}$ represent Bayes factors $\pm$ an error term in %. Bayesian t-tests were performed with the *BayesFactor* package[?] in R[?] for which the default prior odds is set at $P(M) = \frac{\sqrt{2}}{2}$. Corresponding SDC CORMATs are visualized in Figures 6 A, 8 A as well as 10 A.

| | | | | | |
|---|---|---|---|---|---|
| | | | **SIM** | | |
| | | | $\Delta[HbO]$ | | |
| | NO SAC | CAR | GCR | SSR | GLM ALL |
| NO SAC | - | - | - | - | - |
| CAR | $BF_{10} = 3.60 \cdot 10^{117} \pm 0\%$ | - | - | - | - |
| GCR | $BF_{10} = 2.47 \cdot 10^{113} \pm 0\%$ | $BF_{10} = 0.07 \pm 0.06\%$ | - | - | - |
| SSR | $BF_{10} = 3.06 \cdot 10^{116} \pm 0\%$ | $BF_{10} = 2.06 \cdot 10^{60} \pm 0\%$ | $BF_{10} = 1.02 \cdot 10^{52} \pm 0\%$ | - | - |
| GLM ALL | $BF_{10} = 2.29 \cdot 10^{115} \pm 0\%$ | $BF_{10} = 7.73 \cdot 10^{35} \pm 0\%$ | $BF_{10} = 4.81 \cdot 10^{28} \pm 0\%$ | $BF_{10} = 5.92 \cdot 10^{25} \pm 0\%$ | - |
| GLM BH | $BF_{10} = 4.17 \cdot 10^{110} \pm 0\%$ | $BF_{10} = 9.07 \cdot 10^{70} \pm 0\%$ | $BF_{10} = 6.72 \cdot 10^{56} \pm 0\%$ | $BF_{10} = 100608 \pm 0\%$ | $BF_{10} = 4.73 \cdot 10^{60} \pm 0\%$ |
| | | | **ME LEFT** | | |
| | | | $\Delta[HbO]$ | | |
| | NO SAC | CAR | GCR | SSR | GLM ALL |
| NO SAC | - | - | - | - | - |
| CAR | $BF_{10} = 1.80 \cdot 10^{113} \pm 0\%$ | - | - | - | - |
| GCR | $BF_{10} = 1.96 \cdot 10^{112} \pm 0\%$ | $BF_{10} = 0.08 \pm 0.05\%$ | - | - | - |
| SSR | $BF_{10} = 5.70 \cdot 10^{114} \pm 0\%$ | $BF_{10} = 4.07 \cdot 10^{64} \pm 0\%$ | $BF_{10} = 5.03 \cdot 10^{60} \pm 0\%$ | - | - |
| GLM ALL | $BF_{10} = 1.52 \cdot 10^{120} \pm 0\%$ | $BF_{10} = 3.91 \cdot 10^{50} \pm 0\%$ | $BF_{10} = 4.25 \cdot 10^{44} \pm 0\%$ | $BF_{10} = 4.69 \cdot 10^{24} \pm 0\%$ | - |
| GLM BH | $BF_{10} = 8.98 \cdot 10^{115} \pm 0\%$ | $BF_{10} = 1.82 \cdot 10^{72} \pm 0\%$ | $BF_{10} = 1.41 \cdot 10^{63} \pm 0\%$ | $BF_{10} = 0.41 \pm 0.01\%$ | $BF_{10} = 2.71 \cdot 10^{38} \pm 0\%$ |
| | | | **ME RIGHT** | | |
| | | | $\Delta[HbO]$ | | |
| | NO SAC | CAR | GCR | SSR | GLM ALL |
| NO SAC | - | - | - | - | - |
| CAR | $BF_{10} = 3.96 \cdot 10^{113} \pm 0\%$ | - | - | - | - |
| GCR | $BF_{10} = 7.64 \cdot 10^{112} \pm 0\%$ | $BF_{10} = 0.07 \pm 0.06\%$ | - | - | - |
| SSR | $BF_{10} = 2.33 \cdot 10^{114} \pm 0\%$ | $BF_{10} = 2.45 \cdot 10^{60} \pm 0\%$ | $BF_{10} = 1.36 \cdot 10^{59} \pm 0\%$ | - | - |
| GLM ALL | $BF_{10} = 1.00 \cdot 10^{121} \pm 0\%$ | $BF_{10} = 3.18 \cdot 10^{46} \pm 0\%$ | $BF_{10} = 1.02 \cdot 10^{41} \pm 0\%$ | $BF_{10} = 2.15 \cdot 10^{32} \pm 0\%$ | - |
| GLM BH | $BF_{10} = 1.85 \cdot 10^{114} \pm 0\%$ | $BF_{10} = 1.09 \cdot 10^{70} \pm 0\%$ | $BF_{10} = 2.21 \cdot 10^{62} \pm 0\%$ | $BF_{10} = 42.26 \pm 0\%$ | $BF_{10} = 3.45 \cdot 10^{54} \pm 0\%$ |
| | | | **MI** | | |
| | | | $\Delta[HbO]$ | | |
| | NO SAC | CAR | GCR | SSR | GLM ALL |
| NO SAC | - | - | - | - | - |
| CAR | $BF_{10} = 7.90 \cdot 10^{113} \pm 0\%$ | - | - | - | - |
| GCR | $BF_{10} = 9.77 \cdot 10^{114} \pm 0\%$ | $BF_{10} = 0.09 \pm 0.05\%$ | - | - | - |
| SSR | $BF_{10} = 8.52 \cdot 10^{118} \pm 0\%$ | $BF_{10} = 8.04 \cdot 10^{62} \pm 0\%$ | $BF_{10} = 3.60 \cdot 10^{61} \pm 0\%$ | - | - |
| GLM ALL | $BF_{10} = 1.62 \cdot 10^{123} \pm 0\%$ | $BF_{10} = 2.74 \cdot 10^{50} \pm 0\%$ | $BF_{10} = 2.22 \cdot 10^{45} \pm 0\%$ | $BF_{10} = 1.70 \cdot 10^{34} \pm 0\%$ | - |
| GLM BH | $BF_{10} = 8.77 \cdot 10^{115} \pm 0\%$ | $BF_{10} = 2.42 \cdot 10^{73} \pm 0\%$ | $BF_{10} = 2.60 \cdot 10^{67} \pm 0\%$ | $BF_{10} = 5.78 \cdot 10^{5} \pm 0\%$ | $BF_{10} = 1.18 \cdot 10^{64} \pm 0\%$ |

**Table 20** Results of the Bayesian t-tests of the SDC CORMAT analysis of the $\Delta[HbR]$ data of SIM, ME LEFT, ME RIGHT and MI data. $BF_{10}$ represent Bayes factors $\pm$ an error term in %. Bayesian t-tests were performed with the *BayesFactor* package[?] in R[?] for which the default prior odds is set at $P(M) = \frac{\sqrt{2}}{2}$. Corresponding SDC CORMATs are visualized in Figures 6 A, 8 A as well as 10 A.

**SIM**

$\Delta[HbR]$

| | NO SAC | CAR | GCR | SSR | GLM ALL |
|---|---|---|---|---|---|
| NO SAC | - | - | - | - | - |
| CAR | $BF_{10} = 2.63 \cdot 10^{91} \pm 0\%$ | - | - | - | - |
| GCR | $BF_{10} = 1.20 \cdot 10^{80} \pm 0\%$ | $BF_{10} = 0.08 \pm 0.05\%$ | - | - | - |
| SSR | $BF_{10} = 5.19 \cdot 10^{74} \pm 0\%$ | $BF_{10} = 3.25 \cdot 10^{57} \pm 0\%$ | $BF_{10} = 5.08 \cdot 10^{49} \pm 0\%$ | - | - |
| GLM ALL | $BF_{10} = 1.04 \cdot 10^{82} \pm 0\%$ | $BF_{10} = 2.03 \cdot 10^{37} \pm 0\%$ | $BF_{10} = 1.84 \cdot 10^{30} \pm 0\%$ | $BF_{10} = 2.81 \cdot 10^{27} \pm 0\%$ | - |
| GLM BH | $BF_{10} = 2.41 \cdot 10^{53} \pm 0\%$ | $BF_{10} = 5.28 \cdot 10^{20} \pm 0\%$ | $BF_{10} = 4.19 \cdot 10^{17} \pm 0\%$ | $BF_{10} = 1.26 \cdot 10^{13} \pm 0\%$ | $BF_{10} = 0.20 \pm 0.02\%$ |

**ME LEFT**

$\Delta[HbR]$

| | NO SAC | CAR | GCR | SSR | GLM ALL |
|---|---|---|---|---|---|
| NO SAC | - | - | - | - | - |
| CAR | $BF_{10} = 1.92 \cdot 10^{63} \pm 0\%$ | - | - | - | - |
| GCR | $BF_{10} = 4.29 \cdot 10^{64} \pm 0\%$ | $BF_{10} = 0.07 \pm 0.06\%$ | - | - | - |
| SSR | $BF_{10} = 2.77 \cdot 10^{24} \pm 0\%$ | $BF_{10} = 7.85 \cdot 10^{46} \pm 0\%$ | $BF_{10} = 5.81 \cdot 10^{43} \pm 0\%$ | - | - |
| GLM ALL | $BF_{10} = 9.94 \cdot 10^{36} \pm 0\%$ | $BF_{10} = 4.51 \cdot 10^{38} \pm 0\%$ | $BF_{10} = 3.62 \cdot 10^{35} \pm 0\%$ | $BF_{10} = 4.70 \cdot 10^{22} \pm 0\%$ | - |
| GLM BH | $BF_{10} = 5.11 \cdot 10^{15} \pm 0\%$ | $BF_{10} = 2.27 \cdot 10^{54} \pm 0\%$ | $BF_{10} = 2.74 \cdot 10^{50} \pm 0\%$ | $BF_{10} = 5.09 \cdot 10^{8} \pm 0\%$ | $BF_{10} = 2.21 \cdot 10^{63} \pm 0\%$ |

**ME RIGHT**

$\Delta[HbR]$

| | NO SAC | CAR | GCR | SSR | GLM ALL |
|---|---|---|---|---|---|
| NO SAC | - | - | - | - | - |
| CAR | $BF_{10} = 8.41 \cdot 10^{85} \pm 0\%$ | - | - | - | - |
| GCR | $BF_{10} = 1.75 \cdot 10^{79} \pm 0\%$ | $BF_{10} = 0.07 \pm 0.06\%$ | - | - | - |
| SSR | $BF_{10} = 2.64 \cdot 10^{74} \pm 0\%$ | $BF_{10} = 2.94 \cdot 10^{57} \pm 0\%$ | $BF_{10} = 1.06 \cdot 10^{53} \pm 0\%$ | - | - |
| GLM ALL | $BF_{10} = 4.36 \cdot 10^{92} \pm 0\%$ | $BF_{10} = 9.68 \cdot 10^{43} \pm 0\%$ | $BF_{10} = 3.42 \cdot 10^{39} \pm 0\%$ | $BF_{10} = 1.44 \cdot 10^{43} \pm 0\%$ | - |
| GLM BH | $BF_{10} = 9.67 \cdot 10^{75} \pm 0\%$ | $BF_{10} = 3.06 \cdot 10^{61} \pm 0\%$ | $BF_{10} = 4.61 \cdot 10^{55} \pm 0\%$ | $BF_{10} = 0.39 \pm 0.01\%$ | $BF_{10} = 1.02 \cdot 10^{70} \pm 0\%$ |

**MI**

$\Delta[HbR]$

| | NO SAC | CAR | GCR | SSR | GLM ALL |
|---|---|---|---|---|---|
| NO SAC | - | - | - | - | - |
| CAR | $BF_{10} = 3.21 \cdot 10^{74} \pm 0\%$ | - | - | - | - |
| GCR | $BF_{10} = 2.44 \cdot 10^{69} \pm 0\%$ | $BF_{10} = 0.07 \pm 0.06\%$ | - | - | - |
| SSR | $BF_{10} = 3.05 \cdot 10^{72} \pm 0\%$ | $BF_{10} = 6.34 \cdot 10^{41} \pm 0\%$ | $BF_{10} = 8.69 \cdot 10^{38} \pm 0\%$ | - | - |
| GLM ALL | $BF_{10} = 4.34 \cdot 10^{91} \pm 0\%$ | $BF_{10} = 2.28 \cdot 10^{33} \pm 0\%$ | $BF_{10} = 2.21 \cdot 10^{30} \pm 0\%$ | $BF_{10} = 1.39 \cdot 10^{33} \pm 0\%$ | - |
| GLM BH | $BF_{10} = 9.59 \cdot 10^{74} \pm 0\%$ | $BF_{10} = 3.45 \cdot 10^{51} \pm 0\%$ | $BF_{10} = 1.03 \cdot 10^{47} \pm 0\%$ | $BF_{10} = 2.85 \cdot 10^{5} \pm 0\%$ | $BF_{10} = 1.56 \cdot 10^{71} \pm 0\%$ |

**Table 21** Results of the Bayesian t-tests of the BETA MAPS analysis of the SIM $\Delta[HbO]$ data. Mean ± SEM represent mean beta values of the respective channel and its standard error of the mean across participants, $BF_{10}$ represent Bayes factors. Bayesian t-tests were performed with the *BayesFactor* package in R for which the default prior odds is set at $P(M) = \frac{\sqrt{2}}{2}$. Corresponding BETA MAPS are visualized in Figure 6 B of the main document.

| | | | | **SIM** | | |
|---|---|---|---|---|---|---|
| Channel (Channel Frequency after Pruning) | **NO SAC** | **CAR** | **GCR** | **SSR** | **GLM ALL** | **GLM BH** |
| 1 (20) | $\overline{\beta} = 0.28 \pm 0.81$<br>$BF_{10} = 0.25$ | $\overline{\beta} = -0.75 \pm 0.38$<br>$BF_{10} = 1.18$ | $\overline{\beta} = -0.10 \pm 0.22$<br>$BF_{10} = 0.25$ | $\overline{\beta} = 0.07 \pm 0.34$<br>$BF_{10} = 0.24$ | $\overline{\beta} = -0.05 \pm 0.15$<br>$BF_{10} = 0.25$ | $\overline{\beta} = 0.00 \pm 0.32$<br>$BF_{10} = 0.23$ |
| 2 (19) | $\overline{\beta} = -0.35 \pm 0.92$<br>$BF_{10} = 0.25$ | $\overline{\beta} = -1.37 \pm 0.38$<br>$\mathbf{BF_{10} = 21.77}$ | $\overline{\beta} = -1.25 \pm 0.25$<br>$\mathbf{BF_{10} = 124.96}$ | $\overline{\beta} = -0.43 \pm 0.41$<br>$BF_{10} = 0.38$ | $\overline{\beta} = -0.04 \pm 0.23$<br>$BF_{10} = 0.24$ | $\overline{\beta} = -0.31 \pm 0.47$<br>$BF_{10} = 0.29$ |
| 4 (19) | $\overline{\beta} = -0.47 \pm 0.82$<br>$BF_{10} = 0.28$ | $\overline{\beta} = -1.54 \pm 0.39$<br>$\mathbf{BF_{10} = 36.18}$ | $\overline{\beta} = -0.95 \pm 0.25$<br>$\mathbf{BF_{10} = 28.35}$ | $\overline{\beta} = -0.15 \pm 0.44$<br>$BF_{10} = 0.25$ | $\overline{\beta} = -0.09 \pm 0.16$<br>$BF_{10} = 0.27$ | $\overline{\beta} = -0.53 \pm 0.31$<br>$BF_{10} = 0.84$ |
| 6 (23) | $\overline{\beta} = 8.16 \pm 0.87$<br>$\mathbf{BF_{10} = 3.42 \cdot 10^{6}}$ | $\overline{\beta} = 7.22 \pm 0.43$<br>$\mathbf{BF_{10} = 1.38 \cdot 10^{11}}$ | $\overline{\beta} = 5.70 \pm 0.32$<br>$\mathbf{BF_{10} = 4.08 \cdot 10^{11}}$ | $\overline{\beta} = 7.96 \pm 0.52$<br>$\mathbf{BF_{10} = 1.93 \cdot 10^{10}}$ | $\overline{\beta} = 7.30 \pm 0.38$<br>$\mathbf{BF_{10} = 1.66 \cdot 10^{12}}$ | $\overline{\beta} = 8.61 \pm 0.49$<br>$\mathbf{BF_{10} = 2.54 \cdot 10^{11}}$ |
| 7 (18) | $\overline{\beta} = 0.00 \pm 0.94$<br>$BF_{10} = 0.24$ | $\overline{\beta} = -0.72 \pm 0.32$<br>$BF_{10} = 1.83$ | $\overline{\beta} = -1.94 \pm 0.29$<br>$\mathbf{BF_{10} = 5120.13}$ | $\overline{\beta} = 0.34 \pm 0.43$<br>$BF_{10} = 0.32$ | $\overline{\beta} = 0.11 \pm 0.13$<br>$BF_{10} = 0.33$ | $\overline{\beta} = 0.09 \pm 0.49$<br>$BF_{10} = 0.25$ |
| 8 (23) | $\overline{\beta} = 4.17 \pm 0.87$<br>$\mathbf{BF_{10} = 303.90}$ | $\overline{\beta} = 3.12 \pm 0.32$<br>$\mathbf{BF_{10} = 4.58 \cdot 10^{6}}$ | $\overline{\beta} = 1.78 \pm 0.26$<br>$\mathbf{BF_{10} = 2.88 \cdot 10^{4}}$ | $\overline{\beta} = 3.95 \pm 0.36$<br>$\mathbf{BF_{10} = 4.47 \cdot 10^{7}}$ | $\overline{\beta} = 3.00 \pm 0.30$<br>$\mathbf{BF_{10} = 1.06 \cdot 10^{7}}$ | $\overline{\beta} = 4.03 \pm 0.41$<br>$\mathbf{BF_{10} = 6.72 \cdot 10^{6}}$ |
| 10 (13) | $\overline{\beta} = -0.59 \pm 1.08$<br>$BF_{10} = 0.32$ | $\overline{\beta} = -1.34 \pm 0.57$<br>$BF_{10} = 2.12$ | $\overline{\beta} = -0.63 \pm 0.39$<br>$BF_{10} = 0.476$ | $\overline{\beta} = 0.44 \pm 0.56$<br>$BF_{10} = 0.36$ | $\overline{\beta} = 0.10 \pm 0.24$<br>$BF_{10} = 0.30$ | $\overline{\beta} = -0.08 \pm 0.40$<br>$BF_{10} = 0.28$ |
| 11 (20) | $\overline{\beta} = 0.05 \pm 0.74$<br>$BF_{10} = 0.23$ | $\overline{\beta} = -1.00 \pm 0.26$<br>$\mathbf{BF_{10} = 38.05}$ | $\overline{\beta} = -0.53 \pm 0.24$<br>$BF_{10} = 1.67$ | $\overline{\beta} = 0.58 \pm 0.31$<br>$BF_{10} = 0.99$ | $\overline{\beta} = 0.09 \pm 0.14$<br>$BF_{10} = 0.28$ | $\overline{\beta} = 0.25 \pm 0.24$<br>$BF_{10} = 0.37$ |
| 12 (18) | $\overline{\beta} = 0.59 \pm 0.95$<br>$BF_{10} = 0.29$ | $\overline{\beta} = -0.56 \pm 0.22$<br>$BF_{10} = 2.97$ | $\overline{\beta} = 0.05 \pm 0.17$<br>$BF_{10} = 0.25$ | $\overline{\beta} = 0.97 \pm 0.49$<br>$BF_{10} = 1.20$ | $\overline{\beta} = 0.49 \pm 0.21$<br>$BF_{10} = 1.91$ | $\overline{\beta} = 0.75 \pm 0.36$<br>$BF_{10} = 1.42$ |
| 13 (17) | $\overline{\beta} = -0.65 \pm 0.77$<br>$BF_{10} = 0.34$ | $\overline{\beta} = -0.87 \pm 0.55$<br>$BF_{10} = 0.72$ | $\overline{\beta} = -0.33 \pm 0.42$<br>$BF_{10} = 0.32$ | $\overline{\beta} = 0.02 \pm 0.53$<br>$BF_{10} = 0.25$ | $\overline{\beta} = 0.10 \pm 0.14$<br>$BF_{10} = 0.31$ | $\overline{\beta} = 0.24 \pm 0.50$<br>$BF_{10} = 0.28$ |
| 15 (23) | $\overline{\beta} = 1.01 \pm 0.80$<br>$BF_{10} = 0.44$ | $\overline{\beta} = -0.16 \pm 0.26$<br>$BF_{10} = 0.26$ | $\overline{\beta} = 0.53 \pm 0.20$<br>$\mathbf{BF_{10} = 4.22}$ | $\overline{\beta} = 0.62 \pm 0.29$<br>$BF_{10} = 1.49$ | $\overline{\beta} = 0.51 \pm 0.16$<br>$\mathbf{BF_{10} = 9.35}$ | $\overline{\beta} = 0.79 \pm 0.34$<br>$BF_{10} = 1.92$ |
| 17 (17) | $\overline{\beta} = 0.36 \pm 1.02$<br>$BF_{10} = 0.26$ | $\overline{\beta} = -0.79 \pm 0.44$<br>$BF_{10} = 0.94$ | $\overline{\beta} = -1.60 \pm 0.43$<br>$\mathbf{BF_{10} = 21.56}$ | $\overline{\beta} = 0.52 \pm 0.34$<br>$BF_{10} = 0.67$ | $\overline{\beta} = 0.15 \pm 0.21$<br>$BF_{10} = 0.31$ | $\overline{\beta} = 0.60 \pm 0.37$<br>$BF_{10} = 0.73$ |
| 18 (22) | $\overline{\beta} = -1.07 \pm 0.0.97$<br>$BF_{10} = 0.38$ | $\overline{\beta} = -1.75 \pm 0.32$<br>$\mathbf{BF_{10} = 1321.84}$ | $\overline{\beta} = -2.50 \pm 0.24$<br>$\mathbf{BF_{10} = 9.20 \cdot 10^{6}}$ | $\overline{\beta} = 0.17 \pm 0.38$<br>$BF_{10} = 0.24$ | $\overline{\beta} = 0.10 \pm 0.28$<br>$BF_{10} = 0.28$ | $\overline{\beta} = -0.39 \pm 0.39$<br>$BF_{10} = 0.35$ |
| 19 (20) | $\overline{\beta} = -1.05 \pm 0.0.95$<br>$BF_{10} = 0.40$ | $\overline{\beta} = -1.83 \pm 0.51$<br>$\mathbf{BF_{10} = 20.41}$ | $\overline{\beta} = -1.21 \pm 0.30$<br>$\mathbf{BF_{10} = 51.04}$ | $\overline{\beta} = -0.07 \pm 0.34$<br>$BF_{10} = 0.24$ | $\overline{\beta} = 0.30 \pm 0.11$<br>$\mathbf{BF_{10} = 3.72}$ | $\overline{\beta} = -0.19 \pm 0.26$<br>$BF_{10} = 0.29$ |
| 21 (23) | $\overline{\beta} = 0.35 \pm 0.79$<br>$BF_{10} = 0.24$ | $\overline{\beta} = 0.06 \pm 0.37$<br>$BF_{10} = 0.22$ | $\overline{\beta} = 0.49 \pm 0.31$<br>$BF_{10} = 0.62$ | $\overline{\beta} = 0.38 \pm 0.33$<br>$BF_{10} = 0.40$ | $\overline{\beta} = 0.18 \pm 0.21$<br>$BF_{10} = 0.31$ | $\overline{\beta} = 0.34 \pm 0.34$<br>$BF_{10} = 0.35$ |
| 22 (21) | $\overline{\beta} = -0.29 \pm 0.92$<br>$BF_{10} = 0.24$ | $\overline{\beta} = -0.86 \pm 0.40$<br>$BF_{10} = 1.43$ | $\overline{\beta} = -0.24 \pm 0.23$<br>$BF_{10} = 0.37$ | $\overline{\beta} = 0.53 \pm 0.43$<br>$BF_{10} = 0.44$ | $\overline{\beta} = 0.16 \pm 0.21$<br>$BF_{10} = 0.29$ | $\overline{\beta} = 0.07 \pm 0.23$<br>$BF_{10} = 0.24$ |

**Table 22** Results of the Bayesian t-tests of the BETA MAPS analysis of the SIM $\Delta[HbR]$ data. Mean $\pm$ SEM represent mean beta values of the respective channel and its standard error of the mean across participants, $BF_{10}$ represent Bayes factors. Bayesian t-tests were performed with the *BayesFactor* package in R for which the default prior odds is set at $P(M) = \frac{\sqrt{2}}{2}$. Corresponding BETA MAPS are visualized in Figure 6 B of the main document.

| | SIM | | | | | |
|---|---|---|---|---|---|---|
| Channel (Channel Frequency after Pruning) | **NO SAC** | **CAR** | **GCR** | **SSR** | **GLM ALL** | **GLM BH** |
| 1 (20) | $\overline{\beta} = 0.02 \pm 0.10$ $BF_{10} = 0.24$ | $\overline{\beta} = 0.27 \pm 0.08$ $\mathbf{BF_{10} = 19.09}$ | $\overline{\beta} = 0.09 \pm 0.06$ $BF_{10} = 0.58$ | $\overline{\beta} = 0.02 \pm 0.09$ $BF_{10} = 0.24$ | $\overline{\beta} = -0.07 \pm 0.05$ $BF_{10} = 0.57$ | $\overline{\beta} = 0.00 \pm 0.05$ $BF_{10} = 0.23$ |
| 2 (19) | $\overline{\beta} = -0.12 \pm 0.30$ $BF_{10} = 0.25$ | $\overline{\beta} = 0.12 \pm 0.29$ $BF_{10} = 0.26$ | $\overline{\beta} = 0.25 \pm 0.18$ $BF_{10} = 0.51$ | $\overline{\beta} = -0.13 \pm 0.32$ $BF_{10} = 0.26$ | $\overline{\beta} = -0.12 \pm 0.14$ $BF_{10} = 0.33$ | $\overline{\beta} = -0.09 \pm 0.31$ $BF_{10} = 0.25$ |
| 4 (19) | $\overline{\beta} = 0.13 \pm 0.13$ $BF_{10} = 0.37$ | $\overline{\beta} = 0.40 \pm 0.08$ $\mathbf{BF_{10} = 276.64}$ | $\overline{\beta} = 0.20 \pm 0.05$ $\mathbf{BF_{10} = 35.93}$ | $\overline{\beta} = 0.12 \pm 0.09$ $BF_{10} = 0.56$ | $\overline{\beta} = -0.04 \pm 0.05$ $BF_{10} = 0.34$ | $\overline{\beta} = 0.03 \pm 0.06$ $BF_{10} = 0.28$ |
| 6 (23) | $\overline{\beta} = -2.47 \pm 0.15$ $\mathbf{BF_{10} = 6.70 \cdot 10^{10}}$ | $\overline{\beta} = -2.25 \pm 0.13$ $\mathbf{BF_{10} = 3.36 \cdot 10^{11}}$ | $\overline{\beta} = -1.73 \pm 0.12$ $\mathbf{BF_{10} = 4.63 \cdot 10^{9}}$ | $\overline{\beta} = -2.46 \pm 0.15$ $\mathbf{BF_{10} = 9.50 \cdot 10^{10}}$ | $\overline{\beta} = -2.15 \pm 0.15$ $\mathbf{BF_{10} = 7.19 \cdot 10^{9}}$ | $\overline{\beta} = -2.57 \pm 0.15$ $\mathbf{BF_{10} = 2.02 \cdot 10^{11}}$ |
| 7 (18) | $\overline{\beta} = 0.00 \pm 0.18$ $BF_{10} = 0.24$ | $\overline{\beta} = 0.24 \pm 0.11$ $BF_{10} = 1.84$ | $\overline{\beta} = 0.61 \pm 0.08$ $\mathbf{BF_{10} = 1.47 \cdot 10^{4}}$ | $\overline{\beta} = 0.04 \pm 0.18$ $BF_{10} = 0.25$ | $\overline{\beta} = -0.10 \pm 0.12$ $BF_{10} = 0.33$ | $\overline{\beta} = -0.04 \pm 0.16$ $BF_{10} = 0.25$ |
| 8 (23) | $\overline{\beta} = -1.16 \pm 0.14$ $\mathbf{BF_{10} = 4.96 \cdot 10^{5}}$ | $\overline{\beta} = -10.97 \pm 0.13$ $\mathbf{BF_{10} = 5.88 \cdot 10^{4}}$ | $\overline{\beta} = -0.51 \pm 0.10$ $\mathbf{BF_{10} = 415.47}$ | $\overline{\beta} = -1.36 \pm 0.15$ $\mathbf{BF_{10} = 1.25 \cdot 10^{6}}$ | $\overline{\beta} = -1.00 \pm 0.10$ $\mathbf{BF_{10} = 4.74 \cdot 10^{6}}$ | $\overline{\beta} = -1.30 \pm 0.12$ $\mathbf{BF_{10} = 2.04 \cdot 10^{7}}$ |
| 10 (13) | $\overline{\beta} = 0.25 \pm 0.14$ $BF_{10} = 1.04$ | $\overline{\beta} = 0.48 \pm 0.14$ $\mathbf{BF_{10} = 9.25}$ | $\overline{\beta} = 0.17 \pm 0.13$ $BF_{10} = 0.55$ | $\overline{\beta} = 0.27 \pm 0.15$ $BF_{10} = 1.04$ | $\overline{\beta} = 0.11 \pm 0.09$ $BF_{10} = 0.56$ | $\overline{\beta} = 0.22 \pm 0.11$ $BF_{10} = 1.23$ |
| 11 (20) | $\overline{\beta} = 0.18 \pm 0.15$ $BF_{10} = 0.43$ | $\overline{\beta} = 0.41 \pm 0.07$ $\mathbf{BF_{10} = 2428.17}$ | $\overline{\beta} = 0.16 \pm 0.06$ $\mathbf{BF_{10} = 3.12}$ | $\overline{\beta} = 0.14 \pm 0.10$ $BF_{10} = 0.52$ | $\overline{\beta} = -0.06 \pm 0.06$ $BF_{10} = 0.37$ | $\overline{\beta} = 0.05 \pm 0.09$ $BF_{10} = 0.27$ |
| 12 (18) | $\overline{\beta} = -0.18 \pm 0.16$ $BF_{10} = 0.42$ | $\overline{\beta} = 0.07 \pm 0.08$ $BF_{10} = 0.33$ | $\overline{\beta} = -0.18 \pm 0.06$ $\mathbf{BF_{10} = 10.78}$ | $\overline{\beta} = -0.34 \pm 0.12$ $\mathbf{BF_{10} = 5.04}$ | $\overline{\beta} = -0.15 \pm 0.07$ $BF_{10} = 1.35$ | $\overline{\beta} = -0.19 \pm 0.09$ $BF_{10} = 1.51$ |
| 13 (17) | $\overline{\beta} = 0.17 \pm 0.15$ $BF_{10} = 0.42$ | $\overline{\beta} = 0.43 \pm 0.12$ $\mathbf{BF_{10} = 19.55}$ | $\overline{\beta} = 0.17 \pm 0.10$ $BF_{10} = 0.83$ | $\overline{\beta} = 0.24 \pm 0.23$ $BF_{10} = 0.39$ | $\overline{\beta} = 0.03 \pm 0.05$ $BF_{10} = 0.28$ | $\overline{\beta} = 0.29 \pm 0.21$ $BF_{10} = 0.53$ |
| 15 (23) | $\overline{\beta} = 0.10 \pm 0.12$ $BF_{10} = 0.29$ | $\overline{\beta} = 0.31 \pm 0.09$ $\mathbf{BF_{10} = 11.06}$ | $\overline{\beta} = 0.05 \pm 0.07$ $BF_{10} = 0.28$ | $\overline{\beta} = 0.10 \pm 0.10$ $BF_{10} = 0.34$ | $\overline{\beta} = -0.04 \pm 0.07$ $BF_{10} = 0.26$ | $\overline{\beta} = -0.05 \pm 0.10$ $BF_{10} = 0.25$ |
| 17 (17) | $\overline{\beta} = 0.09 \pm 0.16$ $BF_{10} = 0.29$ | $\overline{\beta} = 0.22 \pm 0.08$ $BF_{10} = 3.00$ | $\overline{\beta} = 0.53 \pm 0.08$ $\mathbf{BF_{10} = 3126.04}$ | $\overline{\beta} = 0.11 \pm 0.13$ $BF_{10} = 0.35$ | $\overline{\beta} = 0.02 \pm 0.07$ $BF_{10} = 0.25$ | $\overline{\beta} = 0.06 \pm 0.08$ $BF_{10} = 0.31$ |
| 18 (22) | $\overline{\beta} = -0.05 \pm 0.13$ $BF_{10} = 0.24$ | $\overline{\beta} = 0.21 \pm 0.09$ $BF_{10} = 1.88$ | $\overline{\beta} = 0.55 \pm 0.06$ $\mathbf{BF_{10} = 1.09 \cdot 10^{6}}$ | $\overline{\beta} = -0.06 \pm 0.13$ $BF_{10} = 0.24$ | $\overline{\beta} = -0.05 \pm 0.06$ $BF_{10} = 0.30$ | $\overline{\beta} = -0.04 \pm 0.08$ $BF_{10} = 0.25$ |
| 19 (20) | $\overline{\beta} = 0.20 \pm 0.12$ $BF_{10} = 0.72$ | $\overline{\beta} = 0.50 \pm 0.11$ $\mathbf{BF_{10} = 188.58}$ | $\overline{\beta} = 0.33 \pm 0.08$ $\mathbf{BF_{10} = 55.39}$ | $\overline{\beta} = 0.15 \pm 0.11$ $BF_{10} = 0.51$ | $\overline{\beta} = -0.01 \pm 0.07$ $BF_{10} = 0.23$ | $\overline{\beta} = 0.10 \pm 0.11$ $BF_{10} = 0.34$ |
| 21 (23) | $\overline{\beta} = -0.09 \pm 0.13$ $BF_{10} = 0.27$ | $\overline{\beta} = 0.21 \pm 0.07$ $\mathbf{BF_{10} = 5.74}$ | $\overline{\beta} = -0.10 \pm 0.06$ $BF_{10} = 0.63$ | $\overline{\beta} = -0.11 \pm 0.11$ $BF_{10} = 0.32$ | $\overline{\beta} = -0.08 \pm 0.07$ $BF_{10} = 0.39$ | $\overline{\beta} = -0.19 \pm 0.10$ $BF_{10} = 1.10$ |
| 22 (21) | $\overline{\beta} = 0.05 \pm 0.14$ $BF_{10} = 0.24$ | $\overline{\beta} = 0.23 \pm 0.07$ $\mathbf{BF_{10} = 8.18}$ | $\overline{\beta} = -0.01 \pm 0.05$ $BF_{10} = 0.23$ | $\overline{\beta} = 0.06 \pm 0.11$ $BF_{10} = 0.26$ | $\overline{\beta} = 0.09 \pm 0.08$ $BF_{10} = 0.39$ | $\overline{\beta} = 0.07 \pm 0.11$ $BF_{10} = 0.27$ |

**Table 23** Results of the Bayesian t-tests of the BETA MAPS analysis of the ME LEFT and ME RIGHT $\Delta[HbO]$ data. Mean ± SEM represent mean beta values of the respective channel and its standard error of the mean across participants, $BF_{10}$ represent Bayes factors. Bayesian t-tests were performed with the *BayesFactor* package in R for which the default prior odds is set at $P(M) = \frac{\sqrt{2}}{2}$. Corresponding BETA MAPS are visualized in Figure 8 B of the main document.

**ME LEFT**

| Channel (Channel Frequency after Pruning) | **NO SAC** | **CAR** | **GCR** | **SSR** | **GLM ALL** | **GLM BH** |
|---|---|---|---|---|---|---|
| 1 (23) | $\beta = 1.91 \pm 0.67$<br>$\mathbf{BF_{10} = 5.04}$ | $\beta = -2.02 \pm 0.41$<br>$\mathbf{BF_{10} = 376.91}$ | $\beta = -1.39 \pm 0.30$<br>$\mathbf{BF_{10} = 249.65}$ | $\beta = 0.56 \pm 0.24$<br>$BF_{10} = 2.16$ | $\beta = -0.21 \pm 0.17$<br>$BF_{10} = 0.43$ | $\beta = -0.29 \pm 0.24$<br>$BF_{10} = 0.42$ |
| 2 (16) | $\beta = 3.12 \pm 0.93$<br>$\mathbf{BF_{10} = 10.92}$ | $\beta = -0.76 \pm 0.52$<br>$BF_{10} = 0.62$ | $\beta = -0.39 \pm 0.43$<br>$BF_{10} = 0.36$ | $\beta = 1.19 \pm 0.47$<br>$BF_{10} = 2.64$ | $\beta = 0.56 \pm 0.33$<br>$BF_{10} = 0.81$ | $\beta = 0.48 \pm 0.46$<br>$BF_{10} = 0.41$ |
| 4 (17) | $\beta = 3.40 \pm 0.91$<br>$\mathbf{BF_{10} = 22.83}$ | $\beta = -0.80 \pm 0.24$<br>$\mathbf{BF_{10} = 9.82}$ | $\beta = -0.29 \pm 0.17$<br>$BF_{10} = 0.89$ | $\beta = 0.73 \pm 0.21$<br>$\mathbf{BF_{10} = 15.76}$ | $\beta = 0.40 \pm 0.13$<br>$\mathbf{BF_{10} = 7.03}$ | $\beta = 0.39 \pm 0.17$<br>$BF_{10} = 1.91$ |
| 6 (24) | $\beta = 3.01 \pm 0.71$<br>$\mathbf{BF_{10} = 96.51}$ | $\beta = -0.71 \pm 0.42$<br>$BF_{10} = 0.75$ | $\beta = 0.01 \pm 0.11$<br>$BF_{10} = 0.22$ | $\beta = -0.69 \pm 0.15$<br>$\mathbf{BF_{10} = 226.55}$ | $\beta = -0.46 \pm 0.10$<br>$\mathbf{BF_{10} = 221.94}$ | $\beta = -0.58 \pm 0.13$<br>$\mathbf{BF_{10} = 159.51}$ |
| 7 (19) | $\beta = 5.10 \pm 0.86$<br>$\mathbf{BF_{10} = 1752.49}$ | $\beta = 0.62 \pm 0.34$<br>$BF_{10} = 0.95$ | $\beta = 0.78 \pm 0.27$<br>$\mathbf{BF_{10} = 5.35}$ | $\beta = 2.27 \pm 0.33$<br>$\mathbf{BF_{10} = 10.77 \cdot 10^3}$ | $\beta = 1.55 \pm 0.29$<br>$\mathbf{BF_{10} = 561.50}$ | $\beta = 2.01 \pm 0.40$<br>$\mathbf{BF_{10} = 318.17}$ |
| 8 (23) | $\beta = 4.34 \pm 0.94$<br>$\mathbf{BF_{10} = 200.72}$ | $\beta = 0.24 \pm 0.40$<br>$BF_{10} = 0.26$ | $\beta = 0.28 \pm 0.36$<br>$BF_{10} = 0.29$ | $\beta = 1.36 \pm 0.38$<br>$\mathbf{BF_{10} = 20.57}$ | $\beta = 0.97 \pm 0.35$<br>$\mathbf{BF_{10} = 4.46}$ | $\beta = 1.45 \pm 0.50$<br>$\mathbf{BF_{10} = 5.63}$ |
| 10 (17) | $\beta = 3.88 \pm 0.83$<br>$\mathbf{BF_{10} = 131.36}$ | $\beta = -0.07 \pm 0.43$<br>$BF_{10} = 0.25$ | $\beta = 0.19 \pm 0.33$<br>$BF_{10} = 0.29$ | $\beta = 1.51 \pm 0.40$<br>$\mathbf{BF_{10} = 26.13}$ | $\beta = 1.05 \pm 0.35$<br>$\mathbf{BF_{10} = 6.57}$ | $\beta = 0.98 \pm 0.42$<br>$BF_{10} = 2.03$ |
| 11 (20) | $\beta = 4.20 \pm 0.92$<br>$\mathbf{BF_{10} = 149.48}$ | $\beta = 0.36 \pm 0.23$<br>$BF_{10} = 0.64$ | $\beta = 0.81 \pm 0.30$<br>$\mathbf{BF_{10} = 4.35}$ | $\beta = 1.25 \pm 0.26$<br>$\mathbf{BF_{10} = 207.60}$ | $\beta = 0.77 \pm 0.18$<br>$\mathbf{BF_{10} = 93.81}$ | $\beta = 1.02 \pm 0.23$<br>$\mathbf{BF_{10} = 111.29}$ |
| 12 (19) | $\beta = 4.31 \pm 0.91$<br>$\mathbf{BF_{10} = 175.62}$ | $\beta = 0.66 \pm 0.35$<br>$BF_{10} = 1.02$ | $\beta = 0.22 \pm 0.29$<br>$BF_{10} = 0.31$ | $\beta = 1.75 \pm 0.50$<br>$\mathbf{BF_{10} = 17.17}$ | $\beta = 1.45 \pm 0.30$<br>$\mathbf{BF_{10} = 238.65}$ | $\beta = 1.46 \pm 0.38$<br>$\mathbf{BF_{10} = 29.79}$ |
| 13 (17) | $\beta = 4.13 \pm 1.03$<br>$\mathbf{BF_{10} = 37.12}$ | $\beta = 0.42 \pm 0.29$<br>$BF_{10} = 0.60$ | $\beta = 0.02 \pm 0.28$<br>$BF_{10} = 0.25$ | $\beta = 1.50 \pm 0.44$<br>$\mathbf{BF_{10} = 12.86}$ | $\beta = 1.01 \pm 0.23$<br>$\mathbf{BF_{10} = 82.38}$ | $\beta = 1.40 \pm 0.24$<br>$\mathbf{BF_{10} = 1150.48}$ |
| 15 (23) | $\beta = 4.01 \pm 0.74$<br>$\mathbf{BF_{10} = 1214.67}$ | $\beta = 0.14 \pm 0.37$<br>$BF_{10} = 0.23$ | $\beta = -0.42 \pm 0.27$<br>$BF_{10} = 0.62$ | $\beta = 3.17 \pm 0.54$<br>$\mathbf{BF_{10} = 3.03 \cdot 10^3}$ | $\beta = 1.75 \pm 0.37$<br>$\mathbf{BF_{10} = 241.18}$ | $\beta = 2.30 \pm 0.52$<br>$\mathbf{BF_{10} = 131.99}$ |
| 17 (20) | $\beta = 5.15 \pm 1.05$<br>$\mathbf{BF_{10} = 281.44}$ | $\beta = 0.83 \pm 0.59$<br>$BF_{10} = 0.55$ | $\beta = 0.77 \pm 0.53$<br>$BF_{10} = 0.58$ | $\beta = 2.07 \pm 0.60$<br>$\mathbf{BF_{10} = 15.86}$ | $\beta = 1.08 \pm 0.44$<br>$BF_{10} = 2.45$ | $\beta = 1.57 \pm 0.47$<br>$\mathbf{BF_{10} = 12.27}$ |
| 18 (21) | $\beta = 3.70 \pm 0.94$<br>$\mathbf{BF_{10} = 44.28}$ | $\beta = -0.44 \pm 0.44$<br>$BF_{10} = 0.35$ | $\beta = -0.44 \pm 0.34$<br>$BF_{10} = 0.47$ | $\beta = 0.66 \pm 0.27$<br>$BF_{10} = 2.28$ | $\beta = 0.00 \pm 0.17$<br>$BF_{10} = 0.23$ | $\beta = 0.33 \pm 0.27$<br>$BF_{10} = 0.44$ |
| 19 (20) | $\beta = 3.04 \pm 1.03$<br>$\mathbf{BF_{10} = 6.00}$ | $\beta = -1.07 \pm 0.48$<br>$BF_{10} = 1.68$ | $\beta = -0.95 \pm 0.31$<br>$\mathbf{BF_{10} = 8.15}$ | $\beta = 0.14 \pm 0.24$<br>$BF_{10} = 0.27$ | $\beta = -0.04 \pm 0.18$<br>$BF_{10} = 0.24$ | $\beta = 0.03 \pm 0.17$<br>$BF_{10} = 0.24$ |
| 21 (23) | $\beta = 5.74 \pm 1.14$<br>$\mathbf{BF_{10} = 503.95}$ | $\beta = 1.86 \pm 0.59$<br>$\mathbf{BF_{10} = 9.27}$ | $\beta = 1.26 \pm 0.40$<br>$\mathbf{BF_{10} = 10.14}$ | $\beta = 3.28 \pm 0.72$<br>$\mathbf{BF_{10} = 188.73}$ | $\beta = 2.79 \pm 0.58$<br>$\mathbf{BF_{10} = 335.61}$ | $\beta = 3.06 \pm 0.69$<br>$\mathbf{BF_{10} = 150.44}$ |
| 22 (21) | $\beta = 4.53 \pm 1.00$<br>$\mathbf{BF_{10} = 143.36}$ | $\beta = 0.39 \pm 0.51$<br>$BF_{10} = 0.29$ | $\beta = -0.01 \pm 0.38$<br>$BF_{10} = 0.23$ | $\beta = 1.68 \pm 0.44$<br>$\mathbf{BF_{10} = 32.79}$ | $\beta = 1.31 \pm 0.39$<br>$\mathbf{BF_{10} = 12.98}$ | $\beta = 1.23 \pm 0.53$<br>$BF_{10} = 2.01$ |

**ME RIGHT**

| Channel (Channel Frequency after Pruning) | **NO SAC** | CAR | **GCR** | **SSR** | **GLM ALL** | **GLM BH** |
|---|---|---|---|---|---|---|
| 1 (23) | $\beta = 1.62 \pm 0.62$<br>$\mathbf{BF_{10} = 3.32}$ | $\beta = -1.61 \pm 0.49$<br>$\mathbf{BF_{10} = 12.46}$ | $\beta = -1.05 \pm 0.31$<br>$\mathbf{BF_{10} = 14.42}$ | $\beta = 0.38 \pm 0.28$<br>$BF_{10} = 0.50$ | $\beta = -0.22 \pm 0.17$<br>$BF_{10} = 0.47$ | $\beta = -0.18 \pm 0.30$<br>$BF_{10} = 0.26$ |
| 2 (16) | $\beta = 2.86 \pm 0.95$<br>$\mathbf{BF_{10} = 6.15}$ | $\beta = -0.13 \pm 0.60$<br>$BF_{10} = 0.26$ | $\beta = -0.29 \pm 0.48$<br>$BF_{10} = 0.30$ | $\beta = 1.01 \pm 0.51$<br>$BF_{10} = 1.20$ | $\beta = 0.42 \pm 0.58$<br>$BF_{10} = 0.32$ | $\beta = 0.79 \pm 0.62$<br>$BF_{10} = 0.51$ |
| 4 (17) | $\beta = 2.76 \pm 0.61$<br>$\mathbf{BF_{10} = 100.62}$ | $\beta = -0.55 \pm 0.21$<br>$\mathbf{BF_{10} = 3.55}$ | $\beta = -0.31 \pm 0.19$<br>$BF_{10} = 0.80$ | $\beta = 0.71 \pm 0.22$<br>$\mathbf{BF_{10} = 8.00}$ | $\beta = 0.10 \pm 0.15$<br>$BF_{10} = 0.30$ | $\beta = 0.41 \pm 0.18$<br>$BF_{10} = 1.82$ |
| 6 (24) | $\beta = 3.04 \pm 0.61$<br>$\mathbf{BF_{10} = 488.28}$ | $\beta = 0.04 \pm 0.36$<br>$BF_{10} = 0.22$ | $\beta = -0.65 \pm 0.32$<br>$BF_{10} = 1.29$ | $\beta = 1.78 \pm 0.45$<br>$\mathbf{BF_{10} = 53.34}$ | $\beta = 1.17 \pm 0.38$<br>$\mathbf{BF_{10} = 8.03}$ | $\beta = 1.43 \pm 0.46$<br>$BF_{10} = 8.84$ |
| 7 (19) | $\beta = 4.54 \pm 0.65$<br>$\mathbf{BF_{10} = 11.68 \cdot 10^3}$ | $\beta = 1.23 \pm 0.31$<br>$\mathbf{BF_{10} = 38.87}$ | $\beta = 0.58 \pm 0.31$<br>$BF_{10} = 1.01$ | $\beta = 2.58 \pm 0.32$<br>$\mathbf{BF_{10} = 77.57 \cdot 10^3}$ | $\beta = 2.09 \pm 0.30$<br>$\mathbf{BF_{10} = 11.47 \cdot 10^3}$ | $\beta = 2.31 \pm 0.40$<br>$\mathbf{BF_{10} = 1.35 \cdot 10^3}$ |
| 8 (23) | $\beta = 4.59 \pm 0.64$<br>$\mathbf{BF_{10} = 43.09 \cdot 10^3}$ | $\beta = 1.39 \pm 0.41$<br>$\mathbf{BF_{10} = 16.25}$ | $\beta = 0.75 \pm 0.34$<br>$BF_{10} = 1.69$ | $\beta = 2.62 \pm 0.43$<br>$\mathbf{BF_{10} = 4.49 \cdot 10^3}$ | $\beta = 2.07 \pm 0.48$<br>$\mathbf{BF_{10} = 105.78}$ | $\beta = 2.47 \pm 0.52$<br>$\mathbf{BF_{10} = 299.58}$ |
| 10 (17) | $\beta = 2.85 \pm 0.46$<br>$\mathbf{BF_{10} = 1.84 \cdot 10^3}$ | $\beta = -0.11 \pm 0.37$<br>$BF_{10} = 0.26$ | $\beta = 0.23 \pm 0.24$<br>$BF_{10} = 0.37$ | $\beta = 1.29 \pm 0.39$<br>$\mathbf{BF_{10} = 10.15}$ | $\beta = 0.22 \pm 0.21$<br>$BF_{10} = 0.40$ | $\beta = 0.76 \pm 0.30$<br>$BF_{10} = 2.92$ |
| 11 (20) | $\beta = 3.30 \pm 0.54$<br>$\mathbf{BF_{10} = 3.19 \cdot 10^3}$ | $\beta = 0.36 \pm 0.16$<br>$BF_{10} = 1.85$ | $\beta = 0.59 \pm 0.19$<br>$\mathbf{BF_{10} = 7.11}$ | $\beta = 1.62 \pm 0.33$<br>$\mathbf{BF_{10} = 279.53}$ | $\beta = 0.79 \pm 0.20$<br>$\mathbf{BF_{10} = 40.48}$ | $\beta = 1.29 \pm 0.26$<br>$\mathbf{BF_{10} = 323.23}$ |
| 12 (19) | $\beta = 2.92 \pm 0.62$<br>$\mathbf{BF_{10} = 171.55}$ | $\beta = 0.09 \pm 0.31$<br>$BF_{10} = 0.25$ | $\beta = 0.57 \pm 0.25$<br>$BF_{10} = 1.83$ | $\beta = 1.17 \pm 0.37$<br>$\mathbf{BF_{10} = 9.55}$ | $\beta = 0.69 \pm 0.25$<br>$\mathbf{BF_{10} = 4.02}$ | $\beta = 0.78 \pm 0.33$<br>$BF_{10} = 2.00$ |
| 13 (17) | $\beta = 2.56 \pm 0.66$<br>$\mathbf{BF_{10} = 29.33}$ | $\beta = -0.29 \pm 0.33$<br>$BF_{10} = 0.35$ | $\beta = 0.19 \pm 0.29$<br>$BF_{10} = 0.30$ | $\beta = 0.66 \pm 0.25$<br>$\mathbf{BF_{10} = 3.14}$ | $\beta = 0.10 \pm 0.23$<br>$BF_{10} = 0.27$ | $\beta = 0.53 \pm 0.17$<br>$\mathbf{BF_{10} = 7.96}$ |
| 15 (23) | $\beta = 2.42 \pm 0.65$<br>$\mathbf{BF_{10} = 31.62}$ | $\beta = -0.67 \pm 0.33$<br>$BF_{10} = 1.26$ | $\beta = -0.10 \pm 0.28$<br>$BF_{10} = 0.23$ | $\beta = 1.82 \pm 0.52$<br>$\mathbf{BF_{10} = 19.24}$ | $\beta = 0.84 \pm 0.37$<br>$BF_{10} = 1.96$ | $\beta = 0.86 \pm 0.45$<br>$BF_{10} = 1.02$ |
| 17 (20) | $\beta = 4.65 \pm 0.76$<br>$\mathbf{BF_{10} = 2.90 \cdot 10^3}$ | $\beta = 1.51 \pm 0.81$<br>$BF_{10} = 0.99$ | $\beta = 0.66 \pm 0.66$<br>$BF_{10} = 0.36$ | $\beta = 2.55 \pm 0.83$<br>$\mathbf{BF_{10} = 7.81}$ | $\beta = 2.06 \pm 0.61$<br>$\mathbf{BF_{10} = 14.00}$ | $\beta = 2.42 \pm 0.68$<br>$\mathbf{BF_{10} = 19.54}$ |
| 18 (21) | $\beta = 3.96 \pm 0.56$<br>$\mathbf{BF_{10} = 25.38 \cdot 10^3}$ | $\beta = 0.70 \pm 0.30$<br>$BF_{10} = 2.07$ | $\beta = -0.13 \pm 0.37$<br>$BF_{10} = 0.24$ | $\beta = 1.94 \pm 0.38$<br>$\mathbf{BF_{10} = 498.90}$ | $\beta = 1.39 \pm 0.33$<br>$\mathbf{BF_{10} = 83.07}$ | $\beta = 1.76 \pm 0.40$<br>$\mathbf{BF_{10} = 106.95}$ |
| 19 (20) | $\beta = 2.30 \pm 0.67$<br>$\mathbf{BF_{10} = 14.96}$ | $\beta = -0.52 \pm 0.40$<br>$BF_{10} = 0.48$ | $\beta = -0.54 \pm 0.26$<br>$BF_{10} = 1.30$ | $\beta = 0.28 \pm 0.22$<br>$BF_{10} = 0.46$ | $\beta = 0.03 \pm 0.22$<br>$BF_{10} = 0.23$ | $\beta = 0.11 \pm 0.27$<br>$BF_{10} = 0.25$ |
| 21 (23) | $\beta = 2.40 \pm 0.63$<br>$\mathbf{BF_{10} = 35.10}$ | $\beta = -0.97 \pm 0.46$<br>$BF_{10} = 1.39$ | $\beta = -0.24 \pm 0.33$<br>$BF_{10} = 0.28$ | $\beta = 1.01 \pm 0.35$<br>$\mathbf{BF_{10} = 5.96}$ | $\beta = 0.38 \pm 0.38$<br>$BF_{10} = 0.34$ | $\beta = 0.23 \pm 0.54$<br>$BF_{10} = 0.24$ |
| 22 (21) | $\beta = 2.35 \pm 0.75$<br>$\mathbf{BF_{10} = 8.61}$ | $\beta = -0.94 \pm 0.42$<br>$BF_{10} = 1.73$ | $\beta = -0.34 \pm 0.33$<br>$BF_{10} = 0.36$ | $\beta = 0.86 \pm 0.43$<br>$BF_{10} = 1.24$ | $\beta = -0.05 \pm 0.23$<br>$BF_{10} = 0.23$ | $\beta = -0.20 \pm 0.36$<br>$BF_{10} = 0.26$ |

**Table 24** Results of the Bayesian t-tests of the BETA MAPS analysis of the ME LEFT and ME RIGHT $\Delta[HbR]$ data. Mean $\pm$ SEM represent mean beta values of the respective channel and its standard error of the mean across participants, $BF_{10}$ represent Bayes factors. Bayesian t-tests were performed with the *BayesFactor* package in R for which the default prior odds is set at $P(M) = \frac{\sqrt{2}}{2}$. Corresponding BETA MAPS are visualized in Figure 8 B of the main document.

**ME LEFT**

| Channel (Channel Frequency after Pruning) | **NO SAC** | **CAR** | **GCR** | **SSR** | **GLM ALL** | **GLM BH** |
|---|---|---|---|---|---|---|
| 1 (23) | $\beta = -0.11 \pm 0.06$<br>$BF_{10} = 0.82$ | $\beta = 0.74 \pm 0.15$<br>$\mathbf{BF_{10} = 445.02}$ | $\beta = 0.56 \pm 0.11$<br>$\mathbf{BF_{10} = 579.87}$ | $\beta = -0.08 \pm 0.06$<br>$BF_{10} = 0.44$ | $\beta = -0.11 \pm 0.07$<br>$BF_{10} = 0.71$ | $\beta = -0.16 \pm 0.07$<br>$BF_{10} = 2.27$ |
| 2 (16) | $\beta = -0.51 \pm 0.17$<br>$\mathbf{BF_{10} = 5.78}$ | $\beta = 0.21 \pm 0.14$<br>$BF_{10} = 0.63$ | $\beta = 0.09 \pm 0.11$<br>$BF_{10} = 0.33$ | $\beta = -0.44 \pm 0.17$<br>$BF_{10} = 3.29$ | $\beta = -0.44 \pm 0.17$<br>$BF_{10} = 3.06$ | $\beta = -0.49 \pm 0.18$<br>$\mathbf{BF_{10} = 3.70}$ |
| 4 (17) | $\beta = -0.16 \pm 0.07$<br>$BF_{10} = 2.08$ | $\beta = 0.44 \pm 0.09$<br>$\mathbf{BF_{10} = 117.86}$ | $\beta = 0.29 \pm 0.06$<br>$\mathbf{BF_{10} = 144.66}$ | $\beta = -0.08 \pm 0.07$<br>$BF_{10} = 0.44$ | $\beta = -0.14 \pm 0.06$<br>$BF_{10} = 1.56$ | $\beta = -0.20 \pm 0.10$<br>$BF_{10} = 1.47$ |
| 6 (24) | $\beta = -0.73 \pm 0.15$<br>$\mathbf{BF_{10} = 279.06}$ | $\beta = 0.04 \pm 0.12$<br>$BF_{10} = 0.22$ | $\beta = 0.01 \pm 0.11$<br>$BF_{10} = 0.22$ | $\beta = -0.69 \pm 0.15$<br>$\mathbf{BF_{10} = 226.55}$ | $\beta = -0.46 \pm 0.10$<br>$\mathbf{BF_{10} = 221.94}$ | $\beta = -0.58 \pm 0.13$<br>$\mathbf{BF_{10} = 159.51}$ |
| 7 (19) | $\beta = -1.08 \pm 0.18$<br>$\mathbf{BF_{10} = 1572.42}$ | $\beta = -0.35 \pm 0.14$<br>$BF_{10} = 2.56$ | $\beta = -0.39 \pm 0.10$<br>$\mathbf{BF_{10} = 24.85}$ | $\beta = -1.09 \pm 0.18$<br>$\mathbf{BF_{10} = 2897.53}$ | $\beta = -0.81 \pm 0.15$<br>$\mathbf{BF_{10} = 570.89}$ | $\beta = -0.94 \pm 0.16$<br>$\mathbf{BF_{10} = 1709.00}$ |
| 8 (23) | $\beta = -0.77 \pm 0.17$<br>$\mathbf{BF_{10} = 183.92}$ | $\beta = 0.02 \pm 0.12$<br>$BF_{10} = 0.22$ | $\beta = 0.03 \pm 0.12$<br>$BF_{10} = 0.22$ | $\beta = -0.69 \pm 0.16$<br>$\mathbf{BF_{10} = 123.15}$ | $\beta = -0.58 \pm 0.19$<br>$\mathbf{BF_{10} = 8.54}$ | $\beta = -0.67 \pm 0.18$<br>$\mathbf{BF_{10} = 28.84}$ |
| 10 (17) | $\beta = -0.90 \pm 0.23$<br>$\mathbf{BF_{10} = 27.97}$ | $\beta = -0.02 \pm 0.12$<br>$BF_{10} = 0.25$ | $\beta = -0.15 \pm 0.12$<br>$BF_{10} = 0.48$ | $\beta = -0.83 \pm 0.24$<br>$\mathbf{BF_{10} = 13.72}$ | $\beta = -0.55 \pm 0.18$<br>$\mathbf{BF_{10} = 6.13}$ | $\beta = -0.69 \pm 0.24$<br>$\mathbf{BF_{10} = 5.24}$ |
| 11 (20) | $\beta = -0.66 \pm 0.12$<br>$\mathbf{BF_{10} = 1358.85}$ | $\beta = -0.05 \pm 0.08$<br>$BF_{10} = 0.27$ | $\beta = -0.21 \pm 0.08$<br>$\mathbf{BF_{10} = 3.80}$ | $\beta = -0.57 \pm 0.13$<br>$\mathbf{BF_{10} = 103.93}$ | $\beta = -0.41 \pm 0.11$<br>$\mathbf{BF_{10} = 23.03}$ | $\beta = -0.45 \pm 0.12$<br>$\mathbf{BF_{10} = 21.32}$ |
| 12 (19) | $\beta = -0.90 \pm 0.18$<br>$\mathbf{BF_{10} = 370.57}$ | $\beta = -0.31 \pm 0.11$<br>$\mathbf{BF_{10} = 3.72}$ | $\beta = -0.13 \pm 0.09$<br>$BF_{10} = 0.60$ | $\beta = -0.83 \pm 0.18$<br>$\mathbf{BF_{10} = 169.47}$ | $\beta = -0.61 \pm 0.12$<br>$\mathbf{BF_{10} = 538.96}$ | $\beta = -0.77 \pm 0.16$<br>$\mathbf{BF_{10} = 225.63}$ |
| 13 (17) | $\beta = -0.63 \pm 0.17$<br>$\mathbf{BF_{10} = 17.50}$ | $\beta = 0.08 \pm 0.13$<br>$BF_{10} = 0.30$ | $\beta = 0.21 \pm 0.15$<br>$BF_{10} = 0.58$ | $\beta = -0.58 \pm 0.17$<br>$\mathbf{BF_{10} = 11.43}$ | $\beta = -0.45 \pm 0.18$<br>$BF_{10} = 2.58$ | $\beta = -0.52 \pm 0.23$<br>$BF_{10} = 1.91$ |
| 15 (23) | $\beta = -0.86 \pm 0.15$<br>$\mathbf{BF_{10} = 2861.99}$ | $\beta = -0.17 \pm 0.15$<br>$BF_{10} = 0.39$ | $\beta = 0.04 \pm 0.14$<br>$BF_{10} = 0.23$ | $\beta = -1.03 \pm 0.18$<br>$\mathbf{BF_{10} = 2030.39}$ | $\beta = -0.74 \pm 0.11$<br>$\mathbf{BF_{10} = 39.20 \cdot 103}$ | $\beta = -0.88 \pm 0.15$<br>$\mathbf{BF_{10} = 3147.56}$ |
| 17 (20) | $\beta = -1.00 \pm 0.19$<br>$\mathbf{BF_{10} = 507.77}$ | $\beta = -0.16 \pm 0.13$<br>$BF_{10} = 0.44$ | $\beta = -0.17 \pm 0.09$<br>$BF_{10} = 1.09$ | $\beta = -0.88 \pm 0.15$<br>$\mathbf{BF_{10} = 1623.04}$ | $\beta = -0.52 \pm 0.14$<br>$\mathbf{BF_{10} = 35.22}$ | $\beta = -0.67 \pm 0.14$<br>$\mathbf{BF_{10} = 200.45}$ |
| 18 (21) | $\beta = -0.64 \pm 0.11$<br>$\mathbf{BF_{10} = 1134.97}$ | $\beta = 0.13 \pm 0.12$<br>$BF_{10} = 0.39$ | $\beta = 0.10 \pm 0.09$<br>$BF_{10} = 0.37$ | $\beta = -0.57 \pm 0.10$<br>$\mathbf{BF_{10} = 1052.78}$ | $\beta = -0.29 \pm 0.07$<br>$\mathbf{BF_{10} = 42.75}$ | $\beta = -0.47 \pm 0.10$<br>$\mathbf{BF_{10} = 310.54}$ |
| 19 (20) | $\beta = -0.34 \pm 0.11$<br>$\mathbf{BF_{10} = 10.68}$ | $\beta = 0.35 \pm 0.11$<br>$\mathbf{BF_{10} = 9.42}$ | $\beta = 0.31 \pm 0.09$<br>$\mathbf{BF_{10} = 17.93}$ | $\beta = -0.27 \pm 0.08$<br>$\mathbf{BF_{10} = 10.77}$ | $\beta = -0.13 \pm 0.05$<br>$\mathbf{BF_{10} = 4.05}$ | $\beta = -0.23 \pm 0.07$<br>$\mathbf{BF_{10} = 8.36}$ |
| 21 (23) | $\beta = -1.46 \pm 0.30$<br>$\mathbf{BF_{10} = 400.93}$ | $\beta = -0.57 \pm 0.17$<br>$\mathbf{BF_{10} = 15.29}$ | $\beta = -0.43 \pm 0.13$<br>$\mathbf{BF_{10} = 11.27}$ | $\beta = -1.32 \pm 0.25$<br>$\mathbf{BF_{10} = 1043.97}$ | $\beta = -1.19 \pm 0.23$<br>$\mathbf{BF_{10} = 625.22}$ | $\beta = -1.29 \pm 0.27$<br>$\mathbf{BF_{10} = 340.11}$ |
| 22 (21) | $\beta = -0.89 \pm 0.22$<br>$\mathbf{BF_{10} = 47.24}$ | $\beta = -0.14 \pm 0.16$<br>$BF_{10} = 0.33$ | $\beta = 0.00 \pm 0.10$<br>$BF_{10} = 0.23$ | $\beta = -0.75 \pm 0.21$<br>$\mathbf{BF_{10} = 18.80}$ | $\beta = -0.65 \pm 0.23$<br>$\mathbf{BF_{10} = 5.02}$ | $\beta = -0.89 \pm 0.25$<br>$\mathbf{BF_{10} = 18.58}$ |

**ME RIGHT**

| Channel (Channel Frequency after Pruning) | **NO SAC** | CAR | **GCR** | **SSR** | **GLM ALL** | **GLM BH** |
|---|---|---|---|---|---|---|
| 1 (23) | $\beta = -0.03 \pm 0.08$<br>$BF_{10} = 0.24$ | $\beta = 0.82 \pm 0.17$<br>$\mathbf{BF_{10} = 209.78}$ | $\beta = 0.57 \pm 0.12$<br>$\mathbf{BF_{10} = 386.50}$ | $\beta = 0.02 \pm 0.08$<br>$BF_{10} = 0.23$ | $\beta = 0.03 \pm 0.06$<br>$BF_{10} = 0.24$ | $\beta = -0.01 \pm 0.07$<br>$BF_{10} = 0.22$ |
| 2 (16) | $\beta = -0.46 \pm 0.14$<br>$\mathbf{BF_{10} = 11.51}$ | $\beta = 0.22 \pm 0.13$<br>$BF_{10} = 0.81$ | $\beta = 0.16 \pm 0.11$<br>$BF_{10} = 0.58$ | $\beta = -0.37 \pm 0.13$<br>$\mathbf{BF_{10} = 3.80}$ | $\beta = -0.42 \pm 0.17$<br>$BF_{10} = 2.86$ | $\beta = -0.46 \pm 0.14$<br>$\mathbf{BF_{10} = 10.08}$ |
| 4 (17) | $\beta = -0.13 \pm 0.08$<br>$BF_{10} = 0.77$ | $\beta = 0.49 \pm 0.11$<br>$\mathbf{BF_{10} = 66.56}$ | $\beta = 0.33 \pm 0.09$<br>$\mathbf{BF_{10} = 19.30}$ | $\beta = -0.06 \pm 0.07$<br>$BF_{10} = 0.32$ | $\beta = -0.08 \pm 0.06$<br>$BF_{10} = 0.56$ | $\beta = -0.15 \pm 0.09$<br>$BF_{10} = 0.77$ |
| 6 (24) | $\beta = -0.83 \pm 0.21$<br>$\mathbf{BF_{10} = 48.42}$ | $\beta = -0.11 \pm 0.12$<br>$BF_{10} = 0.31$ | $\beta = 0.16 \pm 0.12$<br>$BF_{10} = 0.44$ | $\beta = -0.83 \pm 0.21$<br>$\mathbf{BF_{10} = 41.68}$ | $\beta = -0.62 \pm 0.16$<br>$\mathbf{BF_{10} = 49.24}$ | $\beta = -0.70 \pm 0.18$<br>$\mathbf{BF_{10} = 54.63}$ |
| 7 (19) | $\beta = -1.15 \pm 0.21$<br>$\mathbf{BF_{10} = 695.14}$ | $\beta = -0.40 \pm 0.15$<br>$\mathbf{BF_{10} = 3.93}$ | $\beta = -0.14 \pm 0.15$<br>$BF_{10} = 0.35$ | $\beta = -1.07 \pm 0.22$<br>$\mathbf{BF_{10} = 286.24}$ | $\beta = -0.89 \pm 0.19$<br>$\mathbf{BF_{10} = 196.74}$ | $\beta = -0.99 \pm 0.21$<br>$\mathbf{BF_{10} = 157.14}$ |
| 8 (23) | $\beta = -1.23 \pm 0.22$<br>$\mathbf{BF_{10} = 1.70 \cdot 10^3}$ | $\beta = -0.45 \pm 0.15$<br>$\mathbf{BF_{10} = 8.15}$ | $\beta = -0.19 \pm 0.13$<br>$BF_{10} = 0.59$ | $\beta = -1.15 \pm 0.20$<br>$\mathbf{BF_{10} = 2.10 \cdot 10^3}$ | $\beta = -0.97 \pm 0.22$<br>$\mathbf{BF_{10} = 133.05}$ | $\beta = -1.05 \pm 0.21$<br>$\mathbf{BF_{10} = 406.28}$ |
| 10 (17) | $\beta = -0.69 \pm 0.16$<br>$\mathbf{BF_{10} = 72.47}$ | $\beta = 0.04 \pm 0.10$<br>$BF_{10} = 0.26$ | $\beta = -0.13 \pm 0.10$<br>$BF_{10} = 0.51$ | $\beta = -0.53 \pm 0.15$<br>$\mathbf{BF_{10} = 18.71}$ | $\beta = -0.44 \pm 0.23$<br>$BF_{10} = 1.09$ | $\beta = -0.44 \pm 0.14$<br>$\mathbf{BF_{10} = 7.38}$ |
| 11 (20) | $\beta = -0.92 \pm 0.21$<br>$\mathbf{BF_{10} = 88.97}$ | $\beta = -0.13 \pm 0.11$<br>$BF_{10} = 0.42$ | $\beta = -0.33 \pm 0.12$<br>$\mathbf{BF_{10} = 4.35}$ | $\beta = -0.76 \pm 0.22$<br>$\mathbf{BF_{10} = 15.01}$ | $\beta = -0.56 \pm 0.21$<br>$\mathbf{BF_{10} = 3.89}$ | $\beta = -0.64 \pm 0.21$<br>$\mathbf{BF_{10} = 6.50}$ |
| 12 (19) | $\beta = -0.88 \pm 0.17$<br>$\mathbf{BF_{10} = 572.69}$ | $\beta = -0.23 \pm 0.12$<br>$BF_{10} = 1.07$ | $\beta = -0.27 \pm 0.11$<br>$BF_{10} = 2.53$ | $\beta = -0.71 \pm 0.18$<br>$\mathbf{BF_{10} = 33.83}$ | $\beta = -0.54 \pm 0.14$<br>$\mathbf{BF_{10} = 40.72}$ | $\beta = -0.67 \pm 0.18$<br>$\mathbf{BF_{10} = 30.09}$ |
| 13 (17) | $\beta = -0.16 \pm 0.11$<br>$BF_{10} = 0.59$ | $\beta = 0.40 \pm 0.09$<br>$\mathbf{BF_{10} = 53.05}$ | $\beta = 0.25 \pm 0.09$<br>$BF_{10} = 3.64$ | $\beta = -0.14 \pm 0.08$<br>$BF_{10} = 0.88$ | $\beta = -0.20 \pm 0.21$<br>$BF_{10} = 0.38$ | $\beta = -0.11 \pm 0.13$<br>$BF_{10} = 0.35$ |
| 15 (23) | $\beta = -0.69 \pm 0.17$<br>$\mathbf{BF_{10} = 71.46}$ | $\beta = 0.07 \pm 0.15$<br>$BF_{10} = 0.42$ | $\beta = -0.11 \pm 0.11$<br>$BF_{10} = 0.34$ | $\beta = -0.71 \pm 0.17$<br>$\mathbf{BF_{10} = 90.63}$ | $\beta = -0.51 \pm 0.13$<br>$\mathbf{BF_{10} = 52.21}$ | $\beta = -0.64 \pm 0.16$<br>$\mathbf{BF_{10} = 44.04}$ |
| 17 (20) | $\beta = -1.90 \pm 0.67$<br>$\mathbf{BF_{10} = 4.85}$ | $\beta = -0.95 \pm 0.50$<br>$BF_{10} = 1.05$ | $\beta = -0.54 \pm 0.37$<br>$BF_{10} = 0.58$ | $\beta = -1.73 \pm 0.69$<br>$BF_{10} = 2.72$ | $\beta = -1.22 \pm 0.51$<br>$\mathbf{BF_{10} = 2.27}$ | $\beta = -1.49 \pm 0.66$<br>$BF_{10} = 1.74$ |
| 18 (21) | $\beta = -1.19 \pm 0.20$<br>$\mathbf{BF_{10} = 30.03 \cdot 10^3}$ | $\beta = -0.29 \pm 0.12$<br>$BF_{10} = 2.46$ | $\beta = 0.08 \pm 0.19$<br>$BF_{10} = 0.25$ | $\beta = -1.03 \pm 0.20$<br>$\mathbf{BF_{10} = 602.32}$ | $\beta = -0.71 \pm 0.16$<br>$\mathbf{BF_{10} = 101.70}$ | $\beta = -0.95 \pm 0.20$<br>$\mathbf{BF_{10} = 264.48}$ |
| 19 (20) | $\beta = -0.24 \pm 0.15$<br>$BF_{10} = 0.67$ | $\beta = 0.39 \pm 0.13$<br>$\mathbf{BF_{10} = 7.54}$ | $\beta = 0.35 \pm 0.10$<br>$\mathbf{BF_{10} = 18.18}$ | $\beta = -0.15 \pm 0.08$<br>$BF_{10} = 0.84$ | $\beta = -0.10 \pm 0.05$<br>$BF_{10} = 1.10$ | $\beta = -0.13 \pm 0.10$<br>$BF_{10} = 0.53$ |
| 21 (23) | $\beta = -0.75 \pm 0.16$<br>$\mathbf{BF_{10} = 194.00}$ | $\beta = 0.04 \pm 0.12$<br>$BF_{10} = 0.23$ | $\beta = -0.11 \pm 0.10$<br>$BF_{10} = 0.39$ | $\beta = -0.59 \pm 0.15$<br>$\mathbf{BF_{10} = 53.12}$ | $\beta = -0.46 \pm 0.12$<br>$\mathbf{BF_{10} = 47.02}$ | $\beta = -0.57 \pm 0.16$<br>$\mathbf{BF_{10} = 20.48}$ |
| 22 (21) | $\beta = -0.41 \pm 0.12$<br>$\mathbf{BF_{10} = 16.41}$ | $\beta = 0.27 \pm 0.08$<br>$\mathbf{BF_{10} = 11.84}$ | $\beta = 0.14 \pm 0.07$<br>$BF_{10} = 1.20$ | $\beta = -0.21 \pm 0.12$<br>$BF_{10} = 0.87$ | $\beta = -0.18 \pm 0.08$<br>$BF_{10} = 1.71$ | $\beta = -0.33 \pm 0.11$<br>$\mathbf{BF_{10} = 6.66}$ |

**Table 25** Results of the Bayesian t-tests of the BETA MAPS analysis of the MI $\Delta[HbO]$ data. Mean $\pm$ SEM represent mean beta values of the respective channel and its standard error of the mean across participants, $BF_{10}$ represent Bayes factors. Bayesian t-tests were performed with the *BayesFactor* package in R for which the default prior odds is set at $P(M) = \frac{\sqrt{2}}{2}$. Corresponding BETA MAPS are visualized in Figure 10 B of the main document.

| | MI | | | | | |
|---|---|---|---|---|---|---|
| Channel (Channel Frequency after Pruning) | **NO SAC** | **CAR** | **GCR** | **SSR** | **GLM ALL** | **GLM BH** |
| 1 (23) | $\beta = 1.33 \pm 0.39$ | $\beta = -0.72 \pm 0.23$ | $\beta = -0.55 \pm 0.19$ | $\beta = 0.39 \pm 0.14$ | $\beta = 0.06 \pm 0.08$ | $\beta = 0.12 \pm 0.16$ |
| | $\mathbf{BF_{10} = 15.67}$ | $\mathbf{BF_{10} = 8.88}$ | $\mathbf{BF_{10} = 5.73}$ | $\mathbf{BF_{10} = 4.42}$ | $BF_{10} = 0.27$ | $BF_{10} = 0.28$ |
| 2 (16) | $\beta = 1.69 \pm 0.47$ | $\beta = 0.06 \pm 0.27$ | $\beta = 0.00 \pm 0.21$ | $\beta = 0.82 \pm 0.33$ | $\beta = 0.55 \pm 0.30$ | $\beta = 0.61 \pm 0.29$ |
| | $\mathbf{BF_{10} = 16.02}$ | $BF_{10} = 0.26$ | $BF_{10} = 0.25$ | $BF_{10} = 2.48$ | $BF_{10} = 1.01$ | $BF_{10} = 1.45$ |
| 4 (17) | $\beta = 1.87 \pm 0.42$ | $\beta = -0.37 \pm 0.18$ | $\beta = -0.25 \pm 0.13$ | $\beta = 0.44 \pm 0.15$ | $\beta = 0.27 \pm 0.12$ | $\beta = 0.35 \pm 0.11$ |
| | $\mathbf{BF_{10} = 79.26}$ | $BF_{10} = 1.37$ | $BF_{10} = 1.01$ | $\mathbf{BF_{10} = 4.62}$ | $BF_{10} = 1.81$ | $BF_{10} = 10.06$ |
| 6 (24) | $\beta = 2.24 \pm 0.38$ | $\beta = 0.11 \pm 0.15$ | $\beta = -0.08 \pm 0.12$ | $\beta = 1.19 \pm 0.23$ | $\beta = 0.77 \pm 0.20$ | $\beta = 0.99 \pm 0.24$ |
| | $\mathbf{BF_{10} = 4086.44}$ | $BF_{10} = 0.28$ | $BF_{10} = 0.27$ | $\mathbf{BF_{10} = 644.22}$ | $\mathbf{BF_{10} = 43.83}$ | $\mathbf{BF_{10} = 88.39}$ |
| 7 (19) | $\beta = 2.80 \pm 0.50$ | $\beta = 0.68 \pm 0.18$ | $\beta = 0.45 \pm 0.16$ | $\beta = 1.56 \pm 0.27$ | $\beta = 1.18 \pm 0.27$ | $\beta = 1.32 \pm 0.31$ |
| | $\mathbf{BF_{10} = 998.17}$ | $\mathbf{BF_{10} = 29.27}$ | $\mathbf{BF_{10} = 5.01}$ | $\mathbf{BF_{10} = 1443.28}$ | $\mathbf{BF_{10} = 86.42}$ | $\mathbf{BF_{10} = 68.68}$ |
| 8 (23) | $\beta = 2.35 \pm 0.44$ | $\beta = 0.09 \pm 0.19$ | $\beta = -0.09 \pm 0.16$ | $\beta = 0.94 \pm 0.23$ | $\beta = 0.58 \pm 0.24$ | $\beta = 0.72 \pm 0.22$ |
| | $\mathbf{BF_{10} = 1084.66}$ | $BF_{10} = 0.24$ | $BF_{10} = 0.25$ | $\mathbf{BF_{10} = 70.38}$ | $BF_{10} = 2.53$ | $\mathbf{BF_{10} = 11.79}$ |
| 10 (17) | $\beta = 2.26 \pm 0.46$ | $\beta = 0.33 \pm 0.30$ | $\beta = 0.35 \pm 0.26$ | $\beta = 0.92 \pm 0.25$ | $\beta = 0.66 \pm 0.21$ | $\beta = 0.82 \pm 0.28$ |
| | $\mathbf{BF_{10} = 189.59}$ | $BF_{10} = 0.41$ | $BF_{10} = 0.54$ | $\mathbf{BF_{10} = 21.59}$ | $\mathbf{BF_{10} = 7.21}$ | $\mathbf{BF_{10} = 5.47}$ |
| 11 (20) | $\beta = 2.00 \pm 0.43$ | $\beta = 0.00 \pm 0.11$ | $\beta = 0.08 \pm 0.15$ | $\beta = 0.60 \pm 0.21$ | $\beta = 0.43 \pm 0.15$ | $\beta = 0.45 \pm 0.17$ |
| | $\mathbf{BF_{10} = 153.19}$ | $BF_{10} = 0.22$ | $BF_{10} = 0.27$ | $\mathbf{BF_{10} = 4.59}$ | $\mathbf{BF_{10} = 4.46}$ | $\mathbf{BF_{10} = 3.56}$ |
| 12 (19) | $\beta = 2.17 \pm 0.43$ | $\beta = 0.35 \pm 0.17$ | $\beta = 0.33 \pm 0.14$ | $\beta = 0.88 \pm 0.26$ | $\beta = 0.55 \pm 0.15$ | $\beta = 0.73 \pm 0.20$ |
| | $\mathbf{BF_{10} = 301.58}$ | $BF_{10} = 1.31$ | $BF_{10} = 2.38$ | $\mathbf{BF_{10} = 12.73}$ | $\mathbf{BF_{10} = 21.22}$ | $\mathbf{BF_{10} = 24.02}$ |
| 13 (17) | $\beta = 2.03 \pm 0.42$ | $\beta = -0.13 \pm 0.18$ | $\beta = -0.01 \pm 0.20$ | $\beta = 0.59 \pm 0.23$ | $\beta = 0.38 \pm 0.14$ | $\beta = 0.52 \pm 0.15$ |
| | $\mathbf{BF_{10} = 156.26}$ | $BF_{10} = 0.32$ | $BF_{10} = 0.25$ | $BF_{10} = 2.78$ | $\mathbf{BF_{10} = 4.04}$ | $\mathbf{BF_{10} = 12.75}$ |
| 15 (23) | $\beta = 2.22 \pm 0.41$ | $\beta = 0.07 \pm 0.21$ | $\beta = 0.19 \pm 0.17$ | $\beta = 1.34 \pm 0.29$ | $\beta = 0.71 \pm 0.23$ | $\beta = 1.16 \pm 0.31$ |
| | $\mathbf{BF_{10} = 1201.92}$ | $BF_{10} = 0.23$ | $BF_{10} = 0.37$ | $\mathbf{BF_{10} = 238.83}$ | $\mathbf{BF_{10} = 8.55}$ | $\mathbf{BF_{10} = 28.62}$ |
| 17 (20) | $\beta = 2.87 \pm 0.57$ | $\beta = 0.64 \pm 0.41$ | $\beta = 0.48 \pm 0.33$ | $\beta = 1.41 \pm 0.48$ | $\beta = 1.10 \pm 0.41$ | $\beta = 1.14 \pm 0.45$ |
| | $\mathbf{BF_{10} = 387.49}$ | $BF_{10} = 0.67$ | $BF_{10} = 0.58$ | $\mathbf{BF_{10} = 5.60}$ | $\mathbf{BF_{10} = 3.69}$ | $BF_{10} = 2.84$ |
| 18 (21) | $\beta = 1.81 \pm 0.34$ | $\beta = -0.25 \pm 0.18$ | $\beta = -043 \pm 0.20$ | $\beta = 0.45 \pm 0.14$ | $\beta = 0.16 \pm 0.11$ | $\beta = 0.30 \pm 0.13$ |
| | $\mathbf{BF_{10} = 838.01}$ | $BF_{10} = 0.53$ | $BF_{10} = 1.45$ | $\mathbf{BF_{10} = 12.37}$ | $BF_{10} = 0.58$ | $BF_{10} = 1.93$ |
| 19 (20) | $\beta = 1.35 \pm 0.43$ | $\beta = -0.48 \pm 0.21$ | $\beta = -0.26 \pm 0.11$ | $\beta = 0.12 \pm 0.17$ | $\beta = -0.09 \pm 0.11$ | $\beta = -0.05 \pm 0.18$ |
| | $\mathbf{BF_{10} = 8.75}$ | $BF_{10} = 1.89$ | $BF_{10} = 1.85$ | $BF_{10} = 0.29$ | $BF_{10} = 0.32$ | $BF_{10} = 0.24$ |
| 21 (23) | $\beta = 1.77 \pm 0.44$ | $\beta = -0.24 \pm 0.23$ | $\beta = -0.06 \pm 0.16$ | $\beta = 0.72 \pm 0.18$ | $\beta = 0.41 \pm 0.17$ | $\beta = 0.49 \pm 0.24$ |
| | $\mathbf{BF_{10} = 58.50}$ | $BF_{10} = 0.35$ | $BF_{10} = 0.24$ | $\mathbf{BF_{10} = 45.69}$ | $BF_{10} = 2.11$ | $BF_{10} = 1.25$ |
| 22 (21) | $\beta = 1.19 \pm 0.57$ | $\beta = -0.70 \pm 0.33$ | $\beta = -0.46 \pm 0.25$ | $\beta = 0.19 \pm 0.30$ | $\beta = -0.20 \pm 0.17$ | $\beta = -0.23 \pm 0.27$ |
| | $BF_{10} = 1.40$ | $BF_{10} = 1.39$ | $BF_{10} = 0.94$ | $BF_{10} = 0.27$ | $BF_{10} = 0.44$ | $BF_{10} = 0.32$ |

**Table 26** Results of the Bayesian t-tests of the BETA MAPS analysis of the $\Delta[HbR]$ of MI data. Mean $\pm$ SEM represent mean beta values of the respective channel and its standard error of the mean across participants, $BF_{10}$ represent Bayes factors. Bayesian t-tests were performed with the *BayesFactor* package in R for which the default prior odds is set at $P(M) = \frac{\sqrt{2}}{2}$. Corresponding BETA MAPS are visualized in Figure 10 B of the main document.

| | | | | **MI** | | |
|---|---|---|---|---|---|---|
| Channel (Channel Frequency after Pruning) | **NO SAC** | **CAR** | **GCR** | **SSR** | **GLM ALL** | **GLM BH** |
| 1 (23) | $\beta = -0.03 \pm 0.04$<br>$BF_{10} = 0.26$ | $\beta = 0.34 \pm 0.06$<br>$\mathbf{BF_{10} = 983.89}$ | $\beta = 0.30 \pm 0.06$<br>$\mathbf{BF_{10} = 989.14}$ | $\beta = 0.01 \pm 0.04$<br>$BF_{10} = 0.22$ | $\beta = 0.00 \pm 0.03$<br>$BF_{10} = 0.22$ | $\beta = -0.03 \pm 0.04$<br>$BF_{10} = 0.27$ |
| 2 (16) | $\beta = -0.19 \pm 0.08$<br>$BF_{10} = 1.86$ | $\beta = 0.11 \pm 0.08$<br>$BF_{10} = 0.62$ | $\beta = 0.10 \pm 0.06$<br>$BF_{10} = 0.75$ | $\beta = -0.13 \pm 0.09$<br>$BF_{10} = 0.64$ | $\beta = -0.16 \pm 0.09$<br>$BF_{10} = 0.86$ | $\beta = -0.17 \pm 0.08$<br>$BF_{10} = 1.39$ |
| 4 (17) | $\beta = -0.14 \pm 0.05$<br>$\mathbf{BF_{10} = 3.85}$ | $\beta = 0.15 \pm 0.05$<br>$\mathbf{BF_{10} = 12.71}$ | $\beta = 0.11 \pm 0.04$<br>$\mathbf{BF_{10} = 10.30}$ | $\beta = -0.07 \pm 0.05$<br>$BF_{10} = 0.64$ | $\beta = -0.10 \pm 0.05$<br>$BF_{10} = 1.35$ | $\beta = -0.13 \pm 0.05$<br>$BF_{10} = 2.74$ |
| 6 (24) | $\beta = -0.67 \pm 0.10$<br>$\mathbf{BF_{10} = 13.94 \cdot 10^3}$ | $\beta = -0.31 \pm 0.07$<br>$\mathbf{BF_{10} = 88.67}$ | $\beta = -0.21 \pm 0.08$<br>$\mathbf{BF_{10} = 3.91}$ | $\beta = -0.64 \pm 0.10$<br>$\mathbf{BF_{10} = 14.03 \cdot 10^3}$ | $\beta = -0.49 \pm 0.06$<br>$\mathbf{BF_{10} = 134 \cdot 10^3}$ | $\beta = -0.56 \pm 0.09$<br>$\mathbf{BF_{10} = 17.26 \cdot 10^3}$ |
| 7 (19) | $\beta = -0.63 \pm 0.09$<br>$\mathbf{BF_{10} = 12.46 \cdot 10^3}$ | $\beta = -0.36 \pm 0.07$<br>$\mathbf{BF_{10} = 206.83}$ | $\beta = -0.26 \pm 0.05$<br>$\mathbf{BF_{10} = 379.26}$ | $\beta = -0.63 \pm 0.10$<br>$\mathbf{BF_{10} = 1.98 \cdot 10^3}$ | $\beta = -0.47 \pm 0.09$<br>$\mathbf{BF_{10} = 404.74}$ | $\beta = -0.51 \pm 0.08$<br>$\mathbf{BF_{10} = 2.35 \cdot 10^3}$ |
| 8 (23) | $\beta = -0.41 \pm 0.10$<br>$\mathbf{BF_{10} = 78.33}$ | $\beta = -0.03 \pm 0.07$<br>$BF_{10} = 0.23$ | $\beta = 0.06 \pm 0.06$<br>$BF_{10} = 0.32$ | $\beta = -0.36 \pm 0.11$<br>$\mathbf{BF_{10} = 10.39}$ | $\beta = -0.29 \pm 0.11$<br>$\mathbf{BF_{10} = 3.41}$ | $\beta = -0.33 \pm 0.09$<br>$\mathbf{BF_{10} = 36.12}$ |
| 10 (17) | $\beta = -0.55 \pm 0.16$<br>$\mathbf{BF_{10} = 15.51}$ | $\beta = -0.18 \pm 0.11$<br>$BF_{10} = 0.74$ | $\beta = -0.20 \pm 0.11$<br>$BF_{10} = 1.08$ | $\beta = -0.46 \pm 0.15$<br>$\mathbf{BF_{10} = 7.41}$ | $\beta = -0.37 \pm 0.15$<br>$\mathbf{BF_{10} = 2.58}$ | $\beta = -0.40 \pm 0.13$<br>$\mathbf{BF_{10} = 7.36}$ |
| 11 (20) | $\beta = -0.35 \pm 0.05$<br>$\mathbf{BF_{10} = 61.24 \cdot 10^3}$ | $\beta = -0.08 \pm 0.03$<br>$\mathbf{BF_{10} = 4.26}$ | $\beta = -0.10 \pm 0.03$<br>$\mathbf{BF_{10} = 15.84}$ | $\beta = -0.28 \pm 0.06$<br>$\mathbf{BF_{10} = 220.29}$ | $\beta = -0.16 \pm 0.04$<br>$\mathbf{BF_{10} = 62.25}$ | $\beta = -0.22 \pm 0.03$<br>$\mathbf{BF_{10} = 8.73 \cdot 0^3}$ |
| 12 (19) | $\beta = -0.43 \pm 0.08$<br>$\mathbf{BF_{10} = 445.39}$ | $\beta = -0.14 \pm 0.06$<br>$BF_{10} = 2.19$ | $\beta = -0.13 \pm 0.04$<br>$\mathbf{BF_{10} = 4.93}$ | $\beta = -0.36 \pm 0.08$<br>$\mathbf{BF_{10} = 86.82}$ | $\beta = -0.24 \pm 0.05$<br>$\mathbf{BF_{10} = 167.63}$ | $\beta = -0.31 \pm 0.07$<br>$\mathbf{BF_{10} = 129.49}$ |
| 13 (17) | $\beta = -0.15 \pm 0.07$<br>$BF_{10} = 1.88$ | $\beta = 0.18 \pm 0.05$<br>$\mathbf{BF_{10} = 28.98}$ | $\beta = 0.11 \pm 0.04$<br>$\mathbf{BF_{10} = 5.99}$ | $\beta = -0.15 \pm 0.06$<br>$BF_{10} = 2.91$ | $\beta = -0.07 \pm 0.07$<br>$BF_{10} = 0.36$ | $\beta = -0.09 \pm 0.05$<br>$\mathbf{BF_{10} = 70.98}$ |
| 15 (23) | $\beta = -0.48 \pm 0.10$<br>$\mathbf{BF_{10} = 532.27}$ | $\beta = -0.15 \pm 0.09$<br>$BF_{10} = 0.78$ | $\beta = -0.16 \pm 0.06$<br>$BF_{10} = 2.38$ | $\beta = -0.49 \pm 0.09$<br>$\mathbf{BF_{10} = 865.75}$ | $\beta = -0.37 \pm 0.08$<br>$\mathbf{BF_{10} = 185.36}$ | $\beta = -0.44 \pm 0.09$<br>$\mathbf{BF_{10} = 357.51}$ |
| 17 (20) | $\beta = -0.47 \pm 0.12$<br>$\mathbf{BF_{10} = 31.40}$ | $\beta = -0.12 \pm 0.10$<br>$BF_{10} = 0.41$ | $\beta = -0.04 \pm 0.08$<br>$BF_{10} = 0.26$ | $\beta = -0.40 \pm 0.12$<br>$\mathbf{BF_{10} = 12.86}$ | $\beta = -0.20 \pm 0.14$<br>$BF_{10} = 0.60$ | $\beta = -0.27 \pm 0.13$<br>$BF_{10} = 1.22$ |
| 18 (21) | $\beta = -0.22 \pm 0.05$<br>$\mathbf{BF_{10} = 96.37}$ | $\beta = 0.11 \pm 0.06$<br>$BF_{10} = 1.00$ | $\beta = 0.15 \pm 0.04$<br>$\mathbf{BF_{10} = 18.69}$ | $\beta = -0.15 \pm 0.04$<br>$\mathbf{BF_{10} = 36.97}$ | $\beta = -0.02 \pm 0.04$<br>$BF_{10} = 0.25$ | $\beta = -0.09 \pm 0.04$<br>$BF_{10} = 1.38$ |
| 19 (20) | $\beta = -0.07 \pm 0.09$<br>$BF_{10} = 0.30$ | $\beta = 0.25 \pm 0.09$<br>$\mathbf{BF_{10} = 4.91}$ | $\beta = 0.16 \pm 0.06$<br>$\mathbf{BF_{10} = 4.16}$ | $\beta = 0.00 \pm 0.07$<br>$BF_{10} = 0.23$ | $\beta = 0.02 \pm 0.05$<br>$BF_{10} = 0.25$ | $\beta = -0.01 \pm 0.06$<br>$BF_{10} = 0.24$ |
| 21 (23) | $\beta = -0.33 \pm 0.08$<br>$\mathbf{BF_{10} = 116.08}$ | $\beta = 0.03 \pm 0.04$<br>$BF_{10} = 0.25$ | $\beta = 0.00 \pm 0.04$<br>$BF_{10} = 0.22$ | $\beta = -0.28 \pm 0.08$<br>$\mathbf{BF_{10} = 28.90}$ | $\beta = -0.18 \pm 0.06$<br>$\mathbf{BF_{10} = 5.67}$ | $\beta = -0.21 \pm 0.06$<br>$\mathbf{BF_{10} = 15.06}$ |
| 22 (21) | $\beta = -0.12 \pm 0.09$<br>$BF_{10} = 0.47$ | $\beta = 0.20 \pm 0.08$<br>$\mathbf{BF_{10} = 3.19}$ | $\beta = 0.13 \pm 0.06$<br>$BF_{10} = 1.40$ | $\beta = -0.03 \pm 0.08$<br>$BF_{10} = 0.24$ | $\beta = 0.01 \pm 0.06$<br>$BF_{10} = 0.23$ | $\beta = -0.04 \pm 0.08$<br>$BF_{10} = 0.25$ |

*Single Trial Time Series*

*SIM data*

The following figures visualize the normalized single-subject single-trial semi-simulated $\Delta[HbO]$ and $\Delta[HbR]$ data of each individually selected channel (based on beta values of GLM ALL corrected data) of all correction methods and all SDCs.

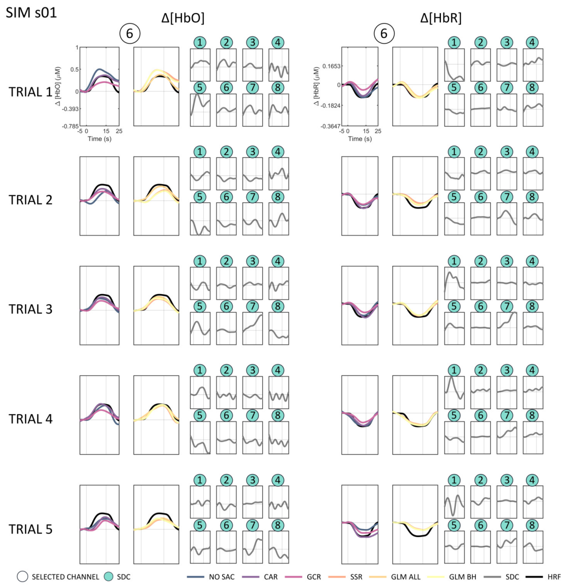


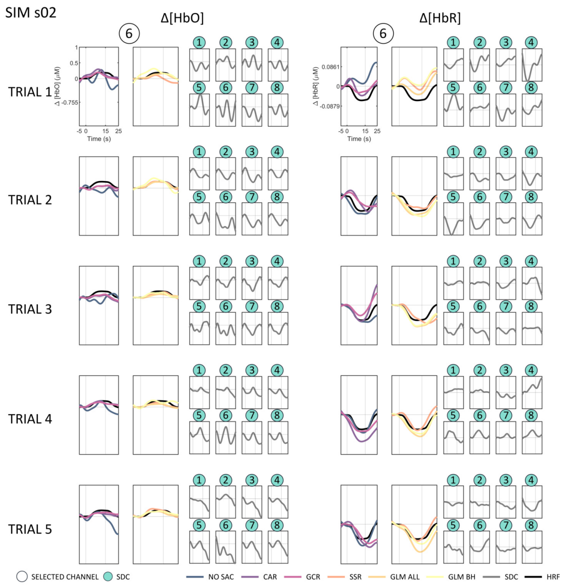


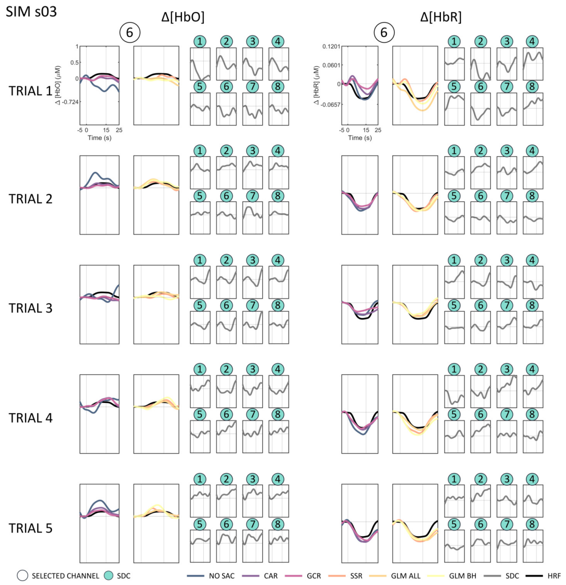


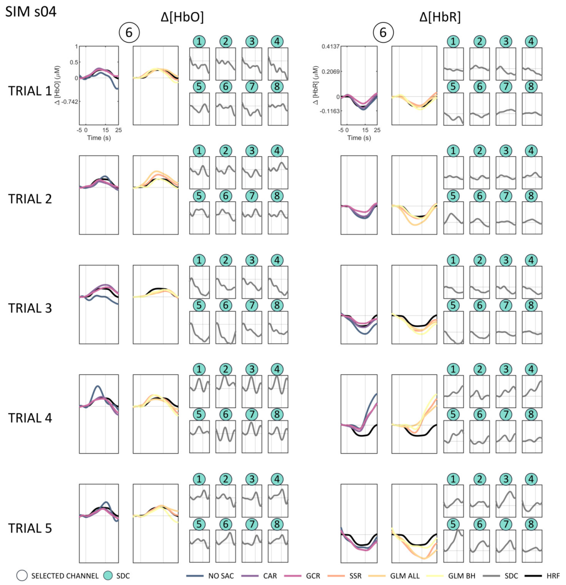

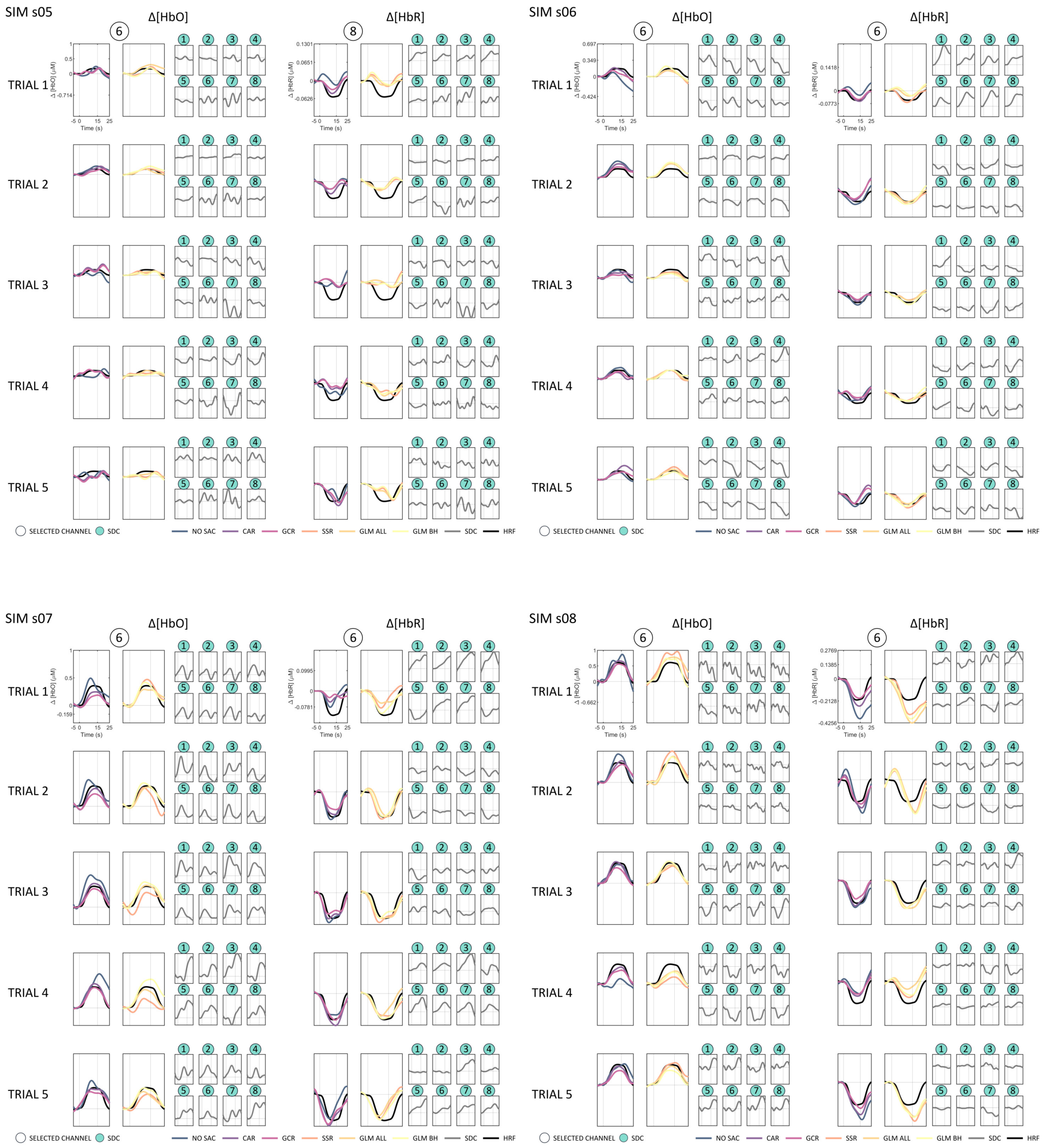
SIM s05
SIM s06
SIM s07
SIM s08
Δ[HbO]
Δ[HbR]
TRIAL 1
TRIAL 2
TRIAL 3
TRIAL 4
TRIAL 5
Time (s)
SELECTED CHANNEL
SDC
NO SAC
CAR
GCR
SSR
GLM ALL
GLM BH
SDC
HRF

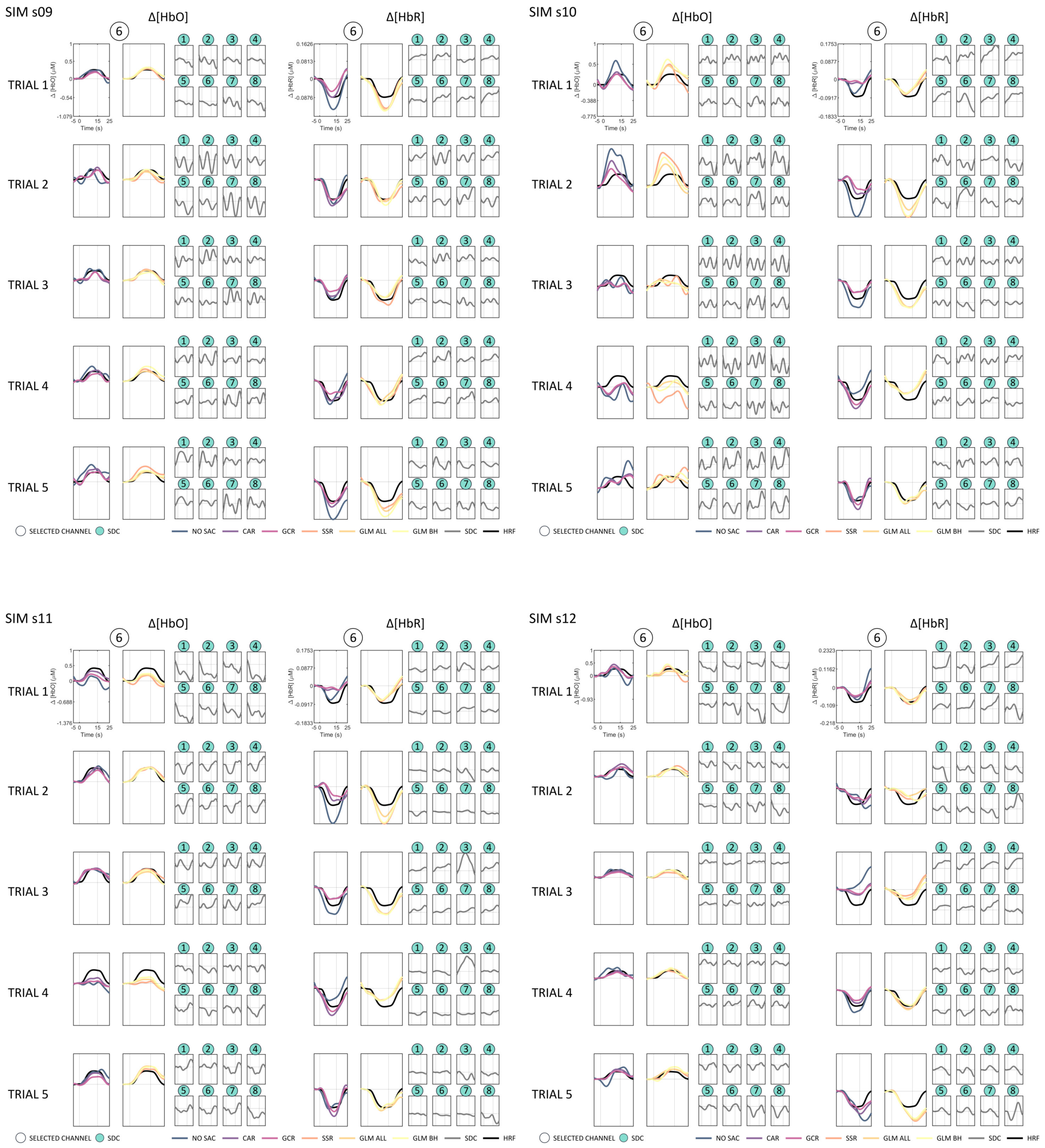
SIM s09
SIM s10
SIM s11
SIM s12
Δ[HbO]
Δ[HbR]
TRIAL 1
TRIAL 2
TRIAL 3
TRIAL 4
TRIAL 5
Time (s)
SELECTED CHANNEL
SDC
NO SAC
CAR
GCR
SSR
GLM ALL
GLM BH
SDC
HRF

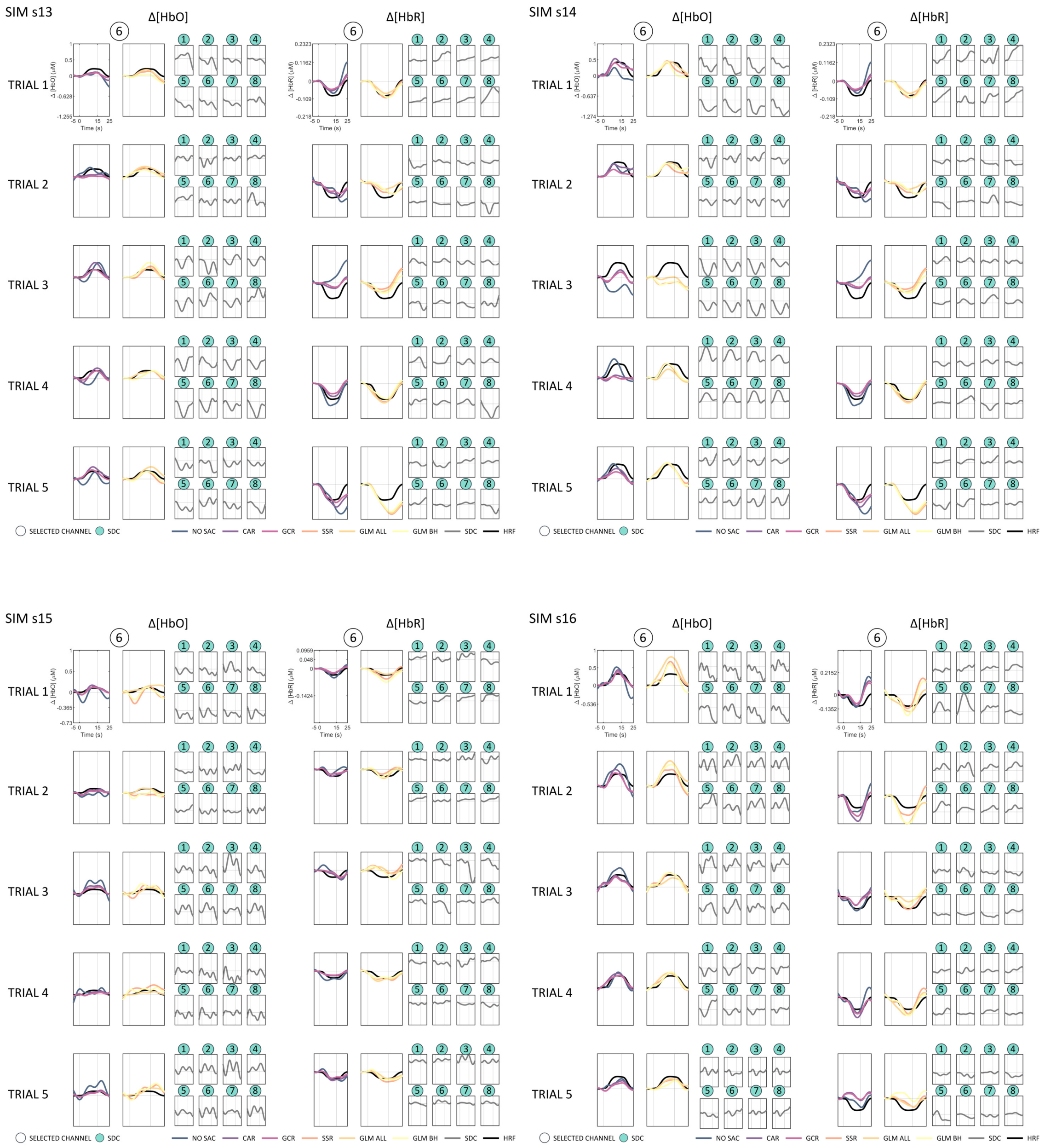
SIM s13
SIM s14
SIM s15
SIM s16
Δ[HbO]
Δ[HbR]
TRIAL 1
TRIAL 2
TRIAL 3
TRIAL 4
TRIAL 5
Time (s)
SELECTED CHANNEL
SDC
NO SAC
CAR
GCR
SSR
GLM ALL
GLM BH
SDC
HRF

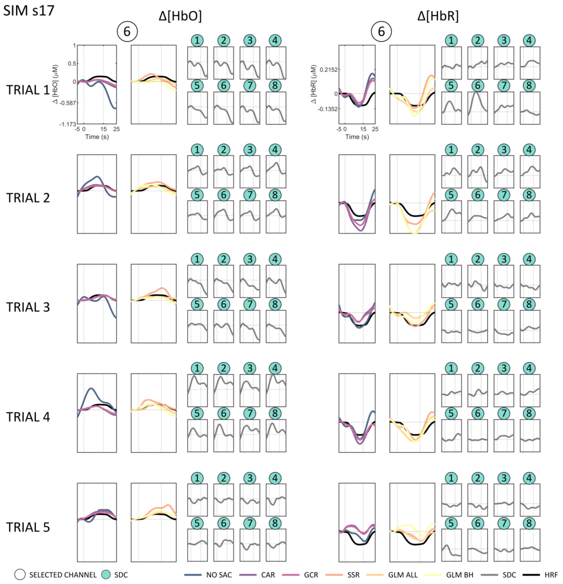
SIM s17
Δ[HbO]
Δ[HbR]
TRIAL 1
TRIAL 2
TRIAL 3
TRIAL 4
TRIAL 5
Time (s)
SELECTED CHANNEL
SDC
NO SAC
CAR
GCR
SSR
GLM ALL
GLM BH
SDC
HRF

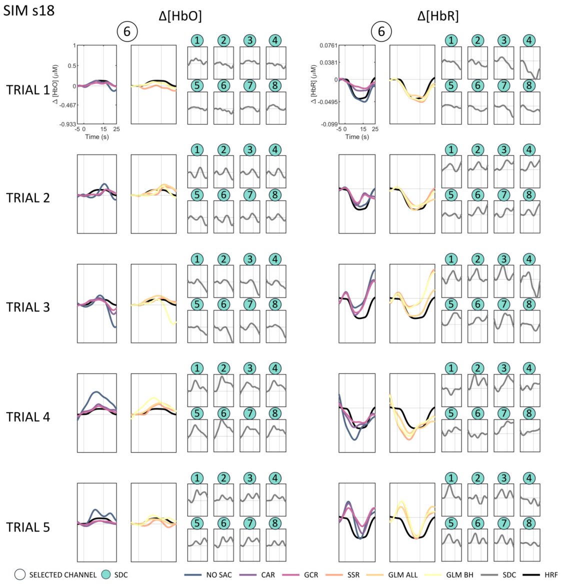
SIM s18
Δ[HbO]
Δ[HbR]
TRIAL 1
TRIAL 2
TRIAL 3
TRIAL 4
TRIAL 5
Time (s)
SELECTED CHANNEL
SDC
NO SAC
CAR
GCR
SSR
GLM ALL
GLM BH
SDC
HRF

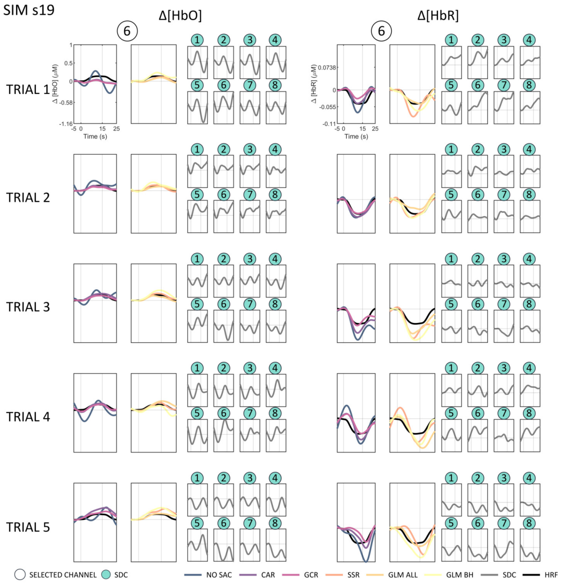
SIM s19
Δ[HbO]
Δ[HbR]
TRIAL 1
TRIAL 2
TRIAL 3
TRIAL 4
TRIAL 5
Time (s)
SELECTED CHANNEL
SDC
NO SAC
CAR
GCR
SSR
GLM ALL
GLM BH
SDC
HRF

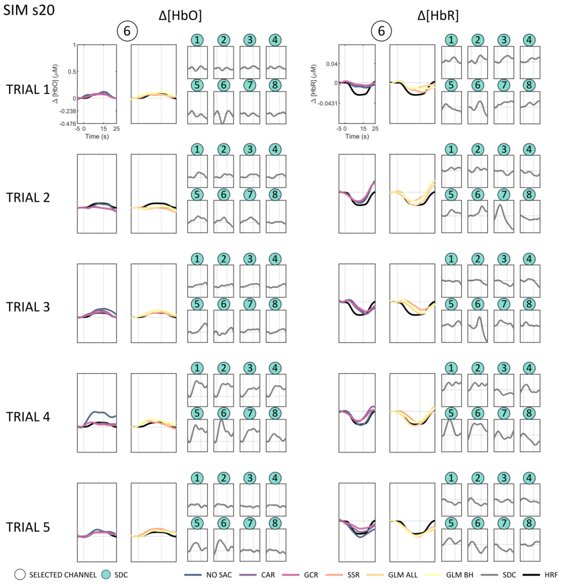
SIM s20
Δ[HbO]
Δ[HbR]
TRIAL 1
TRIAL 2
TRIAL 3
TRIAL 4
TRIAL 5
Time (s)
SELECTED CHANNEL
SDC
NO SAC
CAR
GCR
SSR
GLM ALL
GLM BH
SDC
HRF

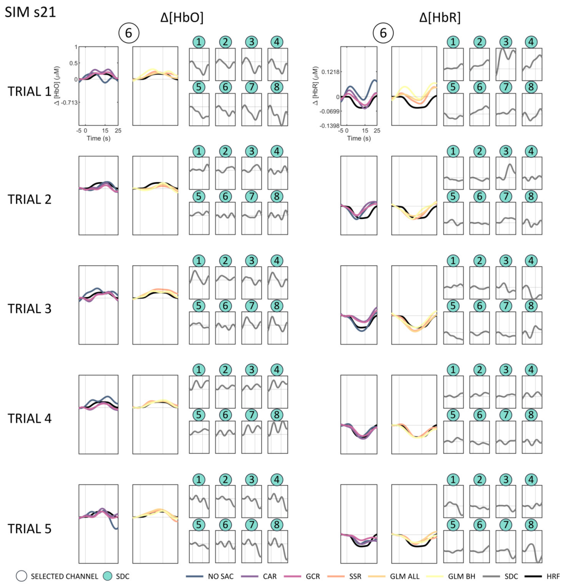
SIM s21
Δ[HbO]
Δ[HbR]
TRIAL 1
TRIAL 2
TRIAL 3
TRIAL 4
TRIAL 5
Time (s)
SELECTED CHANNEL
SDC
NO SAC
CAR
GCR
SSR
GLM ALL
GLM BH
SDC
HRF

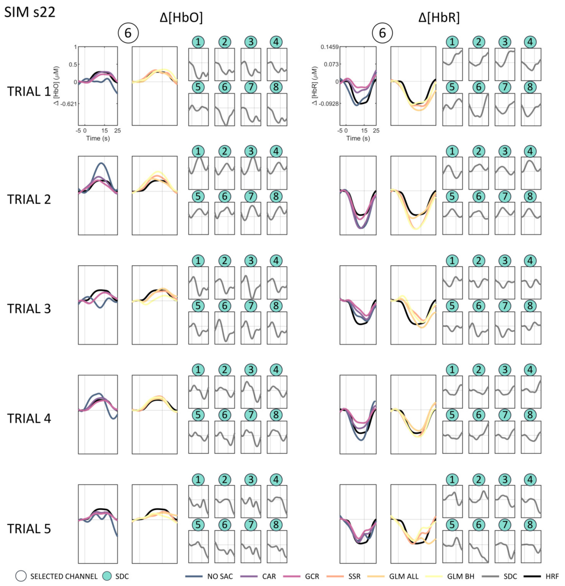
SIM s22
Δ[HbO]
Δ[HbR]
TRIAL 1
TRIAL 2
TRIAL 3
TRIAL 4
TRIAL 5
Time (s)
SELECTED CHANNEL
SDC
NO SAC
CAR
GCR
SSR
GLM ALL
GLM BH
SDC
HRF

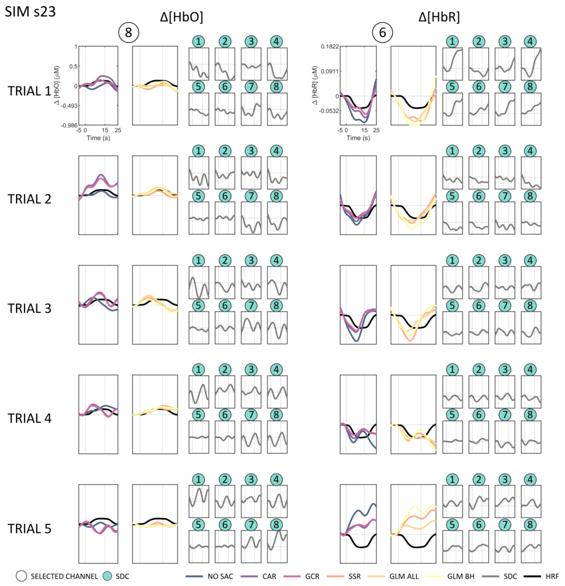
SIM s23
Δ[HbO]
Δ[HbR]
TRIAL 1
TRIAL 2
TRIAL 3
TRIAL 4
TRIAL 5
Time (s)
SELECTED CHANNEL
SDC
NO SAC
CAR
GCR
SSR
GLM ALL
GLM BH
SDC
HRF

*ME/MI data*

The following figures visualize the normalized single-subject single-trial $\Delta[HbO]$ and $\Delta[HbR]$ data resulting from the real data set of each individually selected channel (based on beta values of GLM ALL corrected data) of all correction methods and all SDCs. Note that only the first five trials are visualized (out of 12 for ME LEFT and ME RIGHT and out of 36 for MI data). As the MI data sets consists of three individual MI tasks (i.e., MI of left hand tapping, MI of right hand tapping and MI of whole body movements) the first five trials always belong to only one of the three tasks and the order of the MI tasks was pseudo-randomized between subjects.

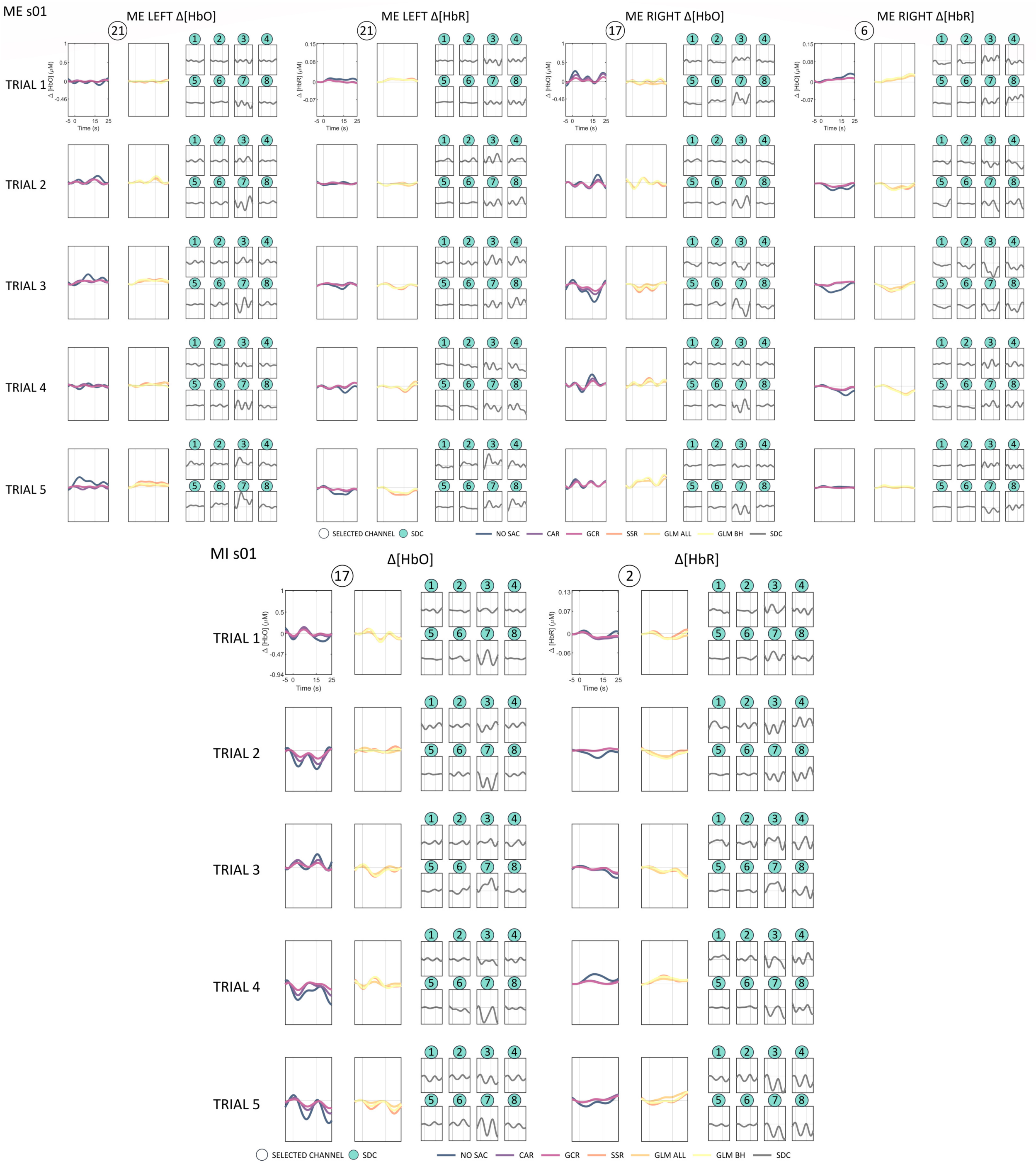
ME s01
ME LEFT Δ[HbO]
ME LEFT Δ[HbR]
ME RIGHT Δ[HbO]
ME RIGHT Δ[HbR]
TRIAL 1
TRIAL 2
TRIAL 3
TRIAL 4
TRIAL 5
SELECTED CHANNEL
SDC
NO SAC
CAR
GCR
SSR
GLM ALL
GLM BH
SDC
MI s01
Δ[HbO]
Δ[HbR]
TRIAL 1
TRIAL 2
TRIAL 3
TRIAL 4
TRIAL 5
SELECTED CHANNEL
SDC
NO SAC
CAR
GCR
SSR
GLM ALL
GLM BH
SDC

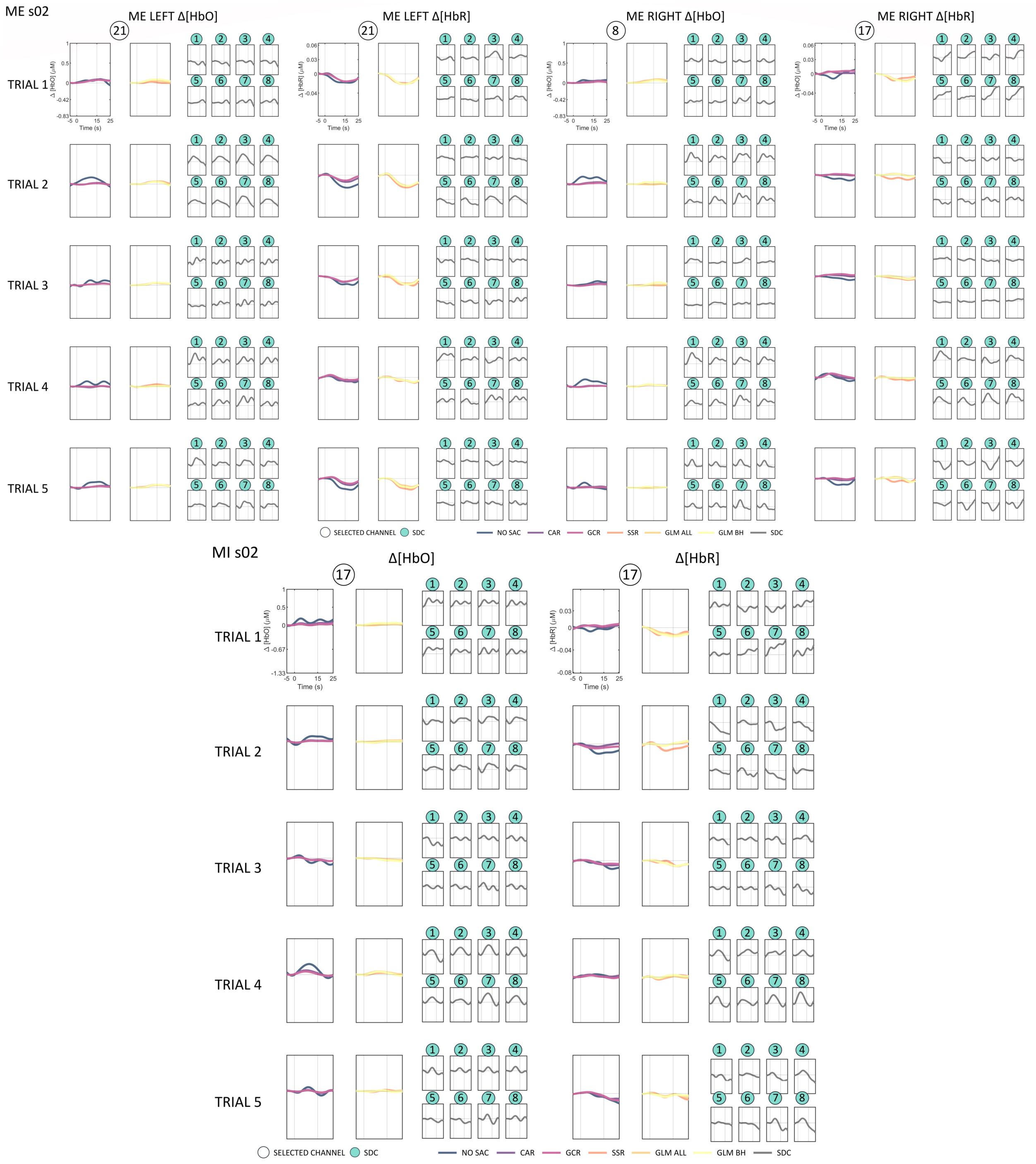

ME s02
ME LEFT Δ[HbO]
ME LEFT Δ[HbR]
ME RIGHT Δ[HbO]
ME RIGHT Δ[HbR]
TRIAL 1
TRIAL 2
TRIAL 3
TRIAL 4
TRIAL 5
Time (s)
SELECTED CHANNEL
SDC
NO SAC
CAR
GCR
SSR
GLM ALL
GLM BH
SDC
MI s02
Δ[HbO]
Δ[HbR]

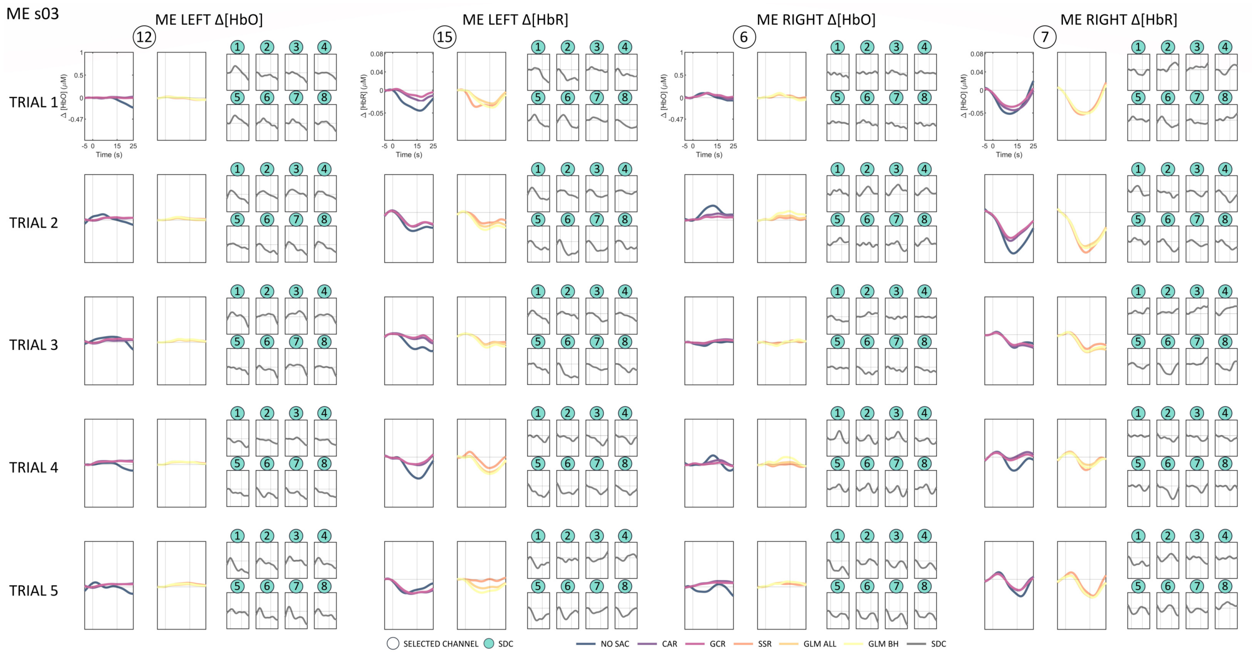
ME s03
ME LEFT Δ[HbO]
ME LEFT Δ[HbR]
ME RIGHT Δ[HbO]
ME RIGHT Δ[HbR]
TRIAL 1
TRIAL 2
TRIAL 3
TRIAL 4
TRIAL 5
Time (s)
SELECTED CHANNEL
SDC
NO SAC
CAR
GCR
SSR
GLM ALL
GLM BH
SDC

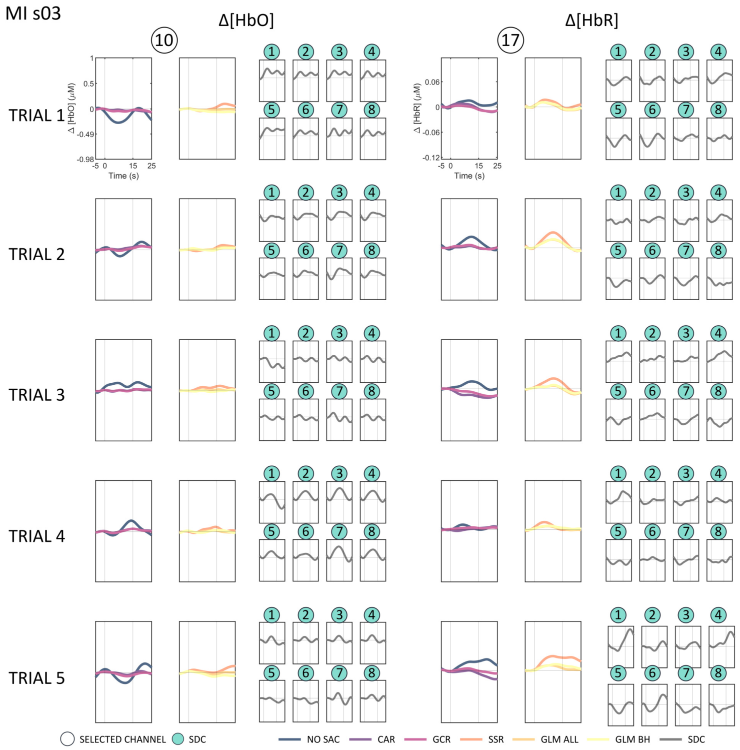
MI s03
Δ[HbO]
Δ[HbR]
TRIAL 1
TRIAL 2
TRIAL 3
TRIAL 4
TRIAL 5
Time (s)
SELECTED CHANNEL
SDC
NO SAC
CAR
GCR
SSR
GLM ALL
GLM BH
SDC

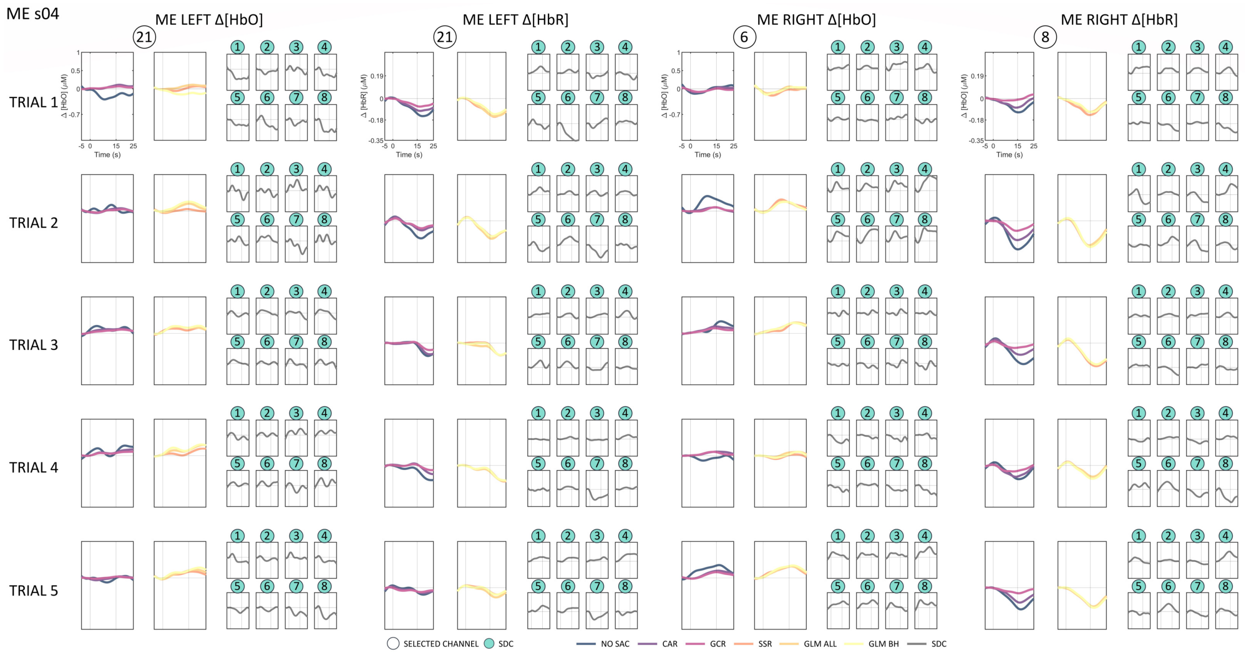
ME s04
ME LEFT Δ[HbO]
ME LEFT Δ[HbR]
ME RIGHT Δ[HbO]
ME RIGHT Δ[HbR]
TRIAL 1
TRIAL 2
TRIAL 3
TRIAL 4
TRIAL 5
Time (s)
SELECTED CHANNEL
SDC
NO SAC
CAR
GCR
SSR
GLM ALL
GLM BH
SDC

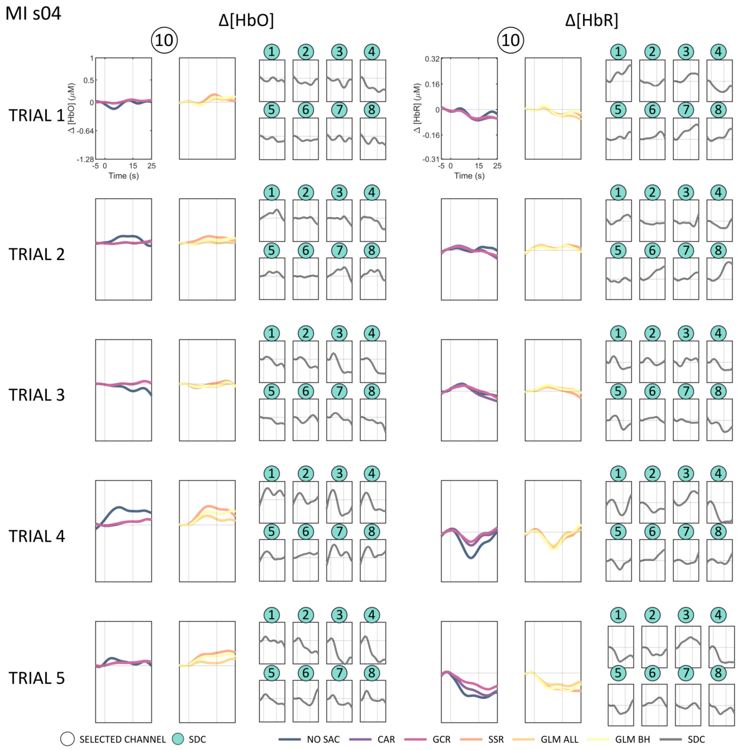
MI s04
Δ[HbO]
Δ[HbR]
TRIAL 1
TRIAL 2
TRIAL 3
TRIAL 4
TRIAL 5
Time (s)
SELECTED CHANNEL
SDC
NO SAC
CAR
GCR
SSR
GLM ALL
GLM BH
SDC

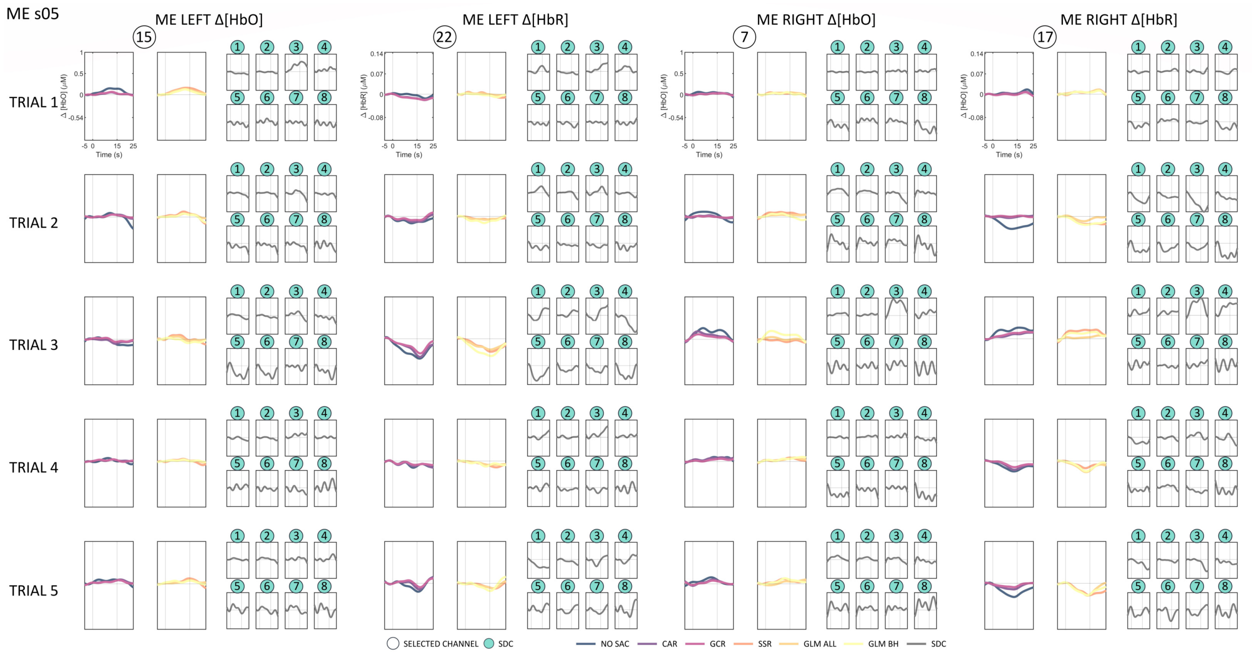
ME s05
ME LEFT Δ[HbO]
ME LEFT Δ[HbR]
ME RIGHT Δ[HbO]
ME RIGHT Δ[HbR]
TRIAL 1
TRIAL 2
TRIAL 3
TRIAL 4
TRIAL 5
SELECTED CHANNEL
SDC
NO SAC
CAR
GCR
SSR
GLM ALL
GLM BH
SDC

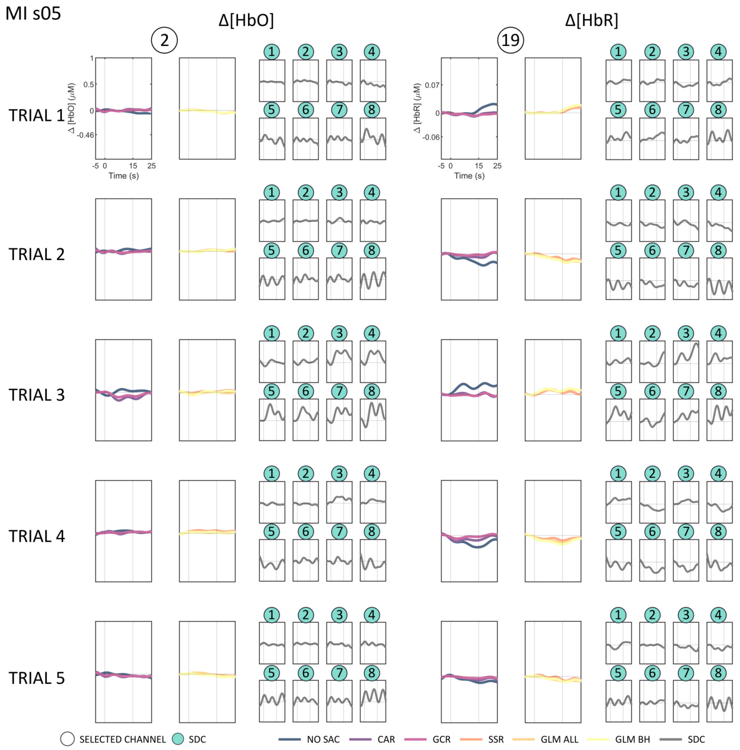
MI s05
Δ[HbO]
Δ[HbR]
TRIAL 1
TRIAL 2
TRIAL 3
TRIAL 4
TRIAL 5
SELECTED CHANNEL
SDC
NO SAC
CAR
GCR
SSR
GLM ALL
GLM BH
SDC

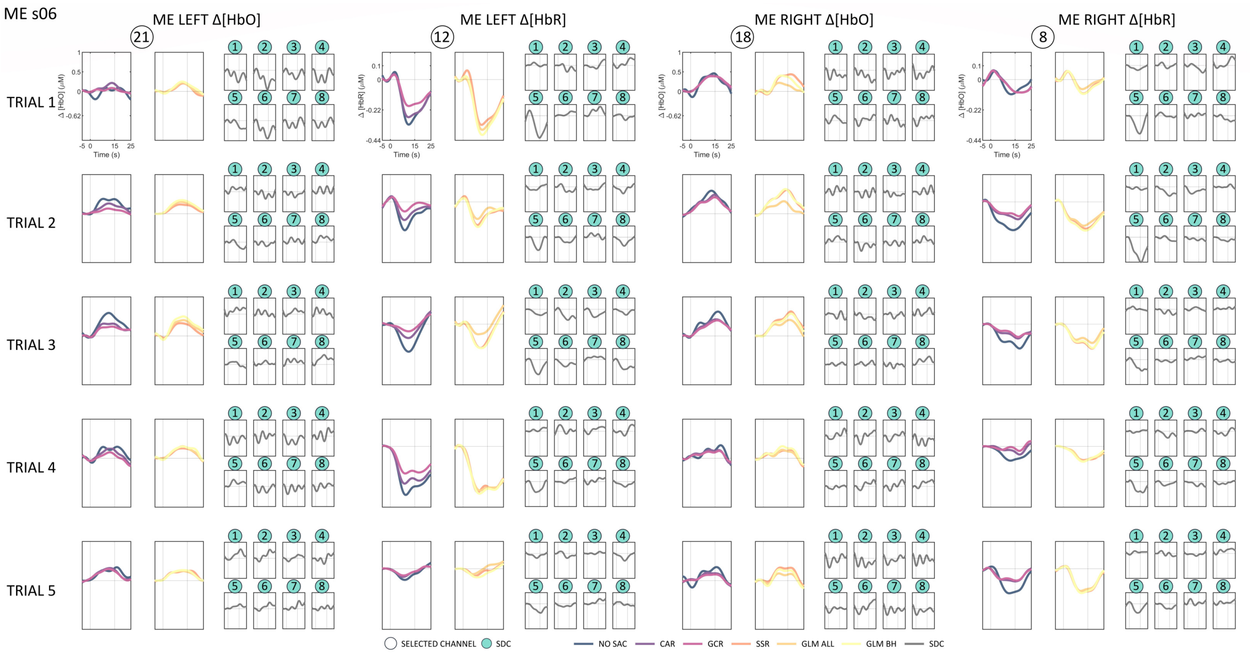
ME s06
ME LEFT Δ[HbO]
ME LEFT Δ[HbR]
ME RIGHT Δ[HbO]
ME RIGHT Δ[HbR]
TRIAL 1
TRIAL 2
TRIAL 3
TRIAL 4
TRIAL 5
Time (s)
SELECTED CHANNEL
SDC
NO SAC
CAR
GCR
SSR
GLM ALL
GLM BH
SDC

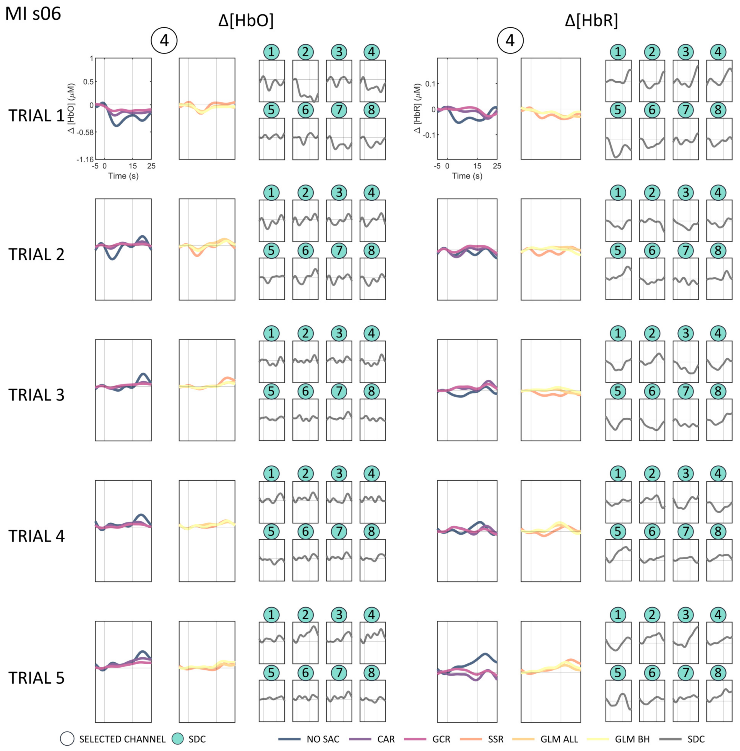
MI s06
Δ[HbO]
Δ[HbR]
TRIAL 1
TRIAL 2
TRIAL 3
TRIAL 4
TRIAL 5
Time (s)
SELECTED CHANNEL
SDC
NO SAC
CAR
GCR
SSR
GLM ALL
GLM BH
SDC

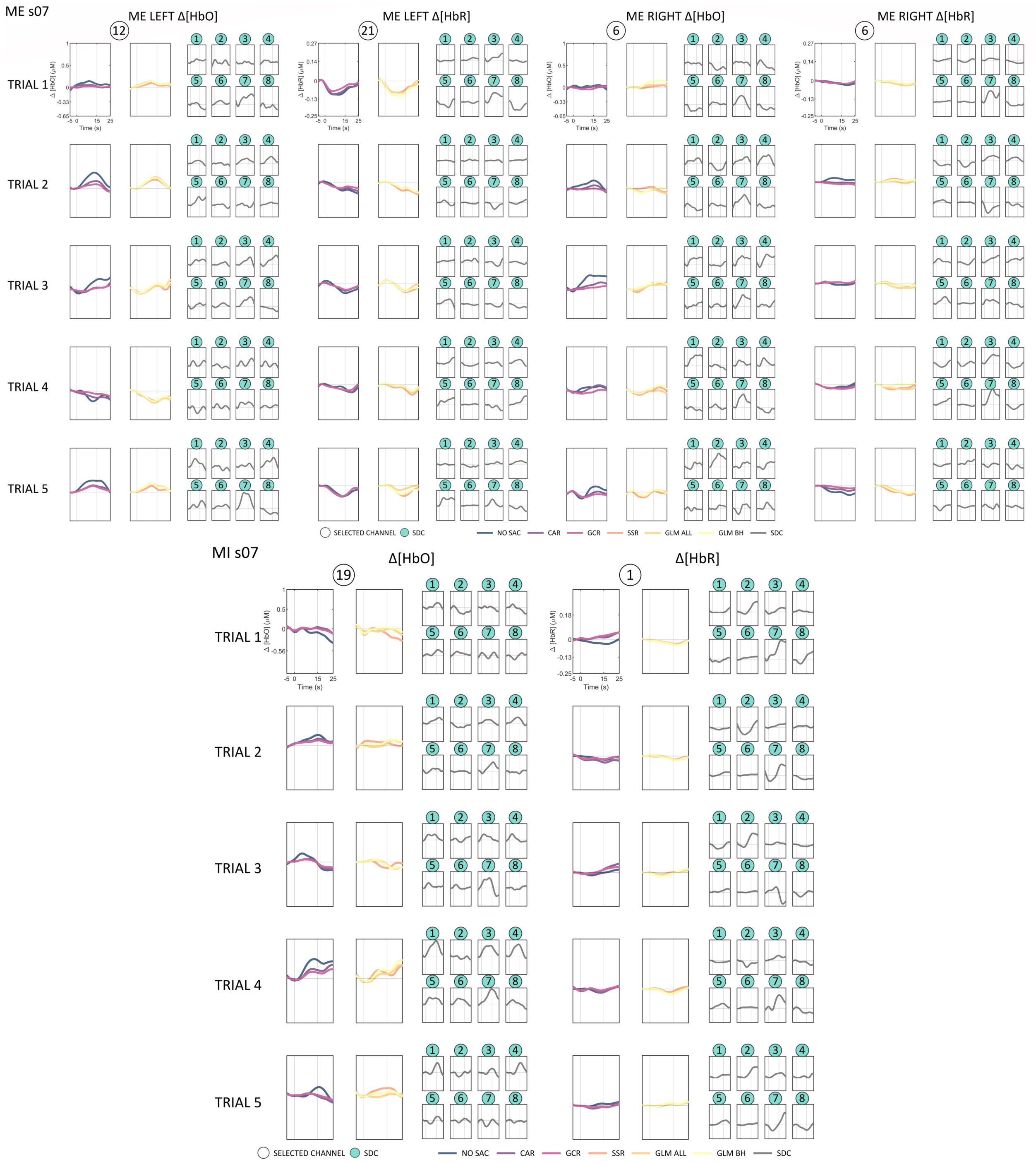
ME s07
ME LEFT Δ[HbO]
ME LEFT Δ[HbR]
ME RIGHT Δ[HbO]
ME RIGHT Δ[HbR]
TRIAL 1
TRIAL 2
TRIAL 3
TRIAL 4
TRIAL 5
SELECTED CHANNEL
SDC
NO SAC
CAR
GCR
SSR
GLM ALL
GLM BH
SDC
MI s07
Δ[HbO]
Δ[HbR]
Time (s)

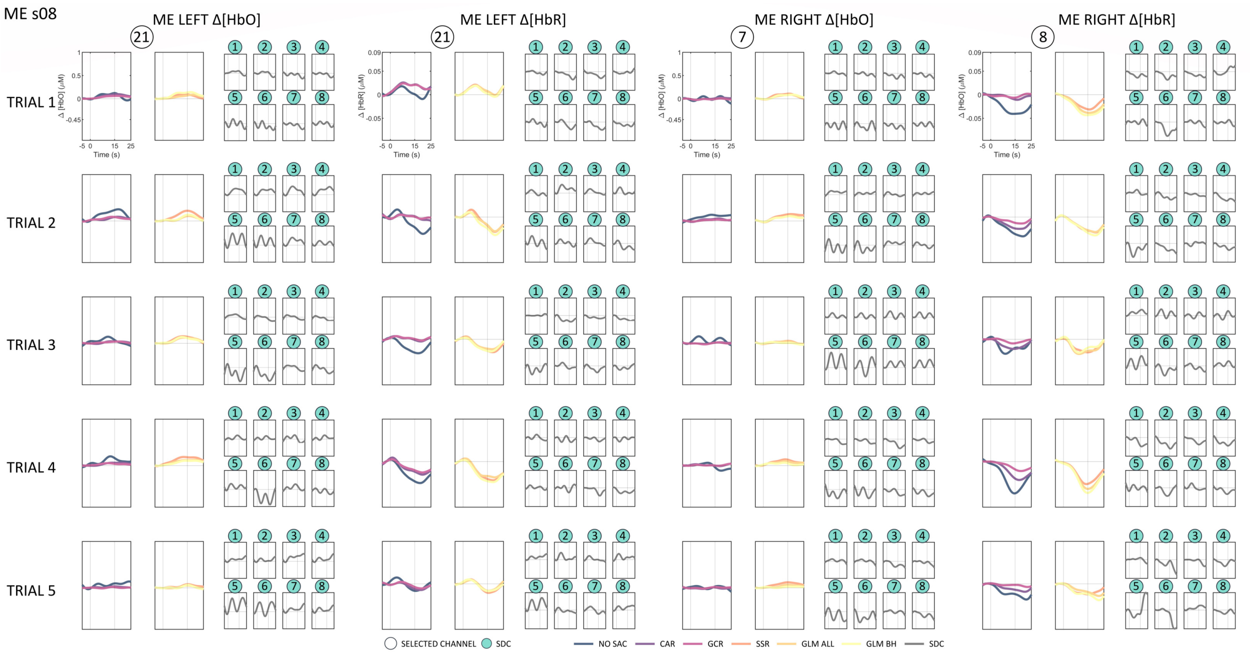
ME s08
ME LEFT Δ[HbO]
ME LEFT Δ[HbR]
ME RIGHT Δ[HbO]
ME RIGHT Δ[HbR]
TRIAL 1
TRIAL 2
TRIAL 3
TRIAL 4
TRIAL 5
Time (s)
SELECTED CHANNEL
SDC
NO SAC
CAR
GCR
SSR
GLM ALL
GLM BH
SDC

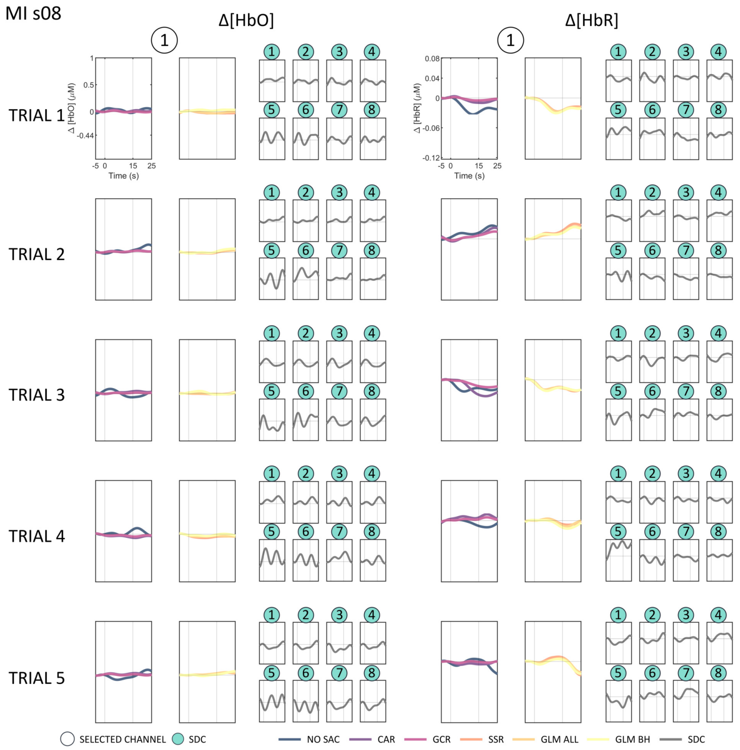
MI s08
Δ[HbO]
Δ[HbR]
TRIAL 1
TRIAL 2
TRIAL 3
TRIAL 4
TRIAL 5
Time (s)
SELECTED CHANNEL
SDC
NO SAC
CAR
GCR
SSR
GLM ALL
GLM BH
SDC

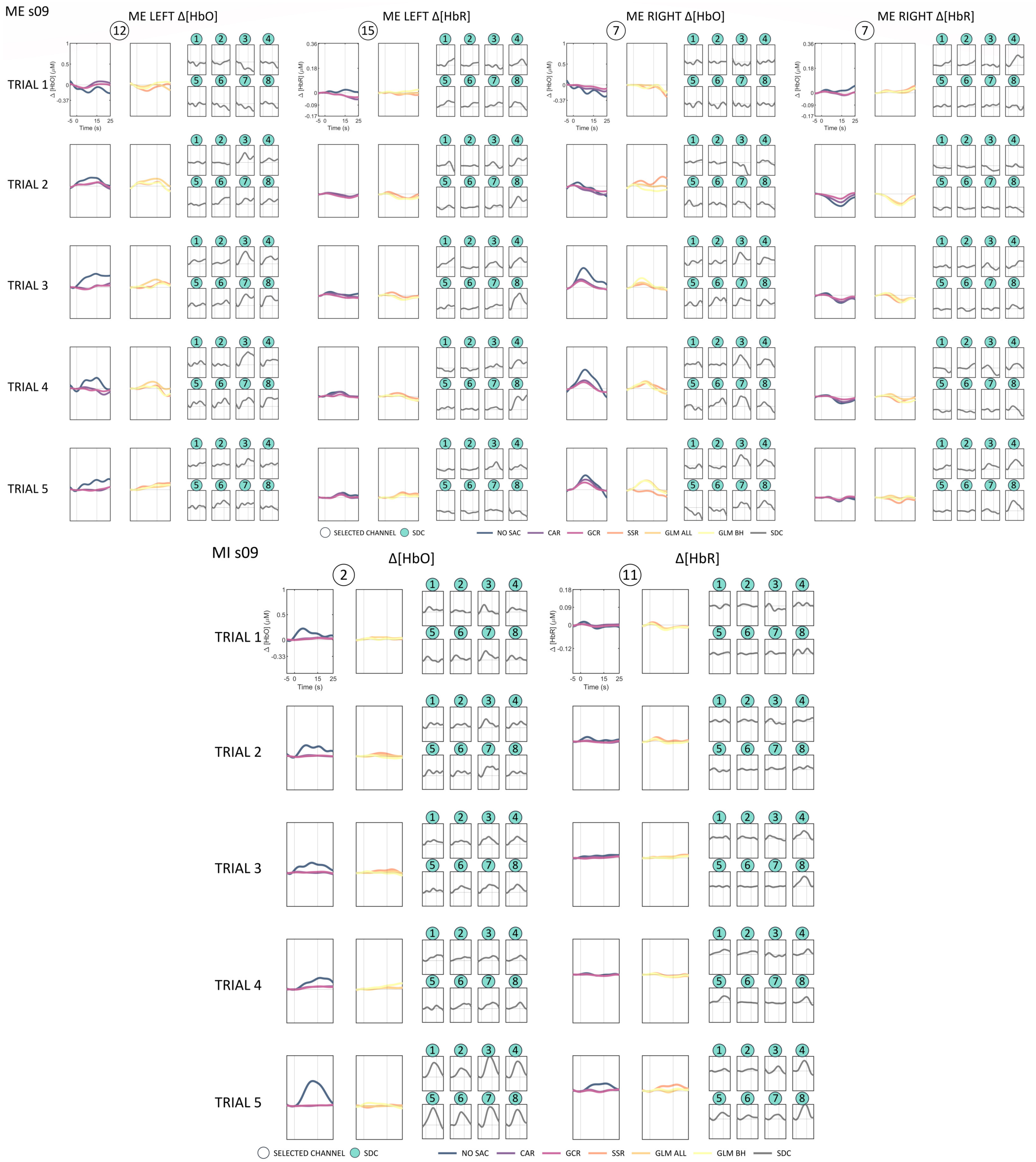
ME s09
ME LEFT Δ[HbO]
ME LEFT Δ[HbR]
ME RIGHT Δ[HbO]
ME RIGHT Δ[HbR]
TRIAL 1
TRIAL 2
TRIAL 3
TRIAL 4
TRIAL 5
Δ [HbO] (μM)
Δ [HbR] (μM)
Time (s)
SELECTED CHANNEL
SDC
NO SAC
CAR
GCR
SSR
GLM ALL
GLM BH
SDC
MI s09
Δ[HbO]
Δ[HbR]

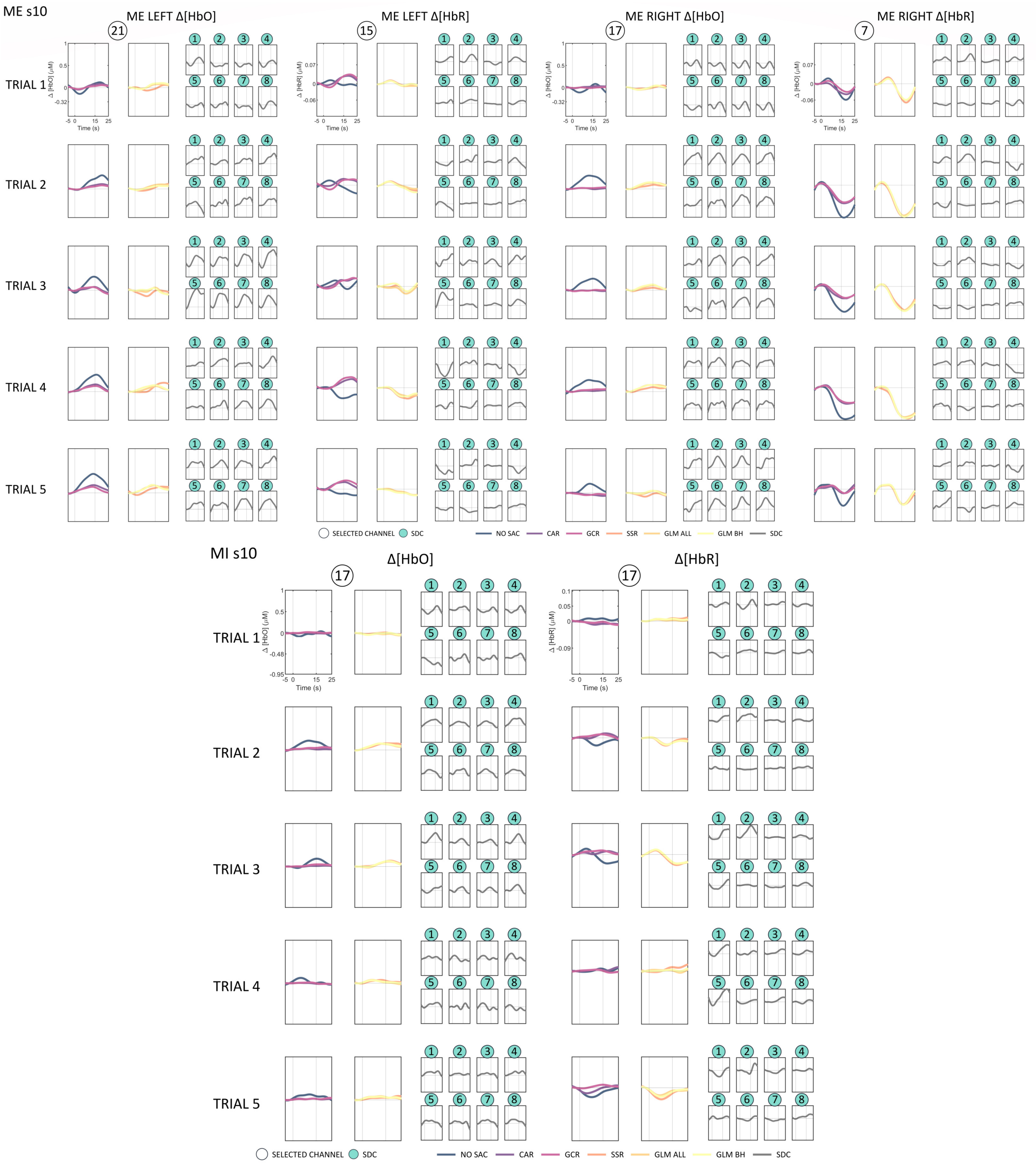
ME s10
ME LEFT Δ[HbO]
ME LEFT Δ[HbR]
ME RIGHT Δ[HbO]
ME RIGHT Δ[HbR]
21
15
17
7
TRIAL 1
TRIAL 2
TRIAL 3
TRIAL 4
TRIAL 5
Δ [HbO] (μM)
Δ [HbR] (μM)
Time (s)
SELECTED CHANNEL
SDC
NO SAC
CAR
GCR
SSR
GLM ALL
GLM BH
SDC
MI s10
Δ[HbO]
Δ[HbR]
17
17

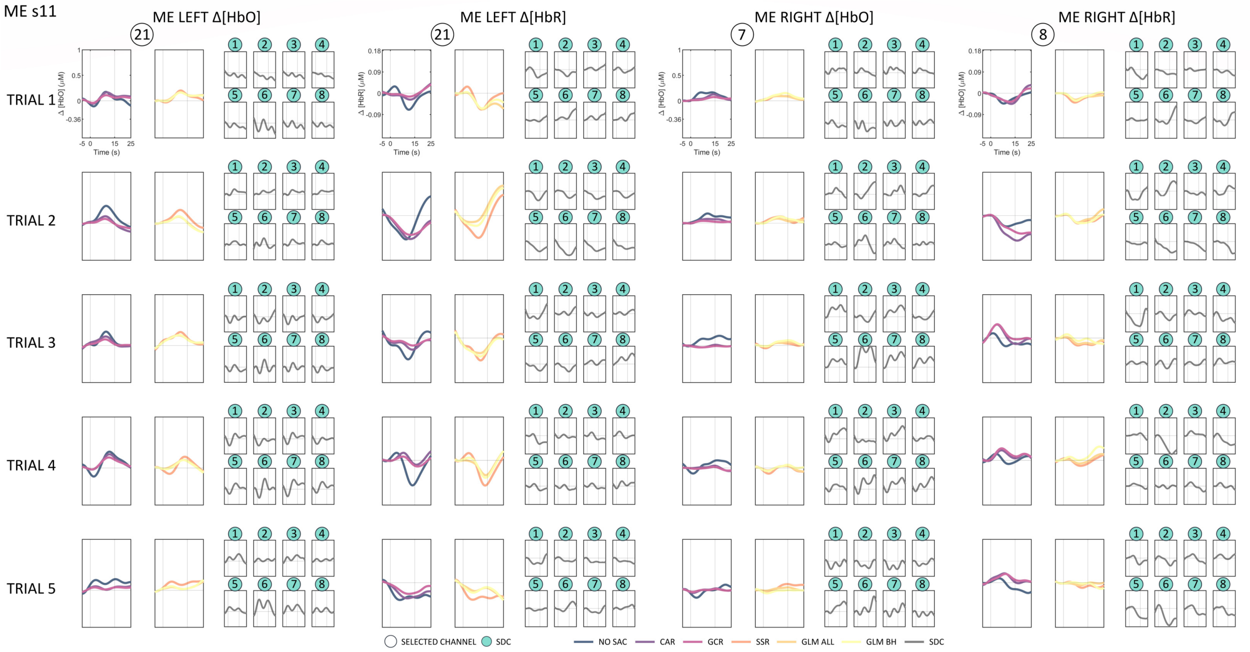
ME s11
ME LEFT Δ[HbO]
ME LEFT Δ[HbR]
ME RIGHT Δ[HbO]
ME RIGHT Δ[HbR]
TRIAL 1
TRIAL 2
TRIAL 3
TRIAL 4
TRIAL 5
Time (s)
SELECTED CHANNEL
SDC
NO SAC
CAR
GCR
SSR
GLM ALL
GLM BH
SDC

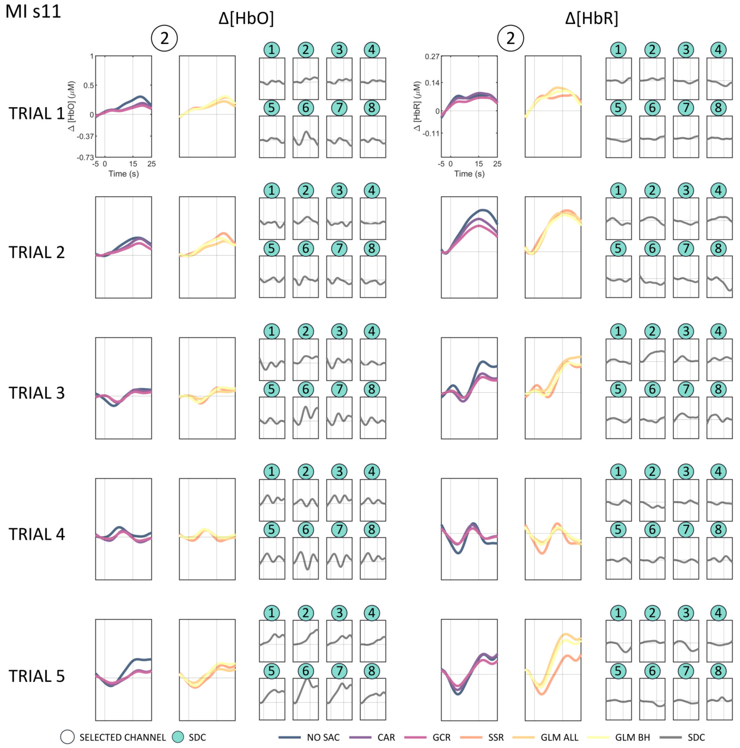
MI s11
Δ[HbO]
Δ[HbR]
TRIAL 1
TRIAL 2
TRIAL 3
TRIAL 4
TRIAL 5
Time (s)
SELECTED CHANNEL
SDC
NO SAC
CAR
GCR
SSR
GLM ALL
GLM BH
SDC

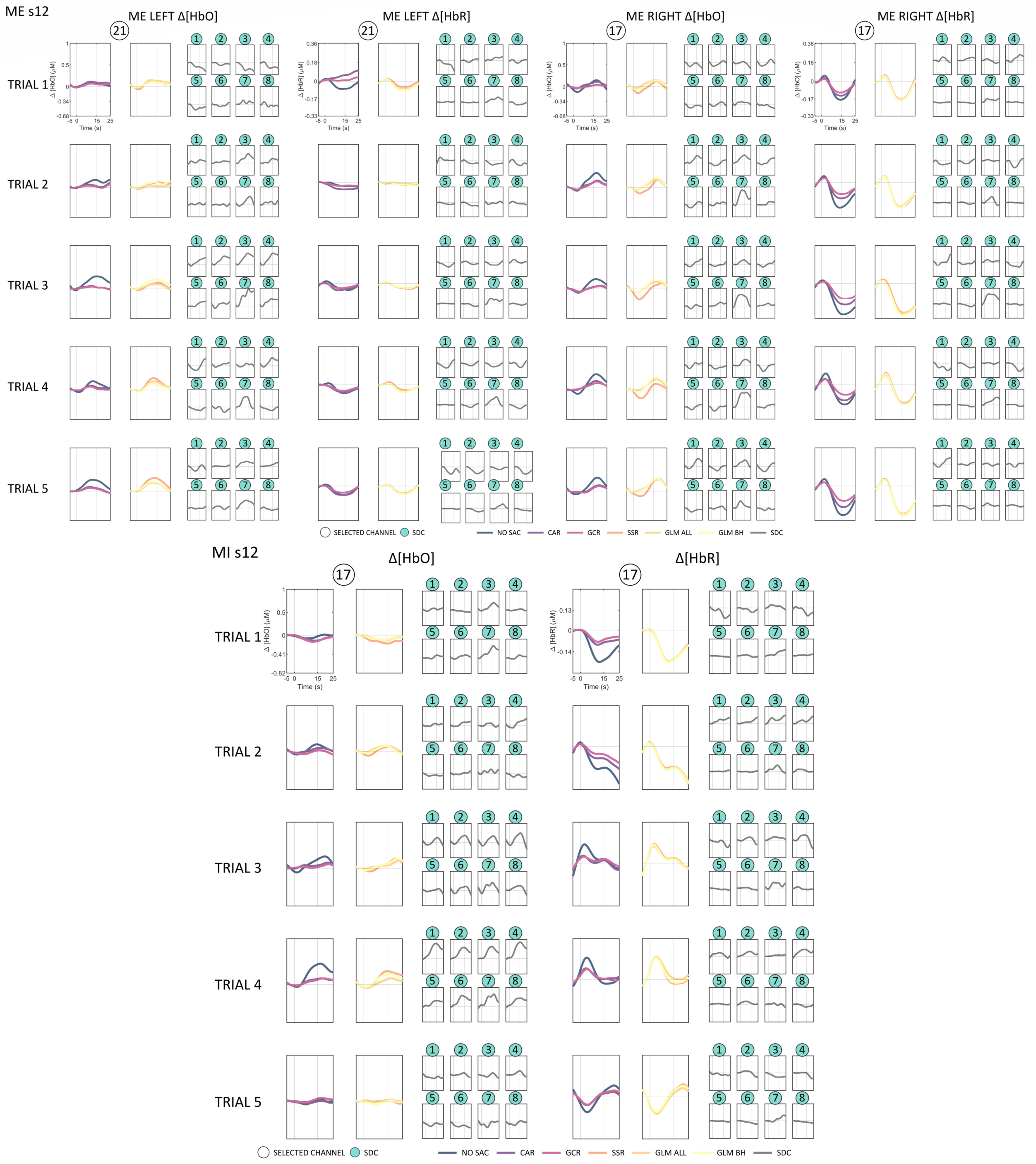
ME s12
ME LEFT Δ[HbO]
ME LEFT Δ[HbR]
ME RIGHT Δ[HbO]
ME RIGHT Δ[HbR]
TRIAL 1
TRIAL 2
TRIAL 3
TRIAL 4
TRIAL 5
SELECTED CHANNEL
SDC
NO SAC
CAR
GCR
SSR
GLM ALL
GLM BH
SDC
MI s12
Δ[HbO]
Δ[HbR]
TRIAL 1
TRIAL 2
TRIAL 3
TRIAL 4
TRIAL 5
SELECTED CHANNEL
SDC
NO SAC
CAR
GCR
SSR
GLM ALL
GLM BH
SDC

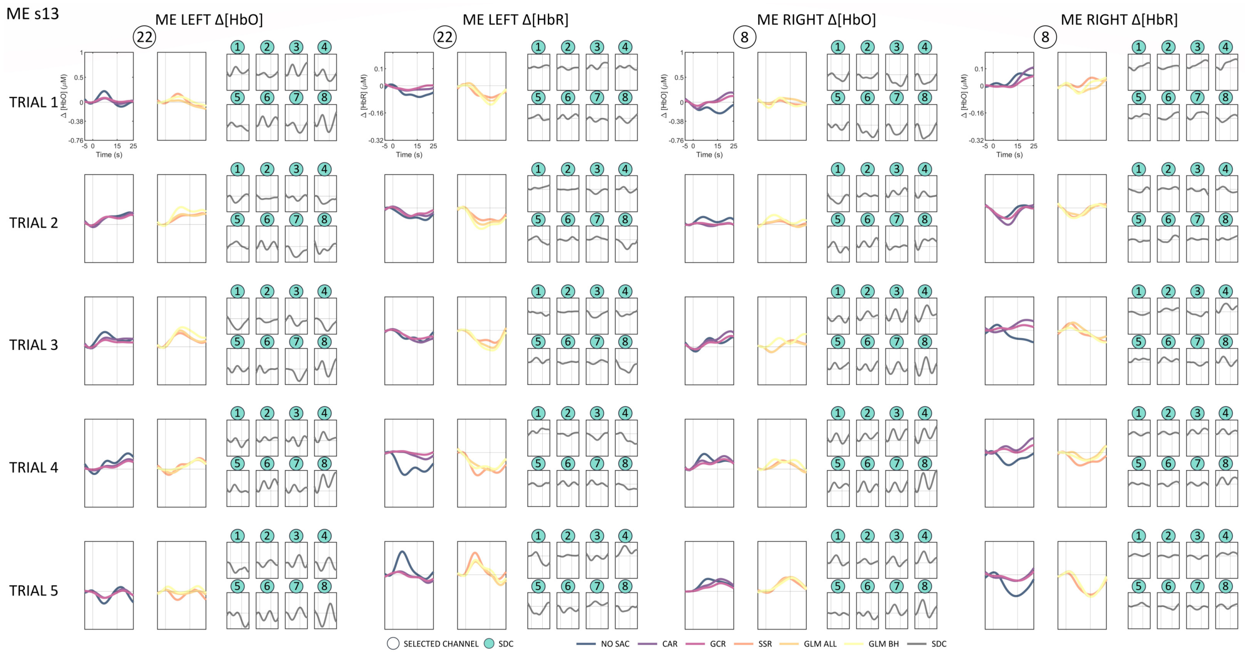
ME s13
ME LEFT Δ[HbO]
ME LEFT Δ[HbR]
ME RIGHT Δ[HbO]
ME RIGHT Δ[HbR]
TRIAL 1
TRIAL 2
TRIAL 3
TRIAL 4
TRIAL 5
SELECTED CHANNEL
SDC
NO SAC
CAR
GCR
SSR
GLM ALL
GLM BH
SDC

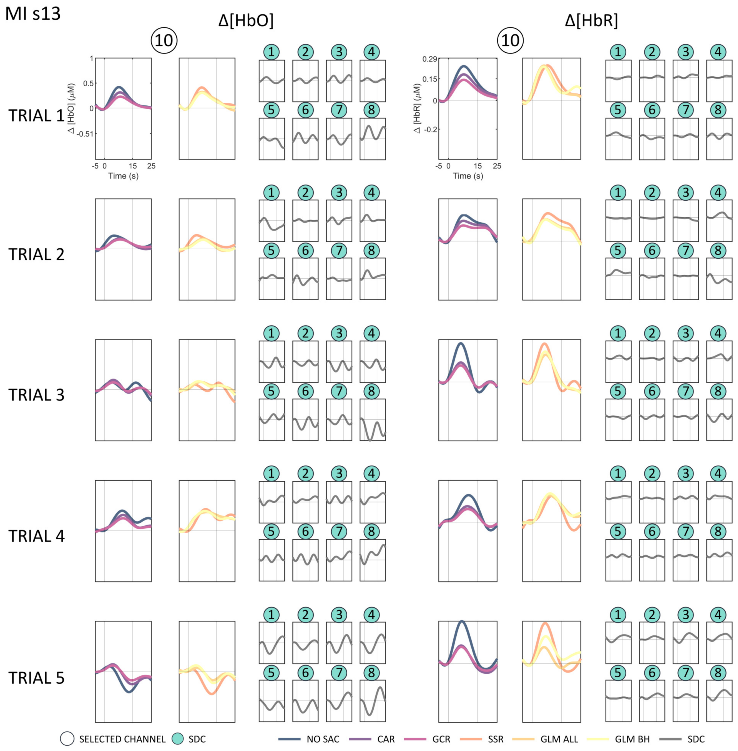
MI s13
Δ[HbO]
Δ[HbR]
TRIAL 1
TRIAL 2
TRIAL 3
TRIAL 4
TRIAL 5
SELECTED CHANNEL
SDC
NO SAC
CAR
GCR
SSR
GLM ALL
GLM BH
SDC

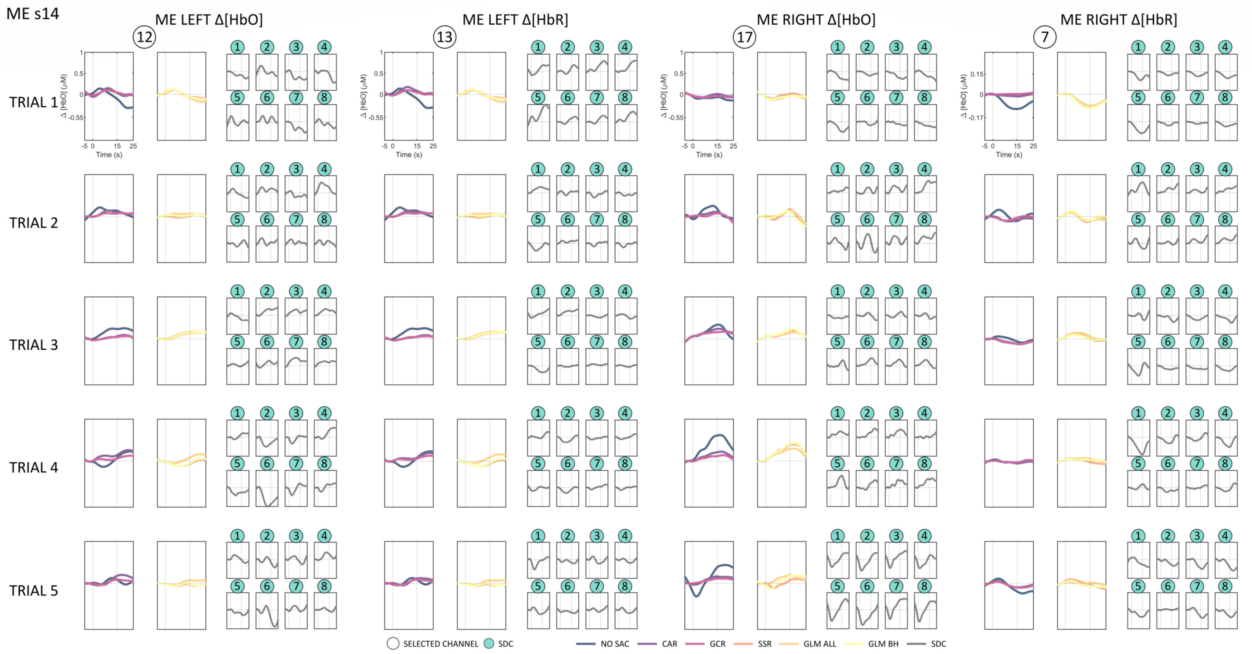
ME s14
ME LEFT Δ[HbO]
ME LEFT Δ[HbR]
ME RIGHT Δ[HbO]
ME RIGHT Δ[HbR]
TRIAL 1
TRIAL 2
TRIAL 3
TRIAL 4
TRIAL 5
Δ [HbO] (μM)
Time (s)
SELECTED CHANNEL
SDC
NO SAC
CAR
GCR
SSR
GLM ALL
GLM BH
SDC

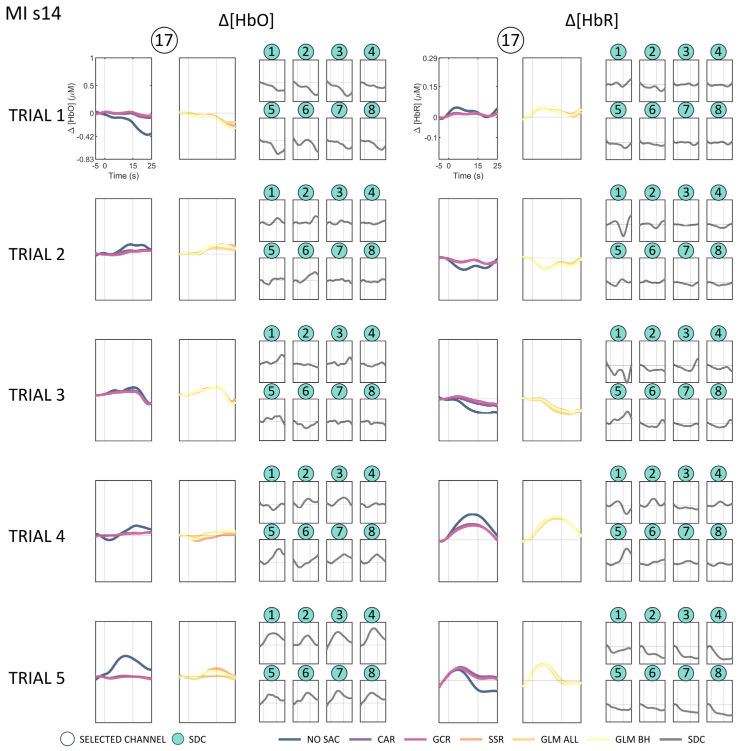
MI s14
Δ[HbO]
Δ[HbR]
TRIAL 1
TRIAL 2
TRIAL 3
TRIAL 4
TRIAL 5
Δ [HbO] (μM)
Δ [HbR] (μM)
Time (s)
SELECTED CHANNEL
SDC
NO SAC
CAR
GCR
SSR
GLM ALL
GLM BH
SDC

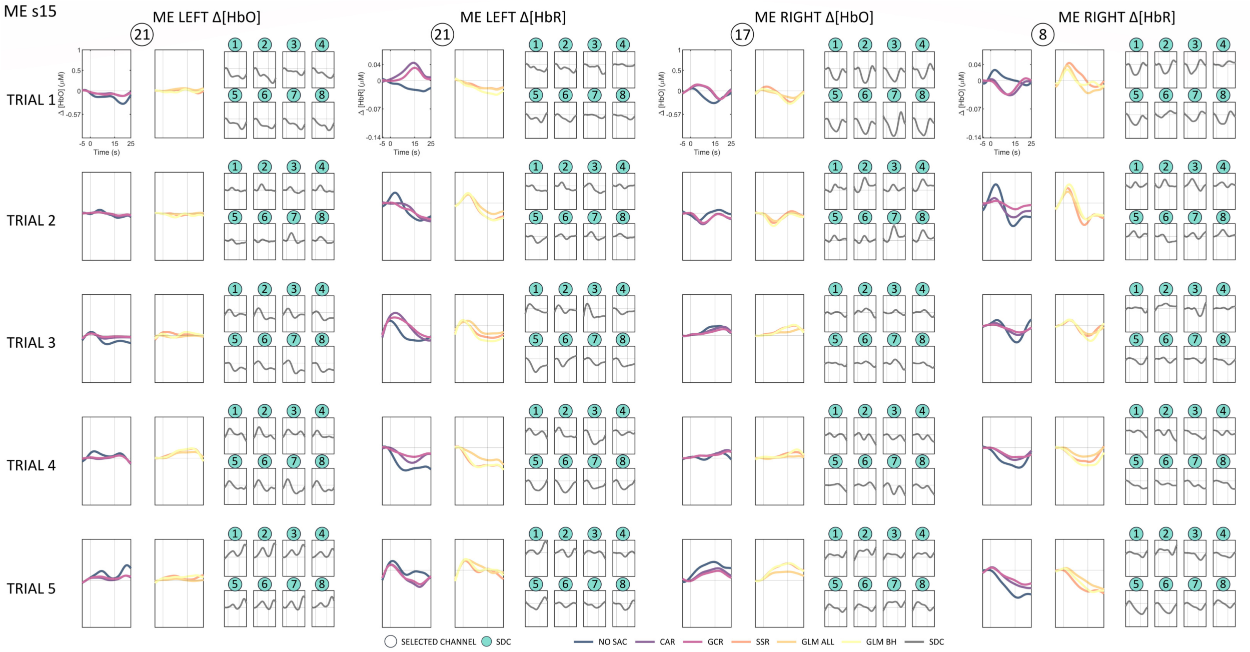
ME s15
ME LEFT Δ[HbO]
ME LEFT Δ[HbR]
ME RIGHT Δ[HbO]
ME RIGHT Δ[HbR]
TRIAL 1
TRIAL 2
TRIAL 3
TRIAL 4
TRIAL 5
Time (s)
SELECTED CHANNEL
SDC
NO SAC
CAR
GCR
SSR
GLM ALL
GLM BH
SDC

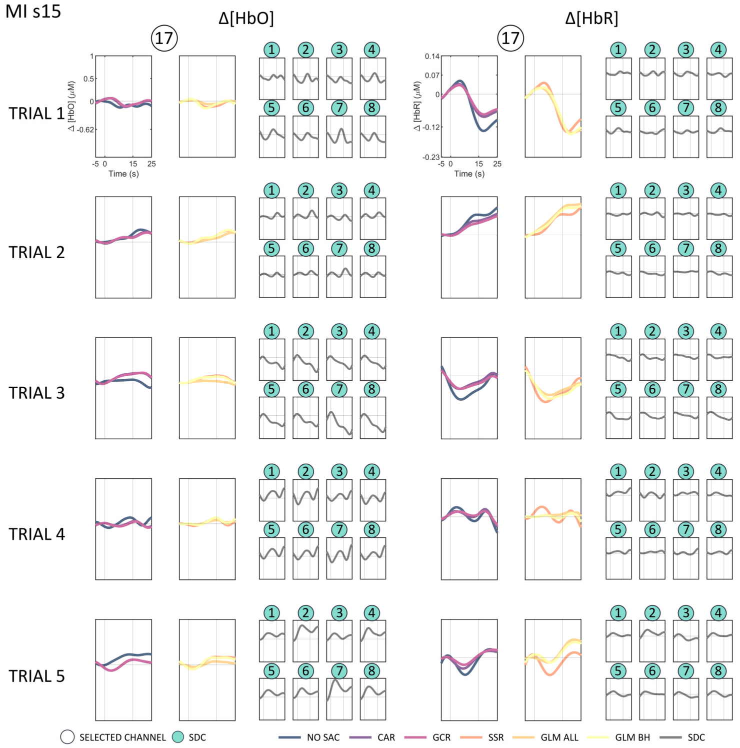
MI s15
Δ[HbO]
Δ[HbR]
TRIAL 1
TRIAL 2
TRIAL 3
TRIAL 4
TRIAL 5
Time (s)
SELECTED CHANNEL
SDC
NO SAC
CAR
GCR
SSR
GLM ALL
GLM BH
SDC

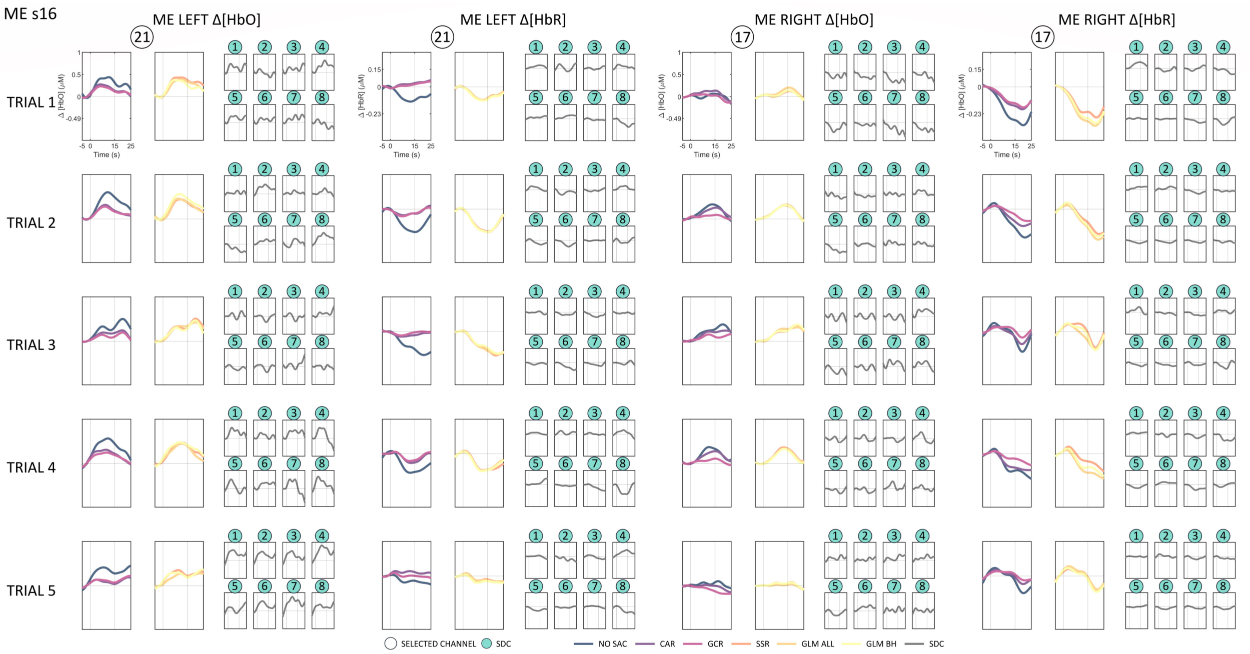
ME s16
ME LEFT Δ[HbO]
ME LEFT Δ[HbR]
ME RIGHT Δ[HbO]
ME RIGHT Δ[HbR]
TRIAL 1
TRIAL 2
TRIAL 3
TRIAL 4
TRIAL 5
Time (s)
SELECTED CHANNEL
SDC
NO SAC
CAR
GCR
SSR
GLM ALL
GLM BH
SDC

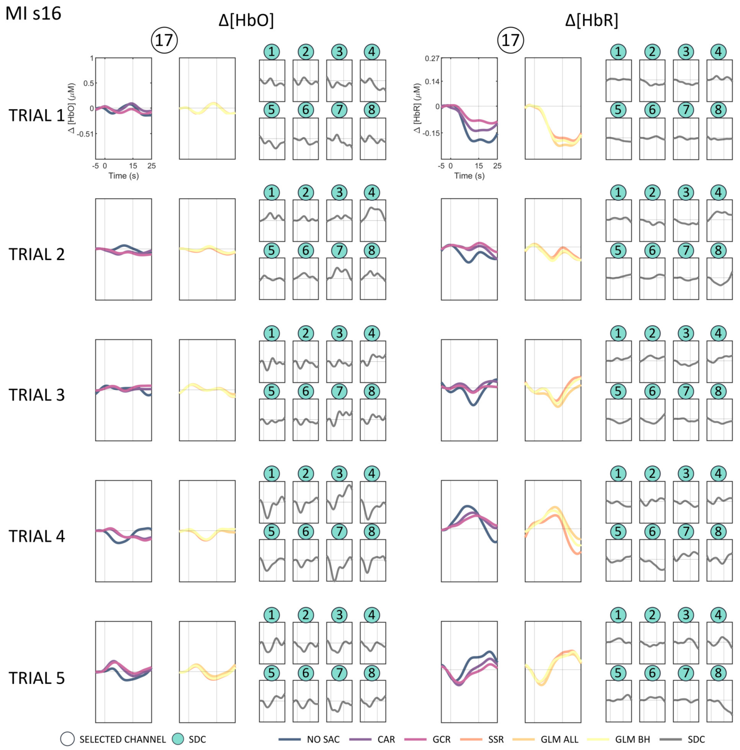
MI s16
Δ[HbO]
Δ[HbR]
TRIAL 1
TRIAL 2
TRIAL 3
TRIAL 4
TRIAL 5
Time (s)
SELECTED CHANNEL
SDC
NO SAC
CAR
GCR
SSR
GLM ALL
GLM BH
SDC

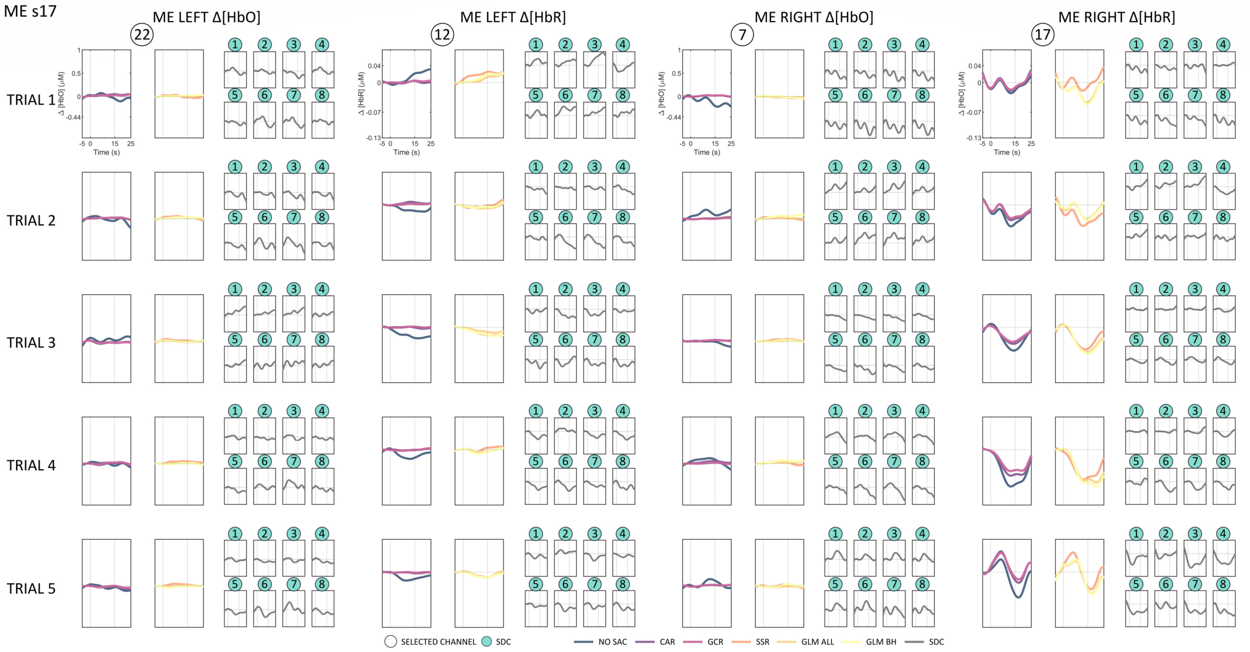
ME s17
ME LEFT Δ[HbO]
ME LEFT Δ[HbR]
ME RIGHT Δ[HbO]
ME RIGHT Δ[HbR]
TRIAL 1
TRIAL 2
TRIAL 3
TRIAL 4
TRIAL 5
SELECTED CHANNEL
SDC
NO SAC
CAR
GCR
SSR
GLM ALL
GLM BH
SDC

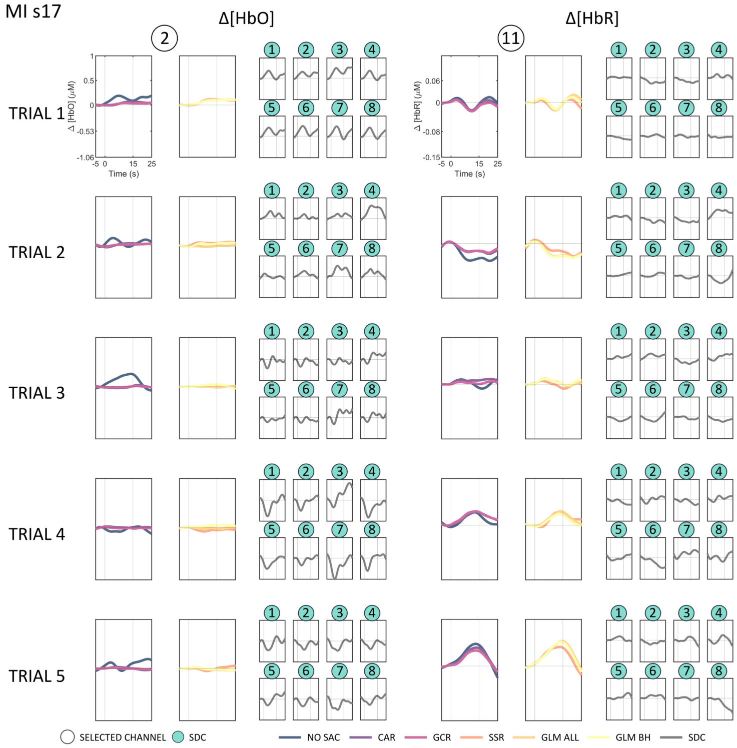
MI s17
Δ[HbO]
Δ[HbR]
TRIAL 1
TRIAL 2
TRIAL 3
TRIAL 4
TRIAL 5
SELECTED CHANNEL
SDC
NO SAC
CAR
GCR
SSR
GLM ALL
GLM BH
SDC

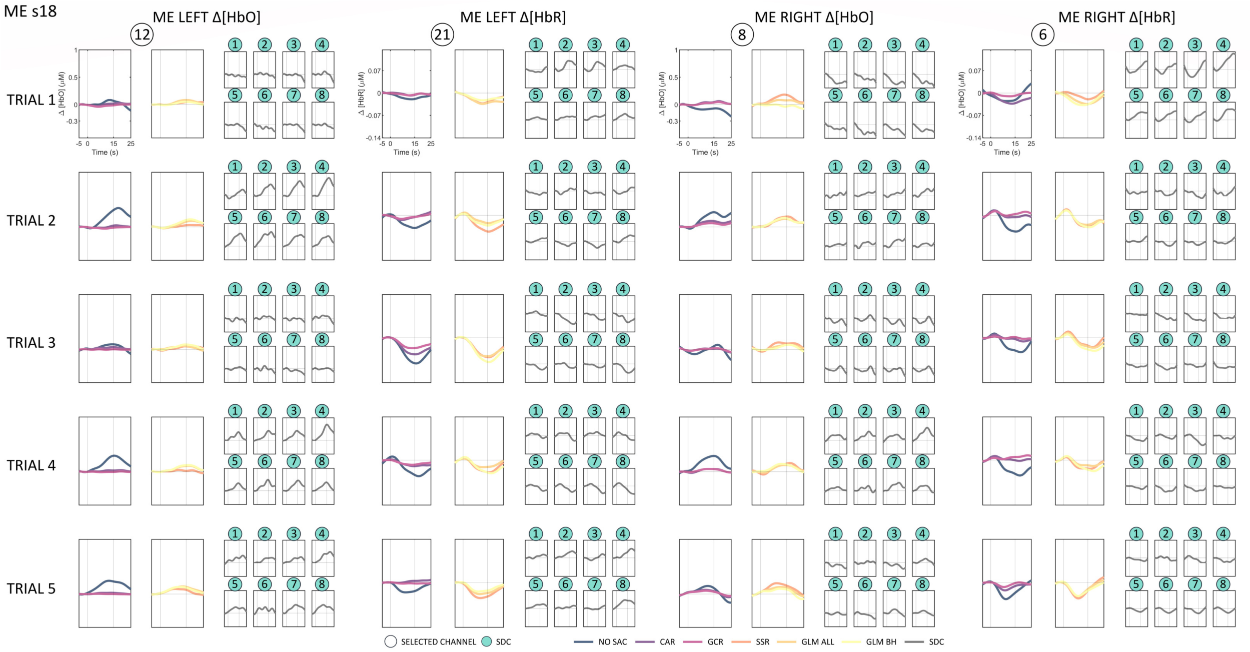
ME s18
ME LEFT Δ[HbO]
ME LEFT Δ[HbR]
ME RIGHT Δ[HbO]
ME RIGHT Δ[HbR]
TRIAL 1
TRIAL 2
TRIAL 3
TRIAL 4
TRIAL 5
Time (s)
SELECTED CHANNEL
SDC
NO SAC
CAR
GCR
SSR
GLM ALL
GLM BH
SDC

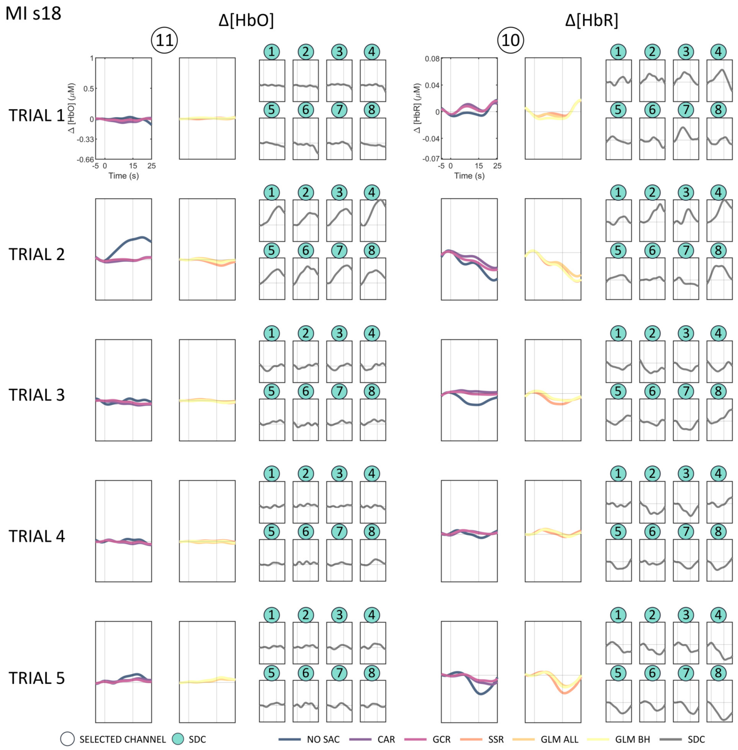
MI s18
Δ[HbO]
Δ[HbR]
TRIAL 1
TRIAL 2
TRIAL 3
TRIAL 4
TRIAL 5
Time (s)
SELECTED CHANNEL
SDC
NO SAC
CAR
GCR
SSR
GLM ALL
GLM BH
SDC

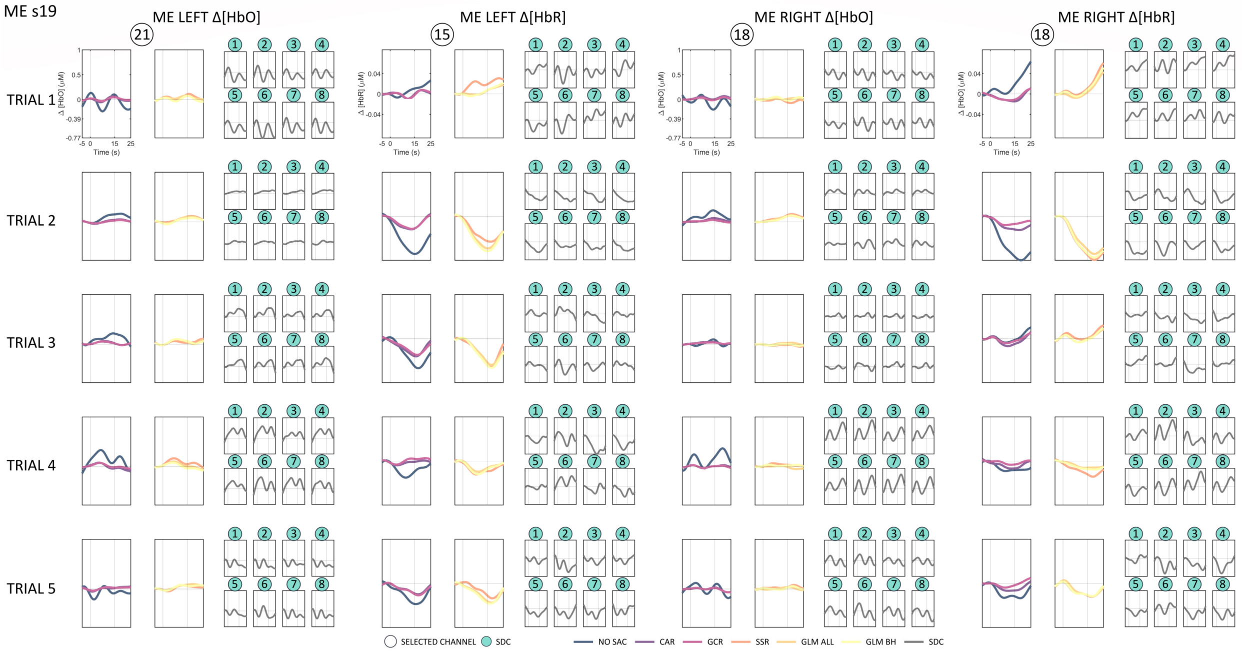
ME s19
ME LEFT Δ[HbO]
ME LEFT Δ[HbR]
ME RIGHT Δ[HbO]
ME RIGHT Δ[HbR]
21
15
18
18
TRIAL 1
TRIAL 2
TRIAL 3
TRIAL 4
TRIAL 5
Δ [HbO] (μM)
Δ [HbR] (μM)
Time (s)
SELECTED CHANNEL
SDC
NO SAC
CAR
GCR
SSR
GLM ALL
GLM BH
SDC

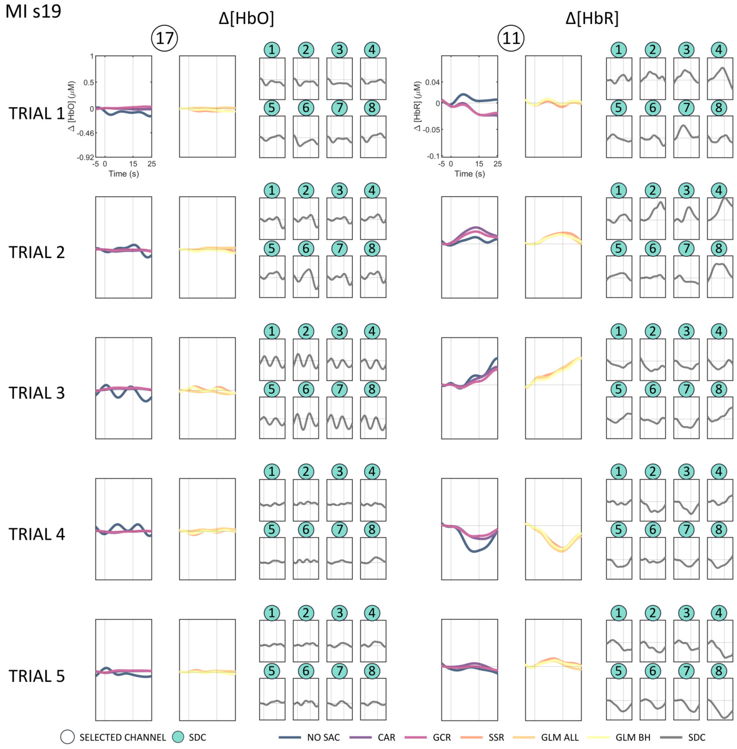
MI s19
Δ[HbO]
Δ[HbR]
17
11
TRIAL 1
TRIAL 2
TRIAL 3
TRIAL 4
TRIAL 5
Δ [HbO] (μM)
Δ [HbR] (μM)
Time (s)
SELECTED CHANNEL
SDC
NO SAC
CAR
GCR
SSR
GLM ALL
GLM BH
SDC

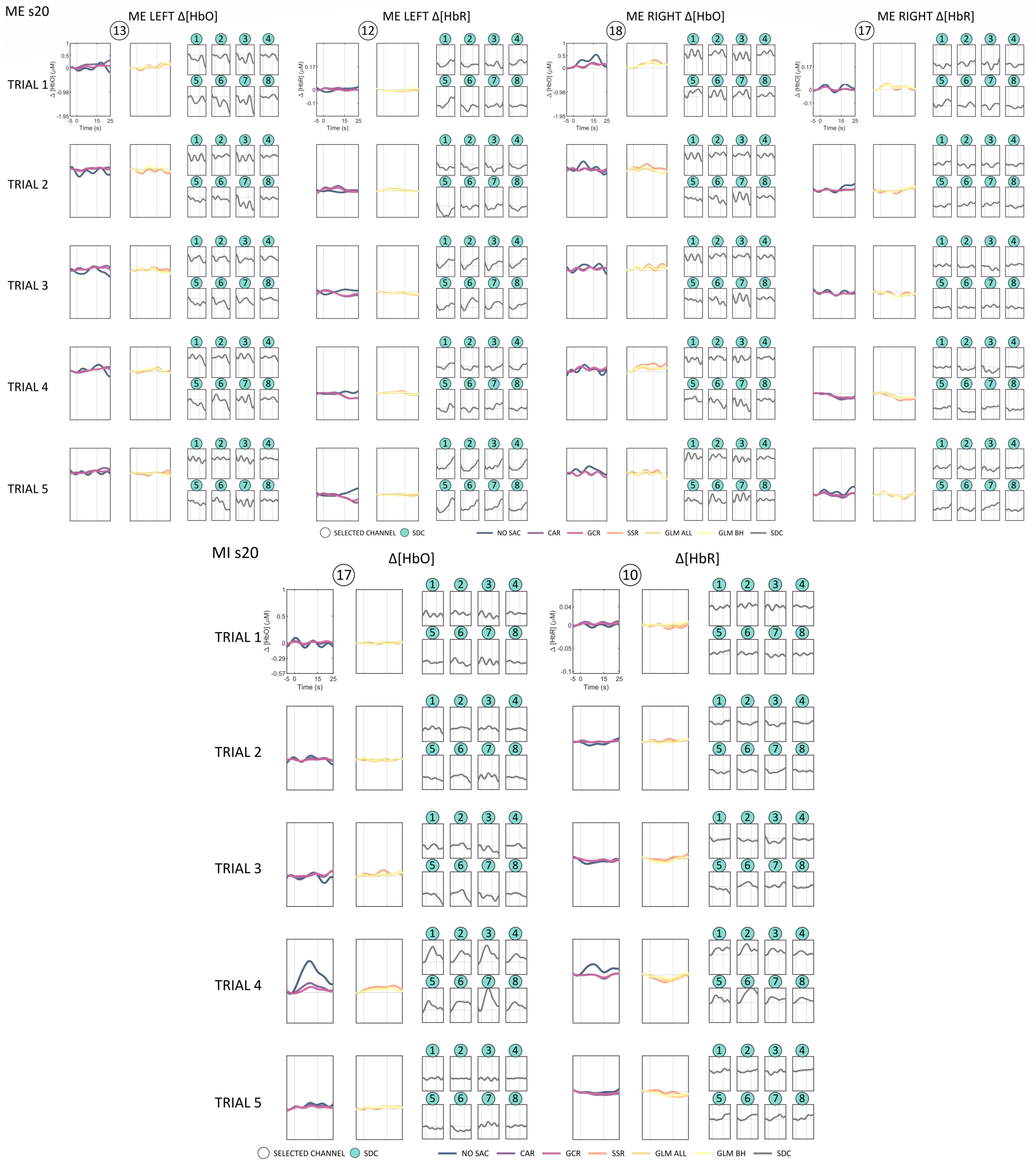
ME s20
ME LEFT Δ[HbO]
ME LEFT Δ[HbR]
ME RIGHT Δ[HbO]
ME RIGHT Δ[HbR]
TRIAL 1
TRIAL 2
TRIAL 3
TRIAL 4
TRIAL 5
Time (s)
SELECTED CHANNEL
SDC
NO SAC
CAR
GCR
SSR
GLM ALL
GLM BH
SDC
MI s20
Δ[HbO]
Δ[HbR]
TRIAL 1
TRIAL 2
TRIAL 3
TRIAL 4
TRIAL 5
Time (s)
SELECTED CHANNEL
SDC
NO SAC
CAR
GCR
SSR
GLM ALL
GLM BH
SDC

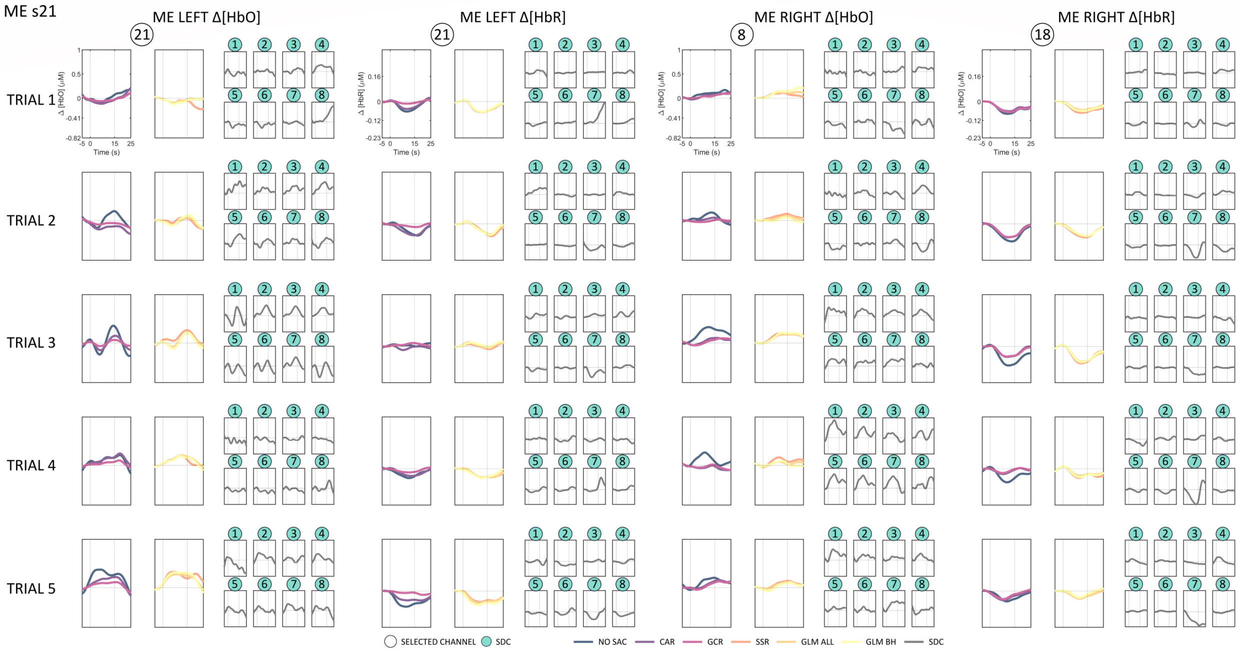
ME s21
ME LEFT Δ[HbO]
ME LEFT Δ[HbR]
ME RIGHT Δ[HbO]
ME RIGHT Δ[HbR]
TRIAL 1
TRIAL 2
TRIAL 3
TRIAL 4
TRIAL 5
Time (s)
SELECTED CHANNEL
SDC
NO SAC
CAR
GCR
SSR
GLM ALL
GLM BH
SDC

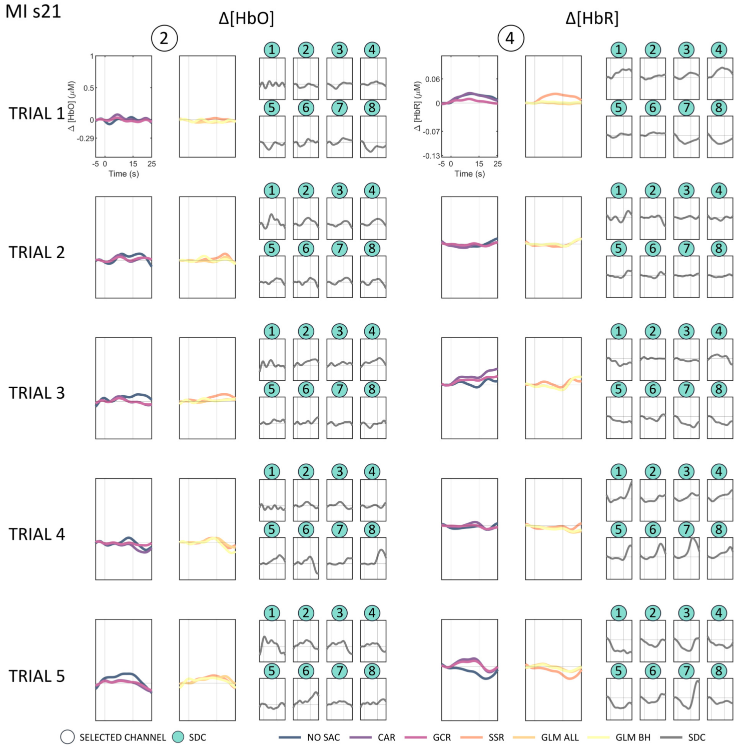
MI s21
Δ[HbO]
Δ[HbR]
TRIAL 1
TRIAL 2
TRIAL 3
TRIAL 4
TRIAL 5
Time (s)
SELECTED CHANNEL
SDC
NO SAC
CAR
GCR
SSR
GLM ALL
GLM BH
SDC

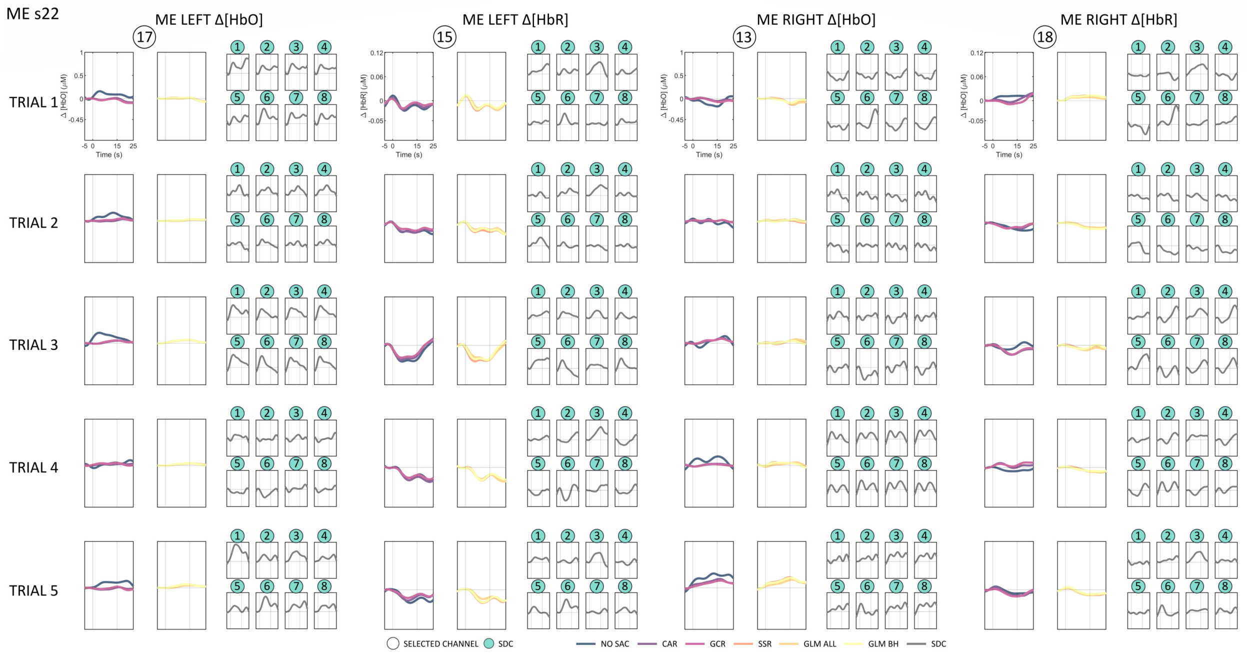
ME s22
ME LEFT Δ[HbO]
ME LEFT Δ[HbR]
ME RIGHT Δ[HbO]
ME RIGHT Δ[HbR]
TRIAL 1
TRIAL 2
TRIAL 3
TRIAL 4
TRIAL 5
Time (s)
SELECTED CHANNEL
SDC
NO SAC
CAR
GCR
SSR
GLM ALL
GLM BH
SDC

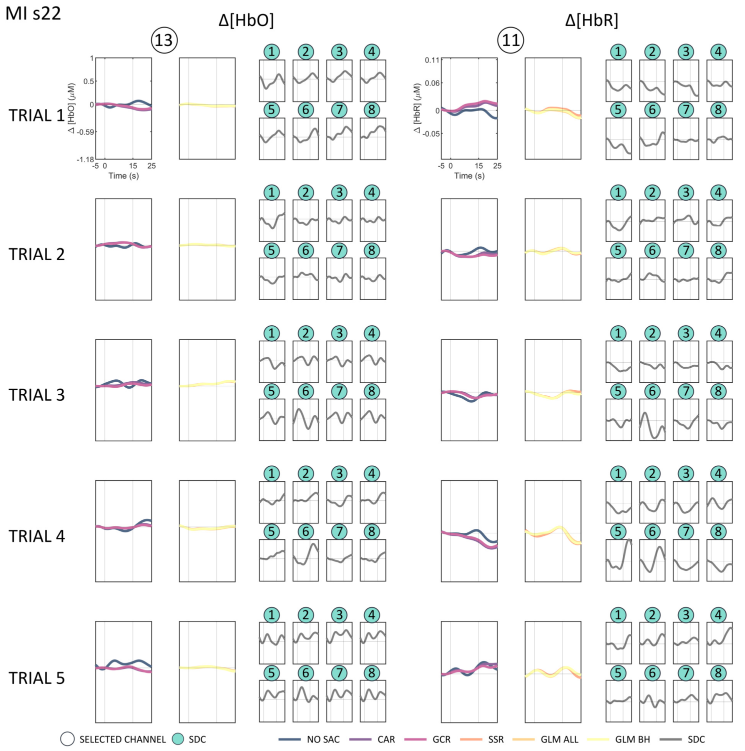
MI s22
Δ[HbO]
Δ[HbR]
TRIAL 1
TRIAL 2
TRIAL 3
TRIAL 4
TRIAL 5
Time (s)
SELECTED CHANNEL
SDC
NO SAC
CAR
GCR
SSR
GLM ALL
GLM BH
SDC

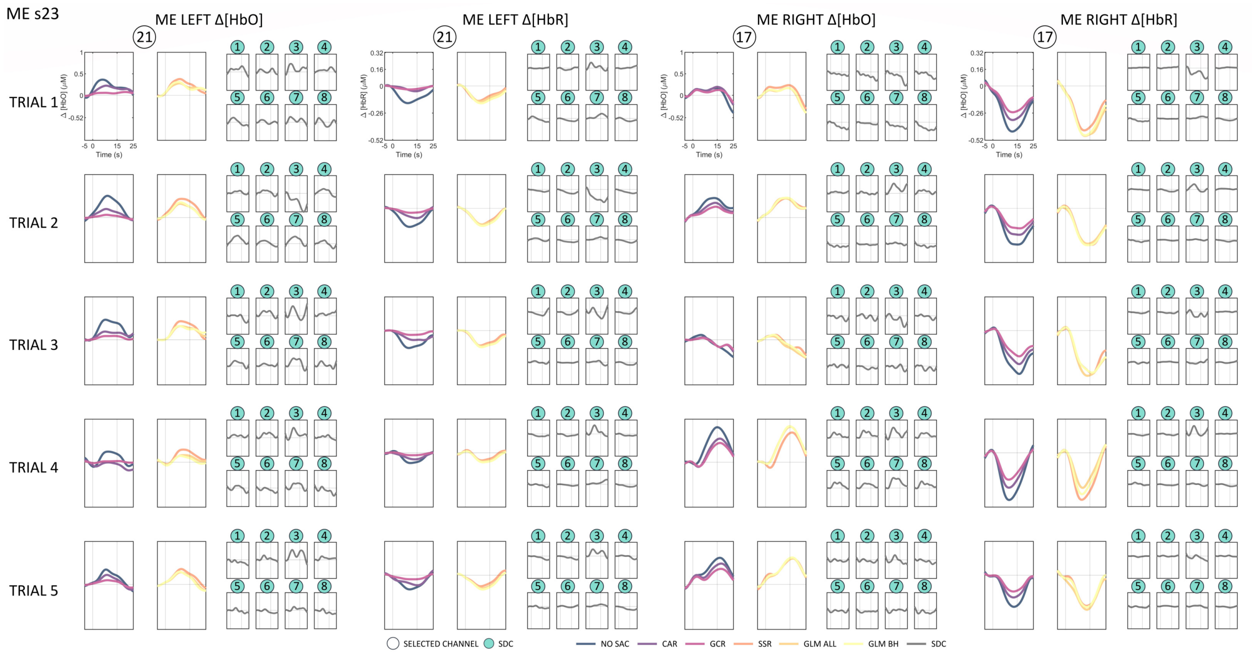
ME s23
ME LEFT Δ[HbO]
ME LEFT Δ[HbR]
ME RIGHT Δ[HbO]
ME RIGHT Δ[HbR]
TRIAL 1
TRIAL 2
TRIAL 3
TRIAL 4
TRIAL 5
SELECTED CHANNEL
SDC
NO SAC
CAR
GCR
SSR
GLM ALL
GLM BH
SDC

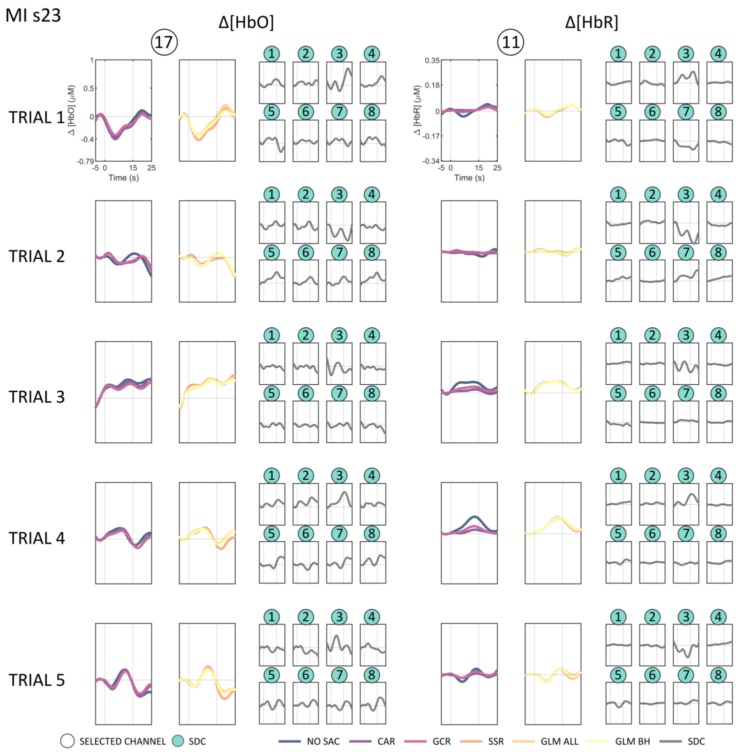
MI s23
Δ[HbO]
Δ[HbR]
TRIAL 1
TRIAL 2
TRIAL 3
TRIAL 4
TRIAL 5
SELECTED CHANNEL
SDC
NO SAC
CAR
GCR
SSR
GLM ALL
GLM BH
SDC

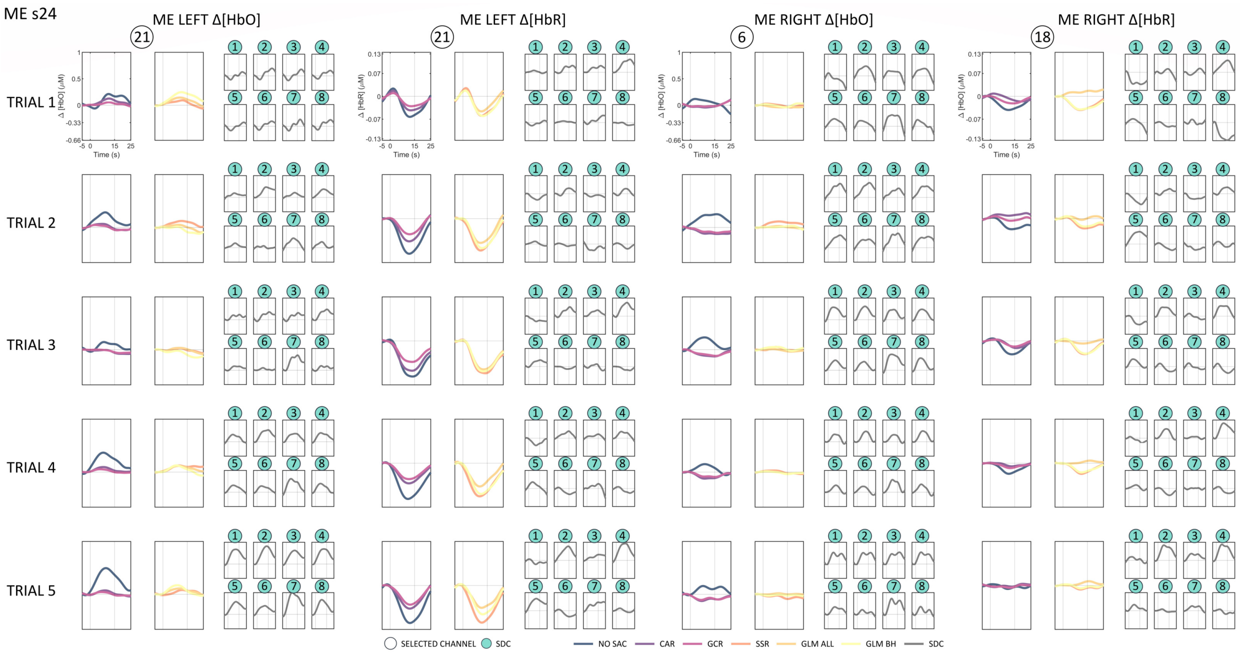
ME s24
ME LEFT Δ[HbO]
ME LEFT Δ[HbR]
ME RIGHT Δ[HbO]
ME RIGHT Δ[HbR]
TRIAL 1
TRIAL 2
TRIAL 3
TRIAL 4
TRIAL 5
Time (s)
SELECTED CHANNEL
SDC
NO SAC
CAR
GCR
SSR
GLM ALL
GLM BH
SDC

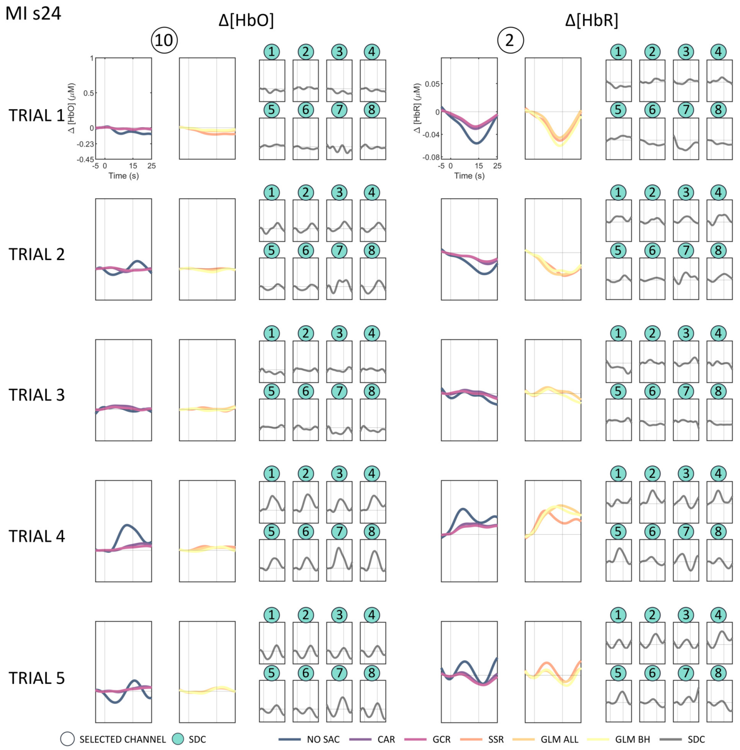
MI s24
Δ[HbO]
Δ[HbR]
TRIAL 1
TRIAL 2
TRIAL 3
TRIAL 4
TRIAL 5
Time (s)
SELECTED CHANNEL
SDC
NO SAC
CAR
GCR
SSR
GLM ALL
GLM BH
SDC
